# A Sexual Bias in mitochondrial protein-coding gene expression across different tissues and the prognostic value in multiple cancers

**DOI:** 10.1101/2022.11.22.517535

**Authors:** Alan Tardin da Silva, Cristina dos Santos Ferreira, Enrique Medina-Acosta

## Abstract

Mitochondria in mammalian cells provide ATP through oxidative phosphorylation. The overproduction of reactive oxygen species (ROS) in mitochondrial cells promotes cancer by modifying gene expression or function. Mating introduces competing mitochondrial (mtDNA) and nuclear DNA (nDNA) gene products, leading to biological differences between males and females for diseases and disorders such as cancer. There is a significant sex bias in aging-related conditions. We aimed to investigate whether sex and age affect mitochondrial protein-coding gene expression in cancer and, if so, to determine the prognosis value in survival outcomes, stemness, and immune cell infiltrates. We compared normal versus primary tumor transcriptomes (bulk RNA-Seq) from The Cancer Genome Atlas (TCGA), and the Genotype-Tissue Expression (GTEx) projects to test these hypotheses. Correlations between gene expression, survival, protective or risk factor, stemness, and immune cell infiltrate were performed in RStudio using UCSC Xena Shiny. Eleven mitochondrial protein-coding genes were altered in brain cancer (*MT-ND2*, *MT-ND1*, *MT-ATP8*, *MT-ATP6*, *MT-CO2*, *MT-CYB*, *MT-CO3*, *MT-ND4L*, *MT-ND4*, *MT-ND3*, *MT-CO1*). MT-ND5 and MT-ND6 are disproportionately expressed in female brain tissues. Mitochondrial global polymorphic expression sites of variation were more significant in the 50-59 and 60-79-year-old age groups than in the 20-49-year-old age groups. Pan-cancer survival analysis revealed a 4-component gene signature (*MT-CO1*, *MT-CO2*, *MT-ND5*, *and MT-ND6*) downregulated in low-grade glioma (LGG). This gene signature increased LGG overall survival, disease-specific survival, and progression-free interval without sex-specific association. However, the correlation with disease-free interval survival was female-specific. The 4-component gene signature was protective in LGG but risky in thymoma cancer and uterine corpus endometrial carcinoma. In LGG, the 4-component gene signature positively correlated with immune monocyte, NK, and B cell infiltrates and negatively correlated with T cell CD4+ Th2, macrophage M1 and M2, myeloid dendritic cell, and neutrophil. We identified a 13-component mitochondrial protein-coding gene signature associated with stemness in kidney chromophobe. A sex-biased effect was observed in mitochondrial protein-coding for brain tissues, with a female bias. However, an aging effect with higher polymorphic site expression was observed in male tissues. We conclude that the differentially expressed mitochondrial protein-coding genes provide new insights into carcinogenesis, helping to identify new prognostic markers. The overexpression of the 4-component gene signature is associated with a better prognosis in LGG, with positive and negative correlations with immune cell infiltrates.

## INTRODUCTION

The mammalian mitochondrial DNA (mtDNA) encodes 13 key proteins essential for the functioning of the oxidative phosphorylation system (OXPHOS) (Larsson NG, 1998;Foster et al., 2006;Gustafsson et al., 2016). The mitochondria are therefore considered a vital component of all cells with a nucleus, so mtDNA-associated disorders or diseases can affect many tissues, and clinical features are variable. Moreover, defects in mitochondrial metabolism cause a wide range of human diseases, such as cancer, which include examples from all medical fields. Thus, mtDNA disorders are associated with a range of multi-systemic diseases in humans, where the understanding of the spectrum of these diseases has been expanded by the recognition of mutations in protein-coding genes in mitochondria caused by reactive oxygen species (ROS), which cause not only several neurodegenerative diseases but also other disorders as cancer (Harman, 1992b;Wallace, 1992;Geromel et al., 2001;Taylor and Turnbull, 2005;Schapira, 2006). Studies in the last ten years have shown that the human mtDNA of 13 protein-coding acts in several more functions than was described (Faure et al., 2011;Lee et al., 2013;Breton et al., 2014;Capt et al., 2016). Thus, investigating these genes’ expression using RNA-Seq experiments is very useful in understanding the dynamics of expression in a population. It is also known that an RNA-Seq analysis allows the identification of gene expression signatures, which can be altered by external factors such as the environment, health status, or disease (da Silva Francisco Junior et al., 2019).

Using RNA-Seq analysis to investigate the extent of allelic specific expression (ASE) becomes a challenging task for mitochondrial protein-coding genes, because mitochondria have a haploid genome, and it is not possible to perform a genome-wide ASE, only for the detection of microheteroplasmy. Since microheteroplasmy is a form of heteroplasmy, this would also damage the mtDNA, but the levels of microheteroplasmy are about 2−5% in mitochondrial genomes. Moreover, the microheteroplasmy and heteroplasmy are present in mitochondrial diseases but also in normal individuals, where heteroplasmic variants without apparent functional consequences are observed in samples from individuals without any overt mitochondrial disease (Calloway et al., 2000;Kirches et al., 2001;Santos et al., 2008;He et al., 2010). Furthermore, there is a variation in pattern expression between mtDNA and the nuclear DNA variants. Consequently, calculating the ASE for mitochondrial protein-coding genes involves using a different methodology to calculate this specific expression in a population (Gemmell et al., 2004;Kassam et al., 2016;da Silva Francisco Junior et al., 2019;Stenton and Prokisch, 2020;Stewart and Chinnery, 2021).

The mtDNA is in the mitochondrial matrix and is closed to the respiratory chains, which are the primary source of ROS from the OXPHOS system. ROS induce somatic mutations in mtDNA due to the lack of protection by histones (Richter, 1995;Shokolenko et al., 2009). Thus, cumulative mutations in mtDNA can result in dysfunction of the OXPHOS system, leading to diseases associated with mitochondria, which is a hallmark of many diseases, such as cancer (van Oven and Kayser, 2009;Srinivasan et al., 2017). Additionally, the mitochondria are both producers and targets of oxidative stress induced by exposure to ROS, resulting in mitochondrial dysfunction and interfering in electron transfer activity in oxidative phosphorylation. Defects in electron transfer activity increase ROS production, thus establishing a “vicious cycle” with aging that interferes with production and target properties (Miquel et al., 1980;Linnane AW, 1989;Papa, 1996;Ozawa, 1997).

Aging is a universal process whose manifestations are familiar and unmistakable, and old age in humans and even animals can be readily recognized after a minimal assessment. Despite this, an accepted definition of aging and a detailed understanding of the biological mechanisms underpinning aging are elusive. So, aging is described as the progressive loss of function accompanied by decreased fertility and increased mortality and disability (Kirkwood, 2000). Thus, aging is associated with evidence of deleterious changes in the molecular structure of DNA, proteins, lipids, and prostaglandins, all markers of oxidative stress (Harman, 1992a;1993). It has been recognized that ROS also play a role in normal signaling processes and that their generation is essential for maintaining homeostasis and cellular responsiveness (Droge, 2002). Since mitochondria are involved in oxidative stress, a mitochondrial theory of aging has been proposed, where an accumulation of somatic mutations in mtDNA induced by exposure to ROS leads to errors in the mtDNA-encoded polypeptides, affecting the electron transfer and oxidative phosphorylation system (Miquel et al., 1980;Linnane AW, 1989). Declines in the activity of the mitochondrial respiratory system and its constituent enzymes have been reported with advancing age, notably in cytochrome c oxidase, in several tissues, including skeletal muscle, heart, and liver (Muller-Hocker, 1989). Thus, the integrity of mitochondrial DNA in these tissues is gradually reduced with age, evidenced by the accumulation of deletions, duplications, and some point mutations in the mtDNA (Nagley and Wei, 1998;Kopsidas et al., 2002).

Having this perspective on the mammalian mtDNA, which encodes 13 proteins essential for the functioning of other systems, like the OXPHOS system, we considered building a computational meta-analysis of the mitochondrial protein-coding genes to investigate the existence of differentially expressed genes (DEG) in healthy tissues versus cancer tissues. We also consider exploring differences in the expression of these 13 genes in different tissues, at different age groups, and between sexes (females and males). In the same way, a possible punctual polymorphic point of variable expression of these genes was also investigated. This study approach explores the possibility of sex bias in the expression of the mitochondrial protein-coding gene expression and the impact on survival outcomes in several cancers. Thus, we hypothesize that a sex-bias expression of the 13 mitochondrial protein-coding genes may exist across different tissues and in the survival outcomes. Additionally, we hypothesize that the polymorphic site expression in mitochondria protein-coding genes may differ between the sexes at different age groups impacting pathologies such as cancer.

## MATERIAL AND METHODS

### Analysis of Differential Expression of Mitochondrial Protein-Coding Genes in Different Tissues

We used 19,131 RNA-Seq experiments samples from 18.972 donors (8516 Female and 10456 Male) from The Genotype-Tissue Expression (GTEx) for 52 non-diseased tissue sites across nearly 1000 individuals and The Cancer Genome Atlas (TCGA) for over 20,000 primary cancer and matched samples from healthy subjects spanning 33 cancer types, to search for a differential expressed genes (DEG). We used the UCSC Xena browser (https://xena.ucsc.edu/) to construct the base of this analysis and selected the TCGA TARGET GTEx cohort study and the phenotypic variables primary_site, study, sample type, main category, and gender as our first variable to organize our data based on these features. Posteriorly, we selected our second variable, genomic, using the mitochondrial protein-coding genes as input. Here, we used a gene expression RNA-Seq - RSEM norm_count, to access the expression of the 13 mitochondrial protein-coding genes. The RSEM was used because it relates to gene abundances from single-end or paired-end RNA-Seq data; therefore, RSEM outputs abundance estimates, 95% credibility intervals, and visualization files and can also simulate RNA-Seq data. After, we analyzed the DEG sets (above 1.5 and below -1.5 log2Fold Change, FDR < 0.05) of each primary health tissue compared to the same tissue in cancer (the primary tumor) to search if there is a tissue with DEG in health samples against tumor samples. We used the pipeline from the Ma’ayan lab’s Appyter bulk RNA-Seq analysis and included the L1000FWD analysis, Limma-Voom Differential Gene Expression. All this analyzed data was downloaded from the UCSC Xena browser and taken to RStudio to organize the data.

### Selection of Mitochondrial Protein-Coding Genes and Polymorphic sites

The mitochondrial protein-coding genes used in this analysis were those already described in the literature and validated by MITOMAP, a human mitochondrial genome database used as a reference for mitochondrial analysis (Table S1), and by NCBI genes. Data collection was performed through the RStudio interface, R programming language (https://cran.r-project.org/), using the data packages from the “BiomaRt”, “Bioconductor,” and “BSgenome” libraries to filter the desired information. The analysis involved extracting the physical positions of the 13 mitochondrial protein-coding genes and global polymorphic sites in the population contained within the genes. For this analysis, we used the UCSC Genome Browser of Santa Cruz, a public repository of databases, to construct a custom track with information on the physical positions of genes and polymorphic sites in the mitochondrial genome. The version genome used was the GRCh38/hg38 because it is the only one with data about genes and global polymorphic sites mapped in a population for the mitochondrial genome. We performed a computational meta-analysis to determine the physical position of each mitochondrial protein-coding gene and the global polymorphic sites (in bp). After, we search for polymorphic sites in UCSC genome browser Data base SNPs (dbSNPs151) to construct a site-specific expression based on polymorphic sites of variation already known by the dbSNPs151. This data was downloaded, and this information was integrated into the RStudio environment.

### Expression Analysis of Mitochondrial Protein-Coding Genes and Polymorphic sites

For the analysis of polymorphic site expression at the mitochondrial protein-coding genes, a data collection was built based on the physical positions of the mitochondrial genome, using samples from public repositories of databases, such as UCSC (https://genome.ucsc.edu/). UCSC is an online genome browser with an extensive collection of assemblies and annotations of data about various model organisms, along with a large set of tools for visualizing, analyzing, and downloading data. In this Browser, we work with the GRCh38/hg38 genome version. We accessed the clade option, for Mammal and the genome option, for Human. We selected the group option to access GTEx RNA-Seq Signal Hub, where we extract normalized transcript expression levels (RPKM - Reads Per Kilobase of transcript, per Million mapped reads). Using the RPKM levels, we perform an intersection of these expression data with a custom track, previously obtained, containing the physical position of the polymorphic sites and the 13 mitochondrial protein-coding genes. Lastly, using the table option, we cross all this information with each patient’s sample available in all 52 tissues (Table S2), going to the option output format where data points were selected. The Signal hub data were also separated according to other criteria, such as age groups, in 20-49 years, 50-59 years, and 60-79 years, by sex, male and female.

After obtaining this data collection, some packages were used in RStudio, such as stringr, plyr, rtracklayer, BSgenome, and XML, to perform another computational meta-analysis filtering the mitochondrial protein-coding genes polymorphic sites. These points were correlated by their physical positions within the protein-coding mitochondrial genes to use these polymorphic points as a filter to obtain the RPKM of each patient sample that crossed the physical position of these points. We averaged the expression of each signal hub in all tissues based on the physical position of the polymorphic sites so that only the signal hubs contained within the physical position were selected. The final data set was separated into two categories based on sex (Female or Male); and based on age, with three great groups, 20-49 years, 50-59 years, and 60-79 years, also considering the sex (Female or Male). We estimate the RPKM mean of the genes and the polymorphic sites in each tissue sample. We also performed the statistical Tukey test for multiple comparisons, using the mean expression of each gene or polymorphic sites in each tissue for the three ages to calculate the RPKM mean of each age group. The False Discovery Rate was also calculated to conceptualize multiple comparisons in the analyzed data. Therefore, the FDR tests using the Benjamini-Hochberg method had a cut-off of ≤ 0.05. This analysis was performed for females and males separately.

### Pan-Cancer Survival Analysis of Mitochondrial Protein-Coding Genes

We used data from the GEPIA2 (http://gepia2.cancer-pku.cn/#index), a platform containing public data for interactive analysis of gene expression profiles in RNA-Seq samples from primary cancers from the TCGA (The Cancer Genome Atlas Program https://www.cancer.gov/about-nci/organization/ccg/research/structural-genomics/tcga), a reference cancer genomics program, covering 33 cancer types (Table S3), with a total of 9,736 samples. We also used data from samples from RNA-Seq experiments available in nontumor primary tissues from the Genotype-Tissue Expression (GTEx) project (8.587 samples) from 52 tissues (Table S2). A computational meta-analysis was performed to determine a pan-cancer overall survival map of the mitochondrial genes in 33 cancer types (Table S3). We set the following parameters: Survival Time Units: months, significance level: 0.05, P-Value Adjustment: False Discovery Rate (FDR), group cutoff: median, cutoff-high: 50% used. We define gene signatures as the additive expression profiles of at least three genes exhibiting the prognostic value per tissue.

### Prognostic Factors of Pan-Cancer Survival Outcomes

The prognostic factor values of the 13 mitochondrial protein-coding genes were estimated by the Kaplan-Meier method using a computational meta-analysis in RStudio, P (version 2022.02.3 build 492 for Windows) (Allaire, 2012) using the UCSC Xena Shiny standalone application (WANG et al., 2021) (http://xena.ucsc.edu/). We queried the expression profiles of each gene using four methods of survival outcomes: overall survival (OS), disease-specific survival (DSS), disease-free interval survival (DFI), and progression-free interval (PFI). OS refers to the percentage of patients alive after cancer diagnosis throughout the study period. DSS describes the percentage of patients who have not died from a cancer type in a defined period, such as patients who died from causes other than the specific cancer type is not counted in this measurement. DFI is the length of time after primary treatment for cancer that the patient survives without any signs or symptoms of that cancer. PFI is the length of time, during and after the treatment of a cancer type, that a patient lives with the disease but does not worsen. We set the parameters s described in the prior session and did not categorize the patients by age. We also investigated whether there is a sex-specific effect on survival outcomes.

### Molecular Profile Analysis for Risk or Protective Effects in Different Cancers

For this analysis, we used mRNA data expression for 33 different types of cancer available through the TCGA for all samples (Table S3). In this analysis, we constitute a gene signature that presents at least three genes per tissue. Essentially, a statistical regression model commonly used in medical research, the Cox model, was used to investigate the association between patient survival time and one or more predictor variables, in this case, the risk or protective effect. Therefore, the effect of covariates estimated by any proportional hazard model can be reported as a hazard ratio. In this way, we aim to identify whether the mitochondrial protein-coding genes have risk or protective effects based on OS (Overall Survivor) having an adjusted threshold of 0.5. We define genes as risky (log (Hazard Ratio) > 0) or protective (log (Hazard Ratio) < 0), or NS (No statistical significance, P-value > 0.05)).

### Analysis of the association of the molecular profile to immune response signatures and Stemness

Subsequently, we investigated the association between the molecular profile of mitochondrial protein-coding genes and stemness (the impact of a stem cell-like tumor phenotype). Here, we aim to obtain a possible relationship, positive (+1), negative (−1), or neutral (0), between gene signatures and Stemness in all available cancer types (Table S3). We also investigate the existence of a correlation between the mitochondrial protein-coding genes and the immune infiltrate signatures. In both analyses, genetic signatures presented at least three genes per tissue or at least three genes per tissue in each immune cell. For both analyses, we used mRNA expression data from patients of Pan-Cancer Atlas for 33 different types of cancer. Spearman’s test was performed to assess the correlation of the signatures, positive (≥0.50) or negative (≤ -0.50). We also filter this data considering its p-value, admitting only data with a p-value lower than 0.05.

## RESULTS

### Differential Expression of Mitochondrial Protein-Coding Genes

We compared the RNA expression profiles of the 13 mitochondrial protein-coding genes in tumor tissues from the TCGA project and matched (n=17) normal tissues from the GTEX project to identify disease conditions in which the genes are differentially expressed. Eleven mitochondrial protein-coding genes are DEG in brain tissue (*MT-ND2*, *MT-ND1*, *MT-ATP8*, *MT-ATP6*, *MT-CO2*, *MT-CYB*, *MT-CO3*, *MT-ND4L*, *MT-ND4*, *MT-ND3*, *MT-CO1*) with LogFC ≥1.5 (p ≤0.05). We also noticed other differentially expressed genes with LogFC ≥1.5 in tissues like the breast (*MT-ATP8*, *MT-ND1*, *MT-ND2*, *MT-CYB*, p ≤0.05), liver (*MT-ND4*, *MT-ATP6*, *MT-ATP8*, *MT-ND4L*, *MT-CYB*, *MT-ND2*, p ≤0.05), pancreas (*MT-ND2*, p ≤0.05) and testis (*MT-ND1*, p ≤0.05). We also observed a LogFC ≤ -1.5 in tissues such as peripheral blood (*MT-ND4*, *MT-ND5*, *MT-CO1*, *MT-ND4L*, p ≤0.05), ovary (*MT-ND6*, *MT-ND1*, *MT-CO1*, p ≤0.05) and thyroid (*MT-ND6*, p ≤0.05) (Figure 1) (Table S4).

**Figure 1.**
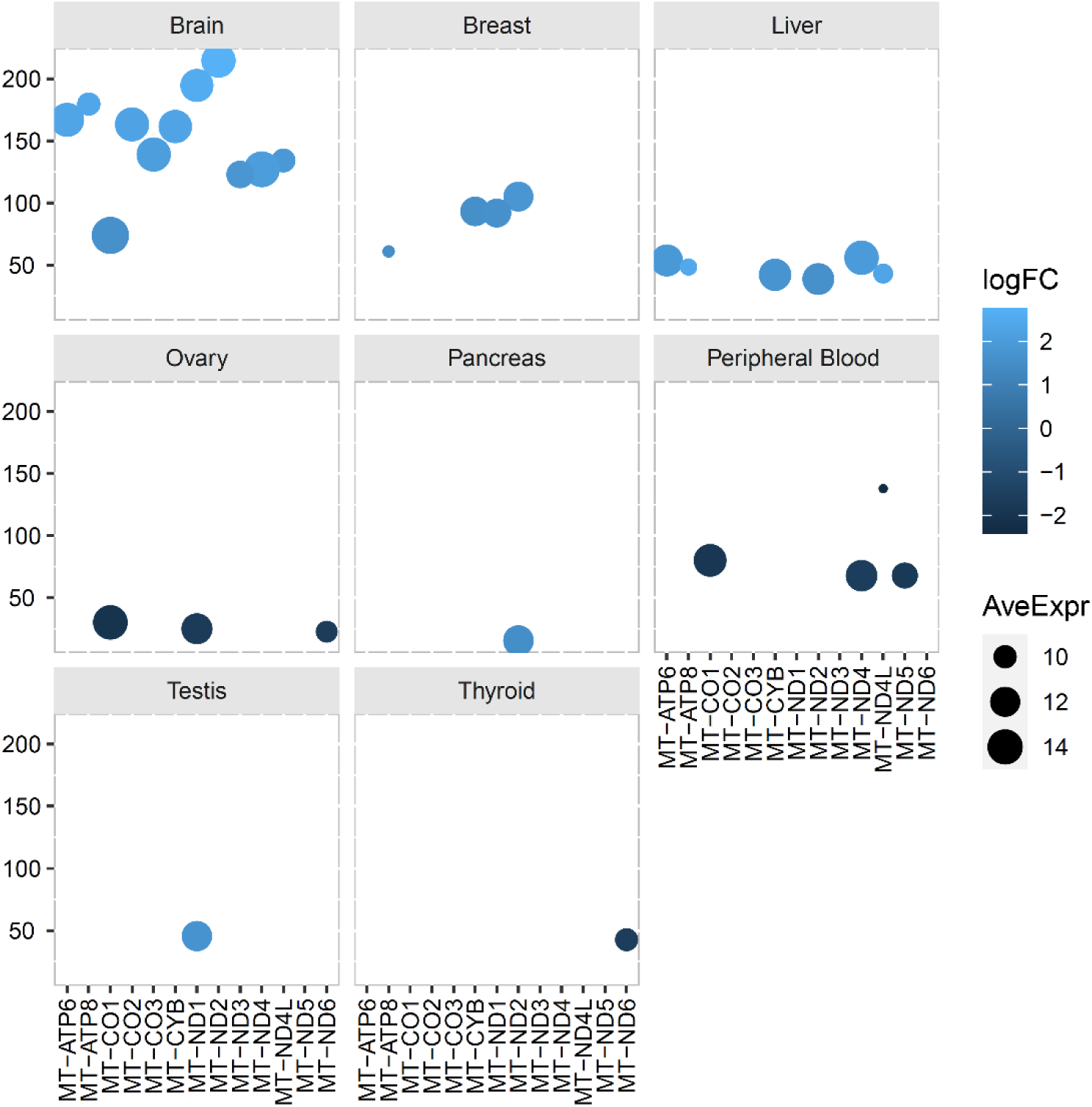
Balloon plot of the differential expressed mitochondrial protein-coding genes in normal versus cancer tissues. Balloon size refers to the average expression (norm_counts); blue color intensity refers to the Log2FC. Overexpression (+); underexpression (-). X-axis: the 13 mitochondrial protein-coding genes; Y-axis, FDR (-log10). The tissues are indicated in each box.

### In Silico Consolidation of the Mitochondrial Protein-Coding Genes and Polymorphic sites

From the names of the mtDNA genes, their physical positions, and to which chain within the mitochondrial genome they belonged (H-heavy; L-light) were obtained, as well as the product of each gene. Subsequently, still using the physical positions of the mtDNA genes, 2297 polymorphic sites were obtained in the mitochondrial genome, of which only 1540 were within the 13 mitochondrial genes (MT-ND1 = 156, MT-ND2 = 126, MT-ND3 = 46, MT-ND4 = 157, MT-ND4L = 39, MT-ND5 = 230, MT-ND6 = 81, MT-CO1 = 162, MT-CO2 = 82, MT-CO3 = 108, MT-CYB = 191, MT-ATP6 = 112, MT-ATP8 = 37) (Figure S1).

### Expression Profile of Sex-specific Aging-related in Mitochondrial Protein-Coding Genes and Polymorphic sites

We compared the expression profiles of the 13 mitochondrial protein-coding genes and polymorphic sites in males and females according to three age groups (20-49-year-old, 50-59-year-old, and 60-79-year-old) in 52 GTEx tissues. We used two query strategies: first, we used the physical coordinates of each gene to extract the normalized transcript expression levels (RPKM) as a metric of the abundance of each donor in the mitochondria genome, and second, we used the physical position of polymorphic sites in population from the dbSNP151 database to capture site-specific abundance. We observed sex-biased effects on expression profiles across different tissues.

Eleven genes (*MT-ND6*, *MT-ND5*, *MT−ND4*, *MT-ND4L*, *MT−ND3*, *MT−ND2*, *MT−ND1*, *MT−CYB*, *MT−CO3*, *MT−CO1*, *MT−ATP6*; Tukey test, p≤0.05) showed differences in expression profile of males in thirty-four tissues across different ages (Figure 2) (Table S5-S6). The most significant male-biased expression across different ages was noticed in *MT-ND5* for Artery – Aorta, Brain – Substantia Nigra, and Esophagus – Mucosa (Wilcox test, p ≤0.001) and in the *MT-ND6* for Brain – Hypothalamus and Kidney – Cortex (Wilcox test, p ≤0.001) (Table S5-S6). For females, ten genes (*MT−ND6*, *MT−ND5*, *MT−ND4*, *MT−ND2*, *MT−ND1*, *MT−CYB*, *MT−CO3*, *MT−CO1*, *MT−ATP8*, *MT−ATP6*; Tukey test, p≤0.05) exhibited differences in expression profile for nineteen tissues across different ages (Figure 3) (Table S5-S6). The most significant female-biased expression across different ages was observed in *MT-ATP6*, *MT-ATP8*, and *MT-CO1* for Artery – Coronary, and in *MT-CO1*, *MT-CO3*, and *MT-CYB* for Adipose – Visceral (Omentum) (Wilcox test, p ≤0.05). We also noticed that *MT-ND2* exhibit a highly significant female-biased expression in Brain – Frontal Cortex (BA9) and Brain – Substantia Nigra across different ages (Wilcox test, p ≤0.001) and in *MT-ND1* only for Brain – Substantia Nigra (Wilcox test, p ≤0.001). Interestingly, the *MT-ND5* also exhibited a female-biased expression in Artery – Aorta, Brain – Frontal Cortex (BA9), and Colon – Transverse (Wilcox test, p ≤0.001) at different ages, and the same was noticed in *MT-ND6* for Brain – Frontal Cortex (BA9) and Brain – Substantia Nigra (Wilcox test, p ≤0.001) (Table S5-S6).

**Figure 2.**
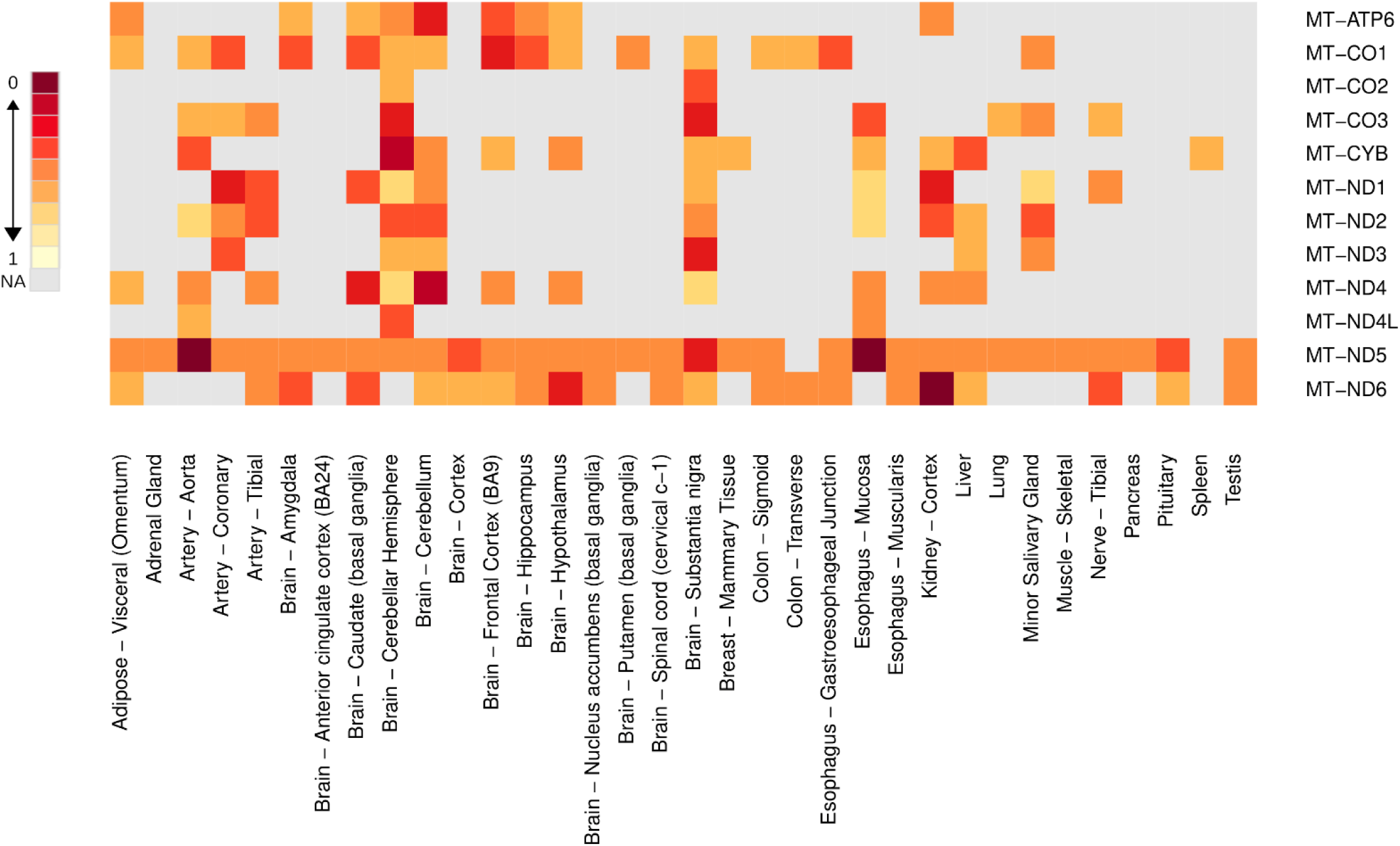
Heatmap of significant mitochondrial protein-coding gene expression across different ages for each tissue in males. The expression of all mitochondrial protein-coding genes was measured using RPKM (reads per kilobase million), and the color scale denotes the level of significance of gene expression in each tissue. The different ages consisted of three groups (20-49 years; 50-59 years, and 60-79 years). The significance level was measured based on the Tukey test and the FDR adjustment, Benjamin-Hochberg.

**Figure 3.**
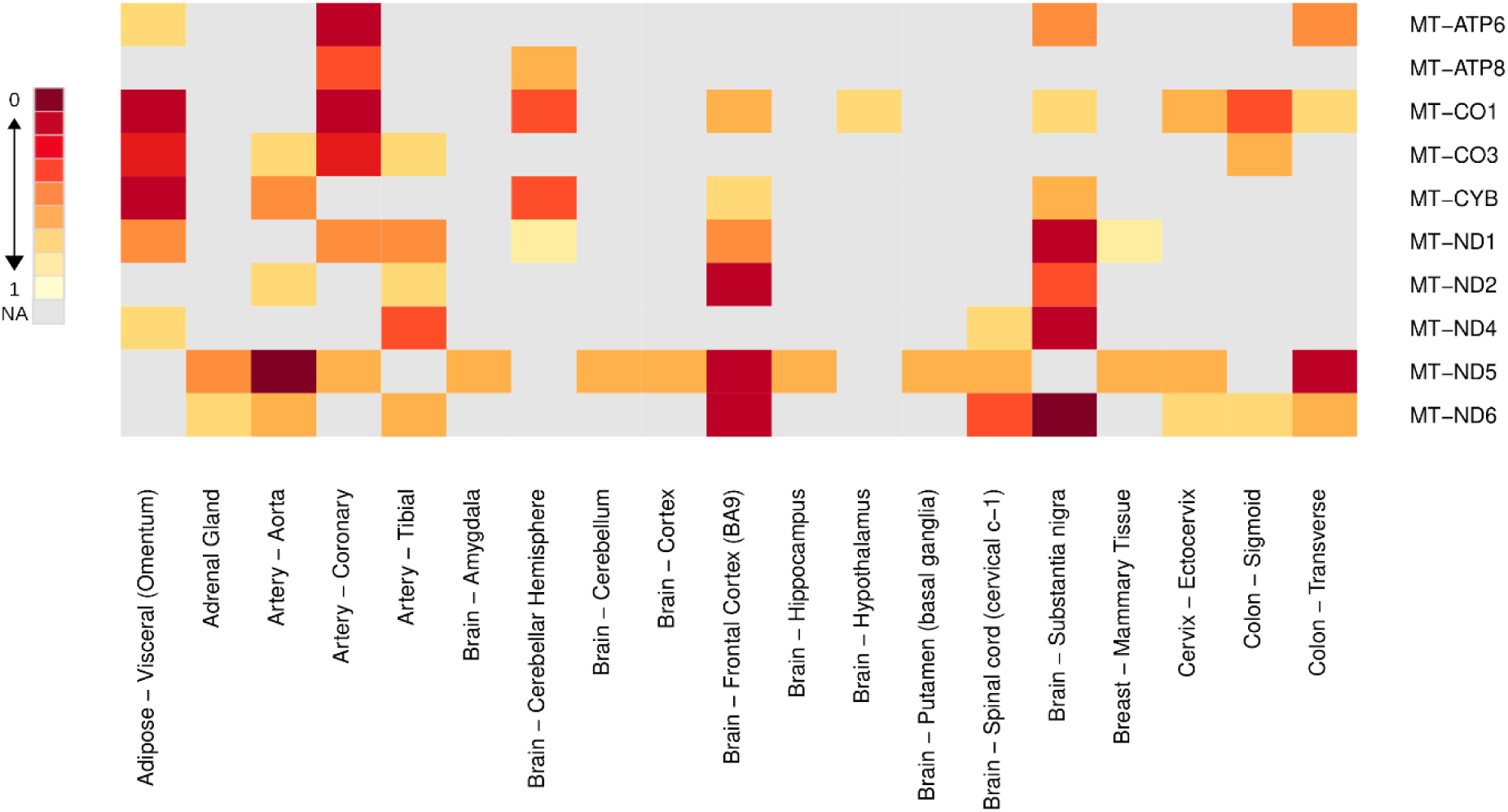
Heatmap of significant mitochondrial protein-coding gene expression across different ages for each tissue in females. The expression of all mitochondrial protein-coding genes was measured using RPKM (reads per kilobase million), and the color scale denotes the level of significance of gene expression in each tissue. The different ages consisted of three groups (20-49 years; 50-59 years, and 60-79 years). The significance level was measured based on the Tukey test and the FDR adjustment, Benjamin-Hochberg.

Considering possible differences in expression between the 52 different tissues analyzed for the 13 mitochondrial protein-coding genes in the three age groups, we observed sex-biased and age-group-dependent effects on expression profiles (Figure 4-6). In the 20-49-year-old group, thirty-four different tissues showed significantly different expression profiles for *MT-ATP6*, *MT-ATP8*, *MT-ND1*, *MT-ND2*, *MT-ND3*, *MT-ND4*, *MT-ND4L*, *MT-ND5*, *MT-ND6*, *MT-CYB*, *MT-CO1*, *MT-CO2*, *MT-CO3* (Wilcox test, p ≤0.05). In this age group, eight genes exhibited the most significant expression profiles in Artery - Coronary (*MT-ATP6*, *MT-ATP8*, *MT-CO1*, *MT-CO2*, *MT-ND1*, *MT-ND2*, *MT-ND4*, and *MT-ND5*; Wilcox test, p≤0.001). For the 50-59-year-old group, we noticed nineteen different tissues where 13 genes exhibited the same effect (Wilcox test, p≤0.05). For this age group, the most significant expression profile between the tissues occurred in *MT-CO3*, *MT-CYB*, and *MT-ND6* in Adipose – Subcutaneous (Wilcox test, p≤0.001); *MT-ATP6*, *MT-ATP8*, *MT-CO1*, *MT-CO3*, and *MT-ND5* in Adipose – Visceral (Omentum) (Wilcox test, p≤0.001); *MT-ATP6*, *MT-ATP8*, *MT-CO1*, *MT-CO2*, *MT-ND1*, *MT-ND2*, *MT-ND5*, and *MT-ND6* in Brain – Cerebellar Hemisphere (Wilcox test, p≤0.001); *MT-CO1*, *MT-ND4*, *MT-ND4L*, and *MT-ND6* in Brain – Hippocampus (Wilcox test, p≤0.001); *MT-ATP8*, *MT-CO1*, *MT-CO3*, *MT-CYB*, *MT-ND4* and *MT-ND5* in Brain – Hypothalamus (Wilcox test, p≤0.001); *MT-ND1*, *MT-ND4*, and *MT-ND6* in Brain – Nucleus accumbens (basal ganglia) (Wilcox test, p≤0.001); *MT-ND1* in Brain – Substantia Nigra, (Wilcox test, p≤0.001). The last age group, 60-79-year-old, exhibited thirty-four tissues with differences in expression for the same 13 mitochondrial protein-coding genes (Wilcox test, p ≤0.05). In this last age group, the most significant expression profile between the tissues and the mitochondrial protein-coding genes happens in *MT-CO1*, *MT-CO3*, *MT-CYB*, and *MT-ND5* in Adipose – Visceral (Omentum) (Wilcox test, p≤0.001); *MT-ND4L* and *MT-ND5* in Artery – Coronary (Wilcox test, p≤0.001); *MT-ATP6*, *MT-ATP8*, *MT-CO1*, *MT-CO2*, *MT-ND1*, *MT-ND2*, *MT-ND4*, *MT-ND4L*, *MT-ND5* and *MT-ND6* in Brain – Amygdala (Wilcox test, p≤0.01); *MT-ND4* and *MT-ND5* in Brain – Putamen (basal ganglia) (Wilcox test, p≤0.05); *MT-CO1*, *MT-CO3*, *MT-ND4* and *MT-ND5* in Colon – Sigmoid (Wilcox test, p≤0.01); *MT-ATP6*, *MT-ATP8*, *MT-CO1*, *MT-ND1*, *MT-ND2*, *MT-ND4* and *MT-ND6* in Esophagus – Muscularis (Wilcox test, p≤0.01); MT-ND1, MT-ND2, MT-ND5 and MT-ND6 in Kidney – Cortex (Wilcox test, p≤0.05); *MT-ATP8*, *MT-CO1*, *MT-CO2*, *MT-CYB*, *MT-ND1*, *MT-ND3*, *MT-ND4*, *MT-ND5* and *MT-ND6* in Minor Salivary Gland (Wilcox test, p≤0.001); *MT-CO1*, *MT-ND1*, *MT-ND5* and *MT-ND6* in Small Intestine – Terminal Ileum (Wilcox test, p≤0.01); *MT-CYB*, *MT-ND4* and *MT-ND5* in Thyroid (Wilcox test, p≤0.01).

**Figure 4.**
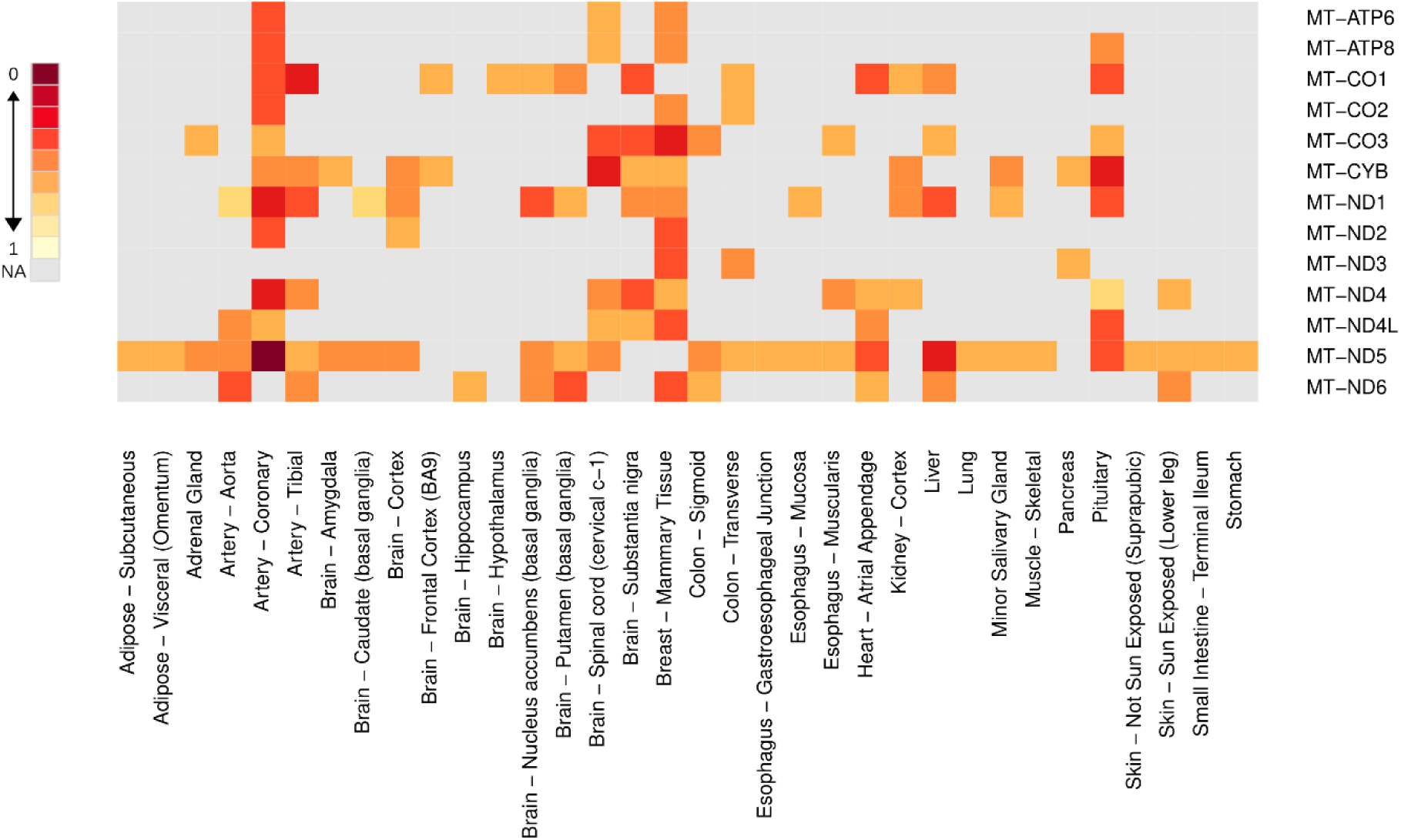
Heatmap of significant mitochondrial protein-coding gene expression between sexes in 20-49-year-old group. All mitochondrial protein-coding genes were measured between the sexes (female and male) using RPKM (reads per kilobase million) across different tissues, and the color scale denotes the level of significance of gene expression in each tissue. The sex-specific tissue samples were excluded, and the significance level was measured based on the Wilcox test and the FDR adjustment, Benjamin-Hochberg.

**Figure 5.**
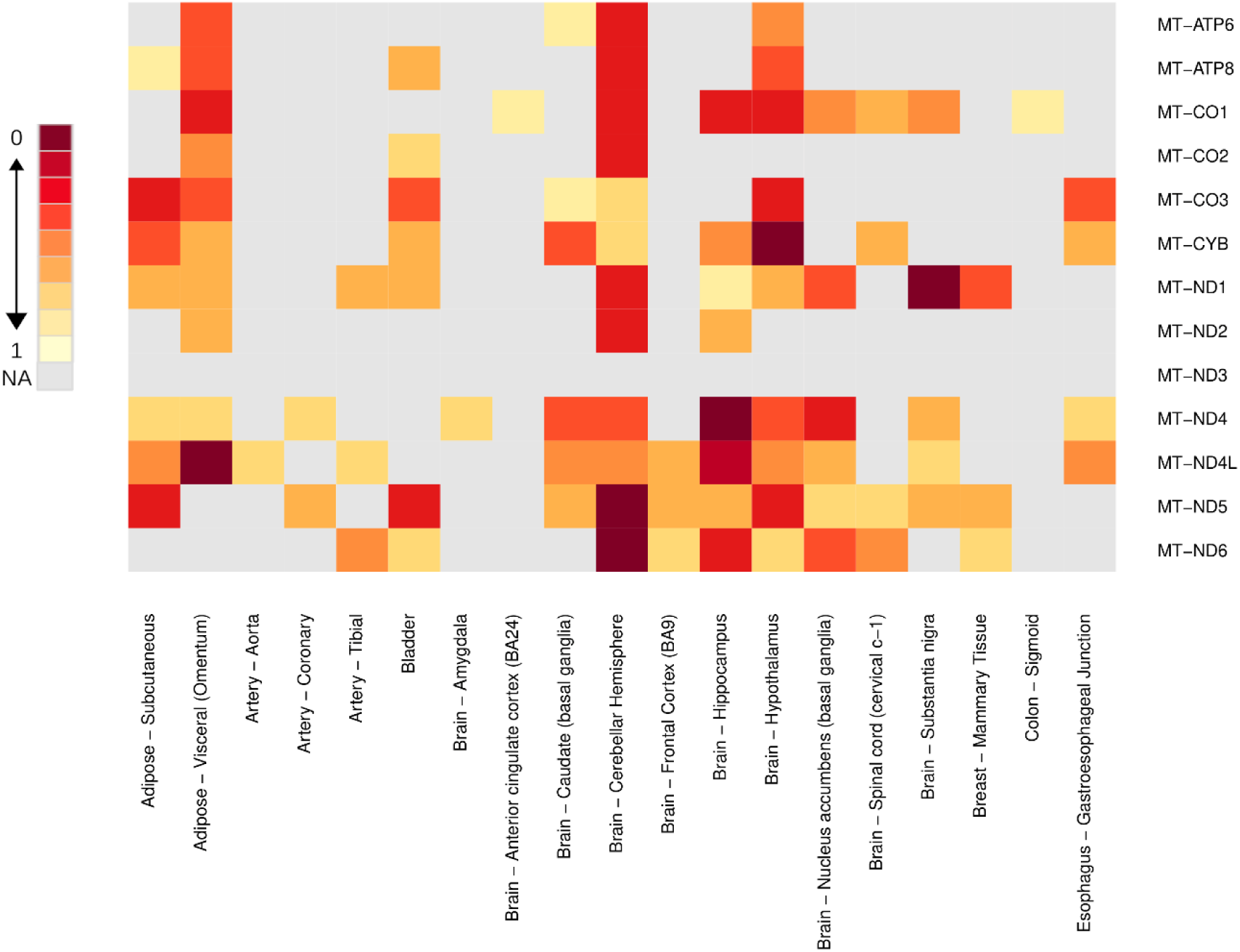
Heatmap of significant mitochondrial protein-coding gene expression between sexes in 50-59-year-old group. All mitochondrial protein-coding genes were measured between the sexes (female and male) using RPKM (reads per kilobase million) across different tissues, and the color scale denotes the level of significance of gene expression in each tissue. The sex-specific tissue samples were excluded, and the significance level was measured based on the Wilcox test and the FDR adjustment, Benjamin-Hochberg.

**Figure 6.**
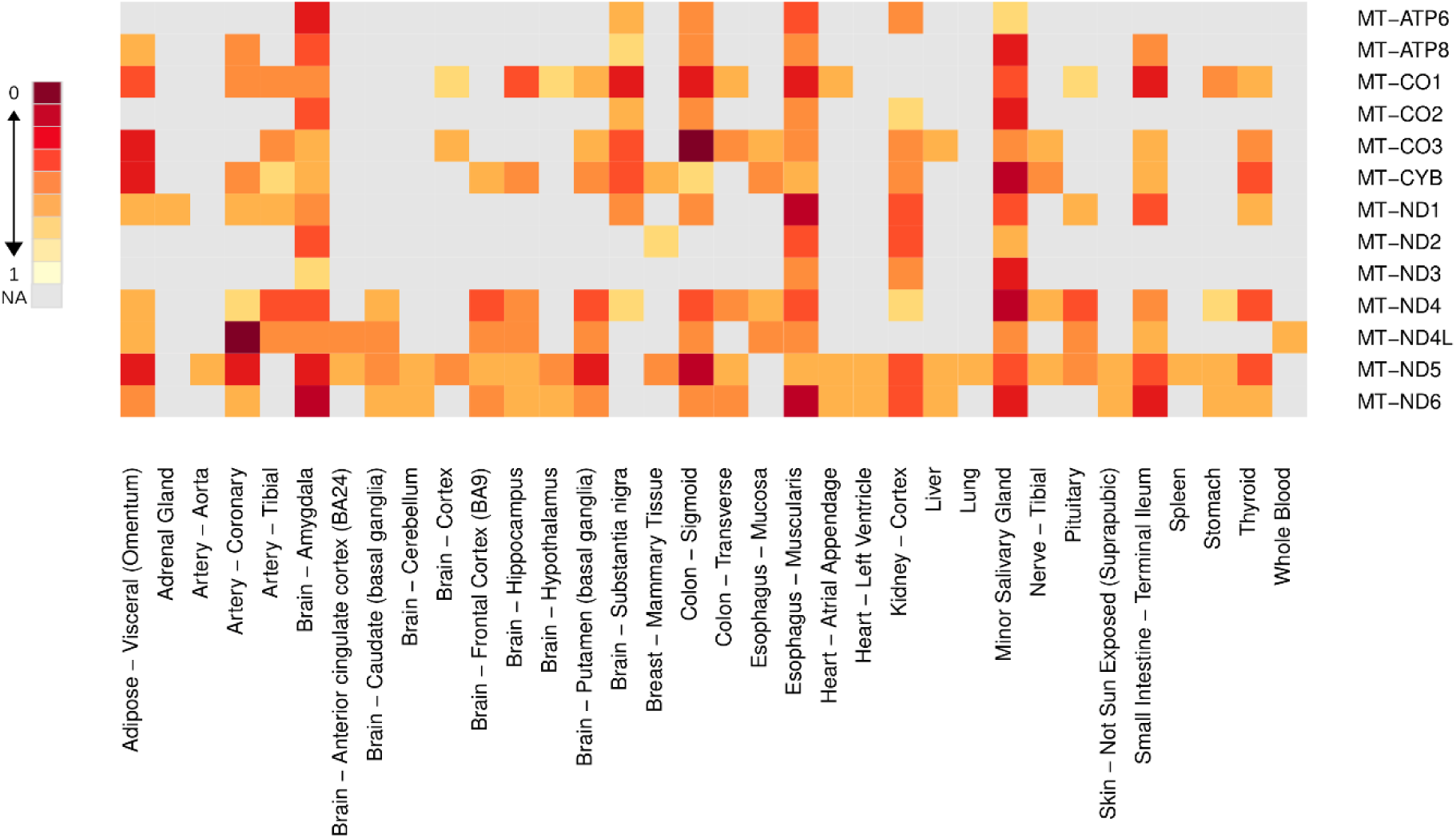
Heatmap of significant mitochondrial protein-coding gene expression between sexes in 60-79-year-old group. All mitochondrial protein-coding genes were measured between the sexes (female and male) using RPKM (reads per kilobase million) across different tissues, and the color scale denotes the level of significance of gene expression in each tissue. The sex-specific tissue samples were excluded, and the significance level was measured based on the Wilcox test and the FDR adjustment, Benjamin-Hochberg.

The polymorphic profile of the variation points in population for the 13 mitochondrial protein-coding genes was clustering in 52 tissues from the GTEx project comprising three different ages (20-49-year-old, 50-59-year-old, 60-79-year-old), which allowed building a landscape of these polymorphic sites related to sexes (males and females) (Figure 7 and 8). This landscape will provide information about the variation of polymorphic points along the transcripts and whether this variation is global or sustained at all points in the population by using the RPKM transcript. We identify an age-group-dependent sex-biased effect on the polymorphic sites in females in ten different tissues among the three age groups (Adipose – visceral (Omentum), Adrenal Gland, Artery Coronary, Brain – Amygdala, Brain – Frontal cortex (BA9), Brain – Hippocampus, Brain – Putamen (basal ganglia), Brain – Spinal cord (cervical c-1), Brain – Substantia nigra, Breast – Mammary tissue, Cervix – Ectocervix; Tukey test, p ≤0.05). In the same way, an age-male-biased of polymorphic sites was found in eight tissues (colon– sigmoid, colon – transverse, esophagus – gastroesophageal junction, liver, lung, muscle-skeletal, nerve tibial, prostate; Tukey test, p ≤0.05).

**Figure 7.**
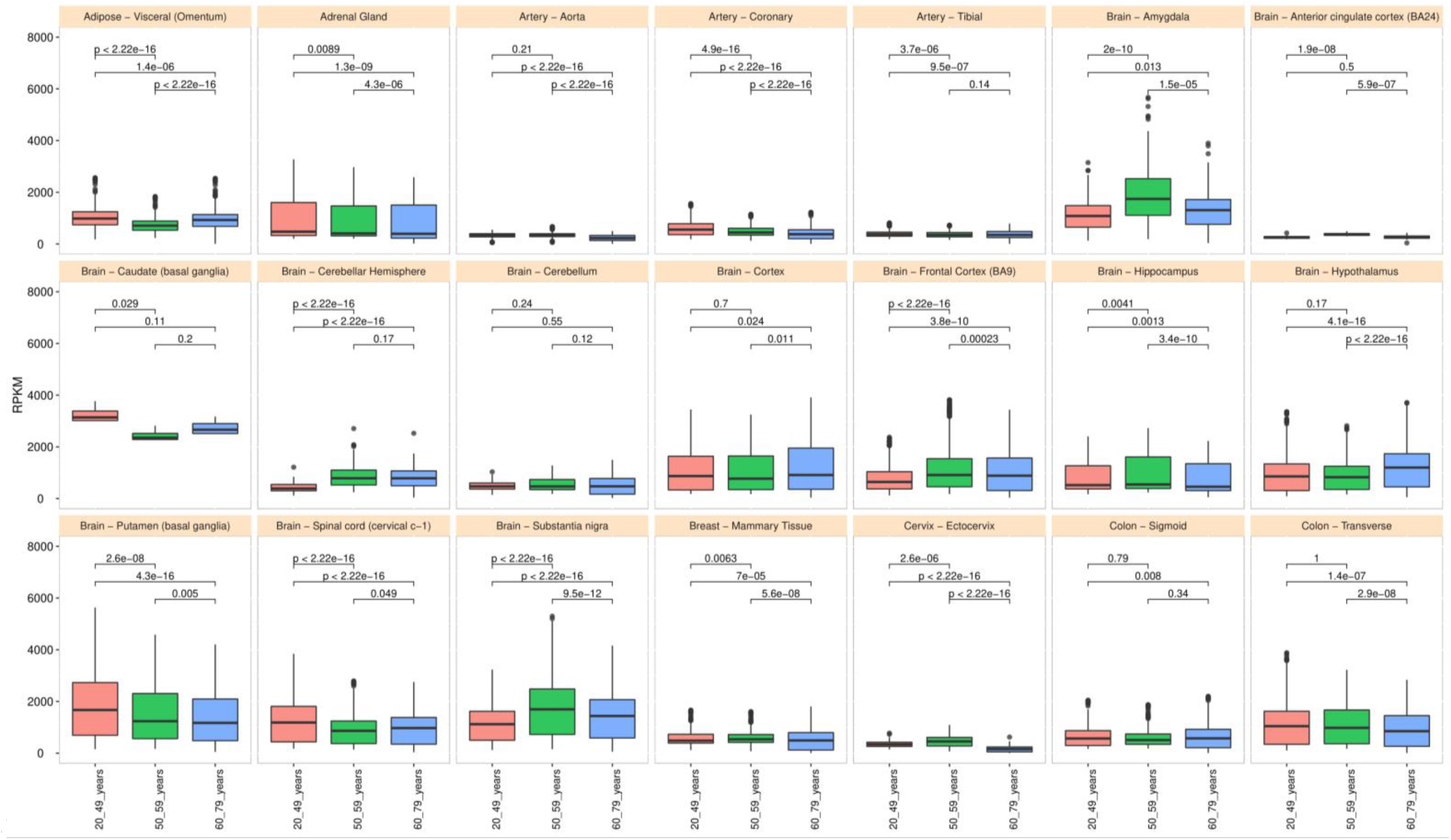
Boxplot of mitochondrial polymorphic site expression in females across tissues at different age groups. The polymorphic site expression at the mitochondrial protein-coding genes was measured using RPKM (reads per kilobase million) in three age groups, 20-49 years (red), 50-59 years (green), and 60-79 years (blue). The Tukey test and the FDR adjustment, Benjamini-Hochberg assessed significant differences between age groups.

**Figure 8.**
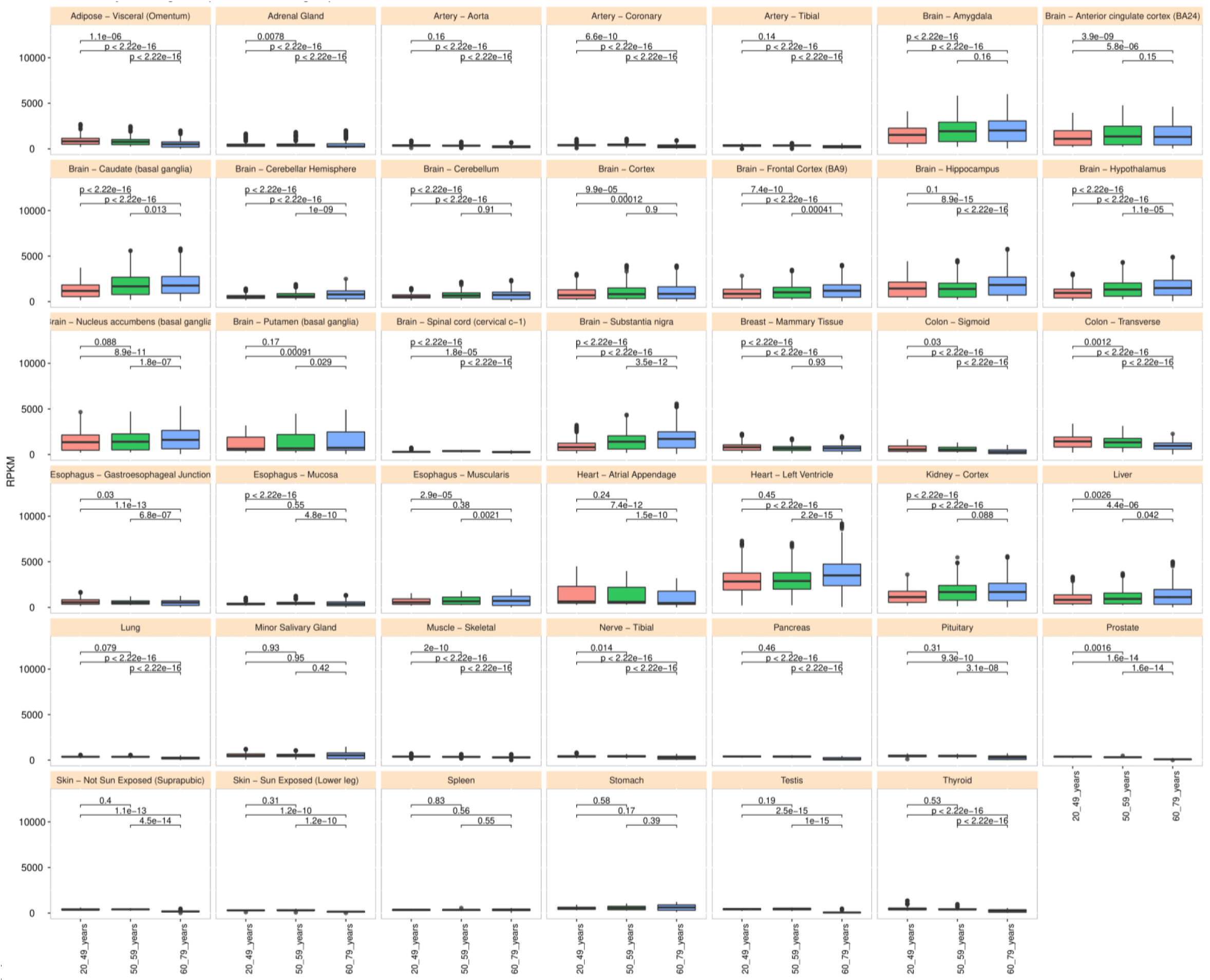
Boxplot of mitochondrial polymorphic site expression in males across tissues at different age groups. The polymorphic site expression at the mitochondrial protein-coding genes was measured using RPKM (reads per kilobase million) in three age groups, 20-49 years (red), 50-59 years (green), and 60-79 years (blue). The Tukey test and the FDR adjustment, Benjamini-Hochberg assessed significant differences between age groups.

### Pan-Cancer Survival Map Analysis Based on the Expression Status of the 13 Mitochondrial Protein-coding Genes

We estimated the overall survival outcome for each of the molecular expression profiles of the 13 mitochondrial protein-coding genes in 33 cancer types to map the hazard risk based on the Cox proportional-hazards model for overexpression and under-expression patient groups. The under-expression of the *MT-CO1*, *MT-CO2*, *MT-ND5*, and *MT-ND6* genes is a hazard risk in lower-grade glioma (hazard ratio range 0.41-0.51, Logrank p ≤0.00026, p(HR) ≤0.00033), while the under-expression of the MT-CO3 gene is a hazard risk in pancreatic adenocarcinoma (PAAD) (hazard ratio 0.45, p(HR)=0.00022, Log-rank p=0.00017), Figure 9 (Table S10).

**FIGURE 9.**
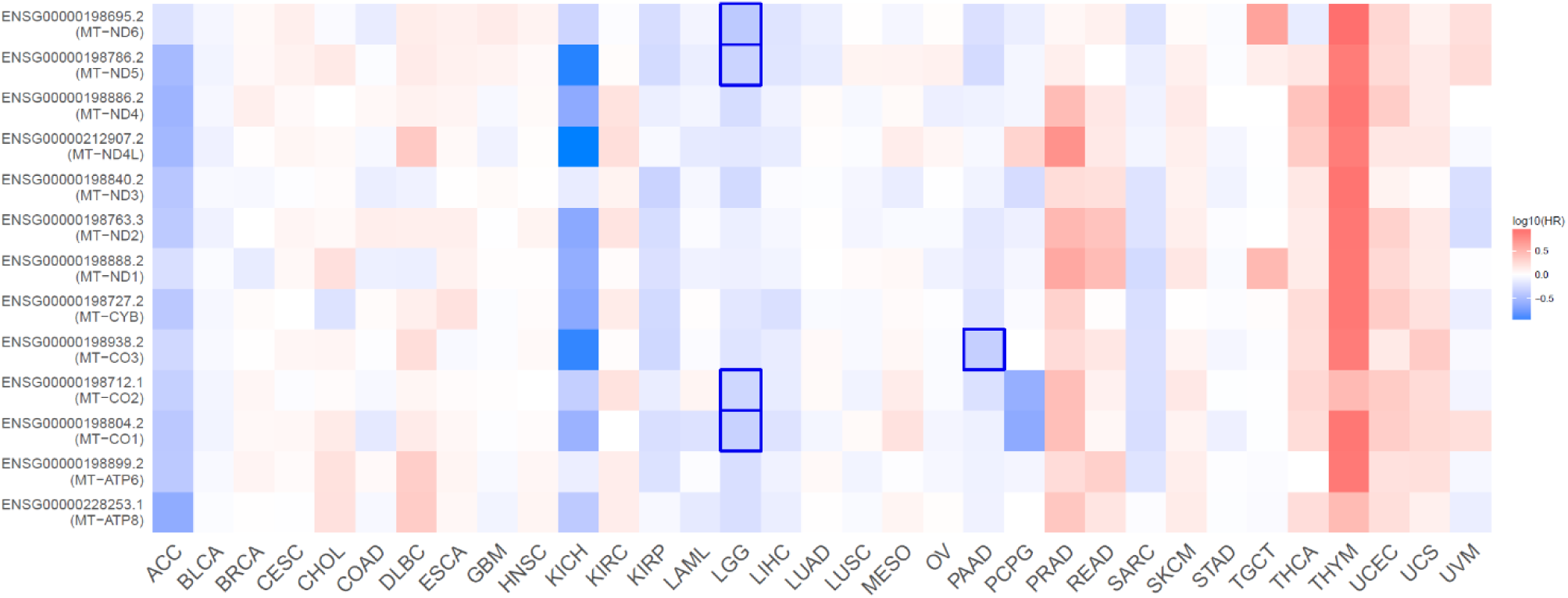
Overall survival heatmap for the 13 mitochondrial protein-coding genes. Squares denote overall survival associated with overexpression (red) or under-expression (blue). Highlighted frames indicate significant associations with overall survival outcomes based on hazard ratios (logarithmic scale log10). Y-axis: mitochondrial genes (Ensemble transcripts and UCSCXena names), X-axis: pan-cancer types. The under-expression profiles of the MT-CO1, MT-CO2, MT-ND5, and MT-ND6 genes have a significant overall survival risk in LGG (hazard ratio range 0.41-0.51, Logrank p ≤0.00026, p(HR) ≤0.00033), while the under-expression of the MT-CO3 gene is a hazard risk in PAAD (hazard ratio 0.45, p(HR)=0.00022, Logrank p=0.00017).

### The 4-component gene signature is a Prognostic Factor of Survival Outcomes in LGG

We estimated the prognostic factor value of the 4-component gene signature (*MT-CO1*, *MT-CO2*, *MT-ND5*, and *MT-ND6*) by correlating the molecular expression profile with rates of survival outcomes in LGG patients (n=521). We used four methods of survival outcomes: overall survival (OS) and disease-specific survival (DSS), disease-free interval (DFI), and progression-free-interval (PFI). Overexpression of the 4-component gene signature was significantly correlated with OS (Log-rank p < 0.0001) (Figure 10, Table S11), DSS (Log-rank p < 0.0001) (Figure 11, Table S12), and PFI (Log-rank p = 2e−04) (Figure 12, Table S13). The favorable prognostic factor of the overexpression was not sex-specific since, in either males or females, the overexpression was significantly correlated with OS, DSS, and PFI (Log-rank p ≤ 0.0076) (Figure S2-S7; Tables S14-S19). However, for the DFI outcome, we observed a sex-specific association in that the downregulation of the 4-component gene signature in female patients exhibited a greater prognostic factor value (Log-rank p= 0.019) compared to males (Log-rank p= 0,14), Figure S8 and Figure S9 (Tables S20 and S21). There are no sufficient data (n ≤10 samples) about the clinical-pathologic stages (Stages I-IV) in LGG subjects. Therefore, we could not estimate the prognostic values with the clinical staging.

**FIGURE 10.**
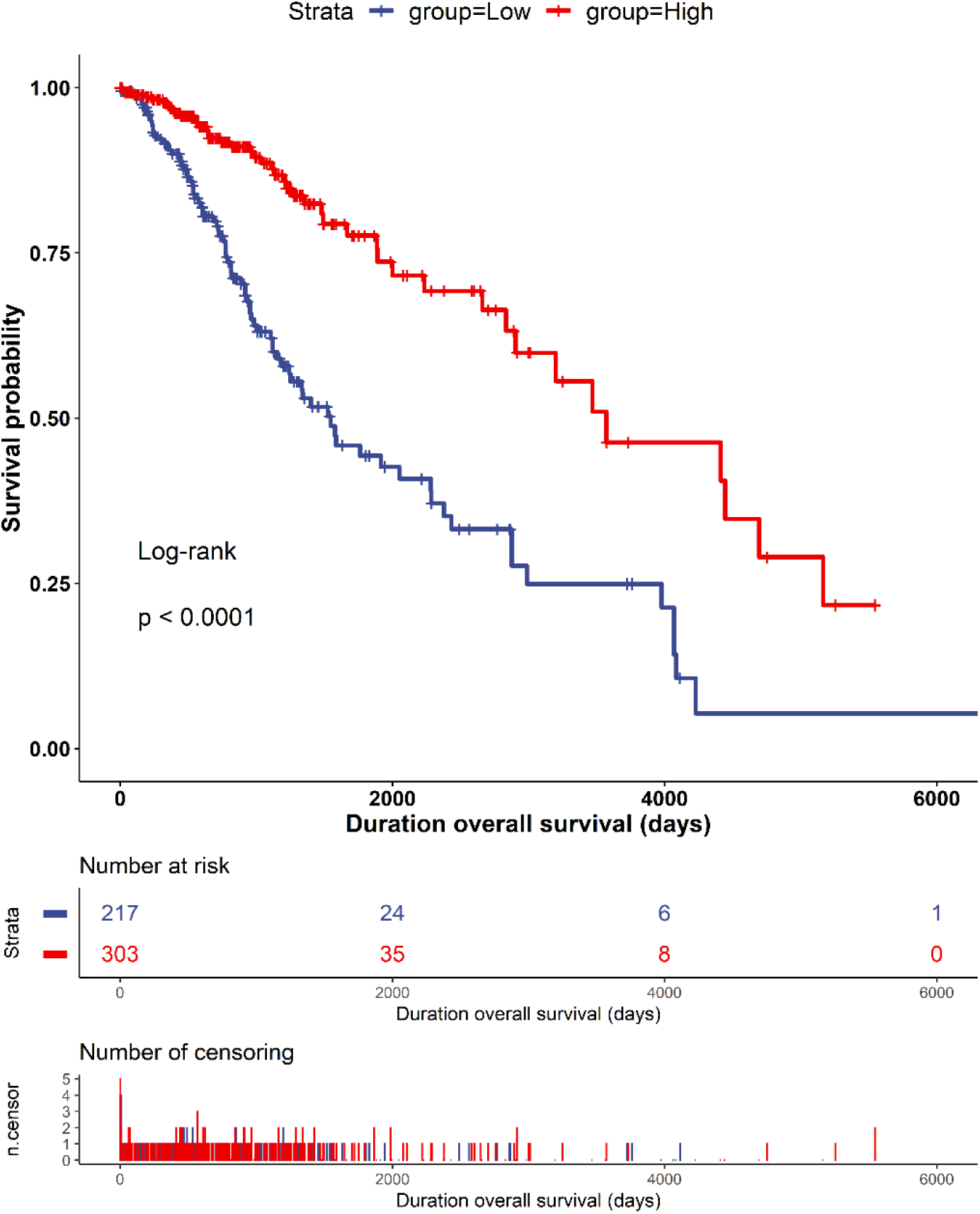
Kaplan-Meier plot of overall survival outcome in low-grade glioma. The curves denote the contribution of the 4-component gene signature (*MT-CO1*, *MT-CO2*, *MT-ND5*, *MT-ND6*) to the overall survival outcome in patients from the low (blue – downregulated) and high (red – upregulated) expression groups. The censoring number refers to patients who did not suffer the outcome of interest during the specified study period. The overexpression of the signature has a more significant prognostic factor value.

**FIGURE 11.**
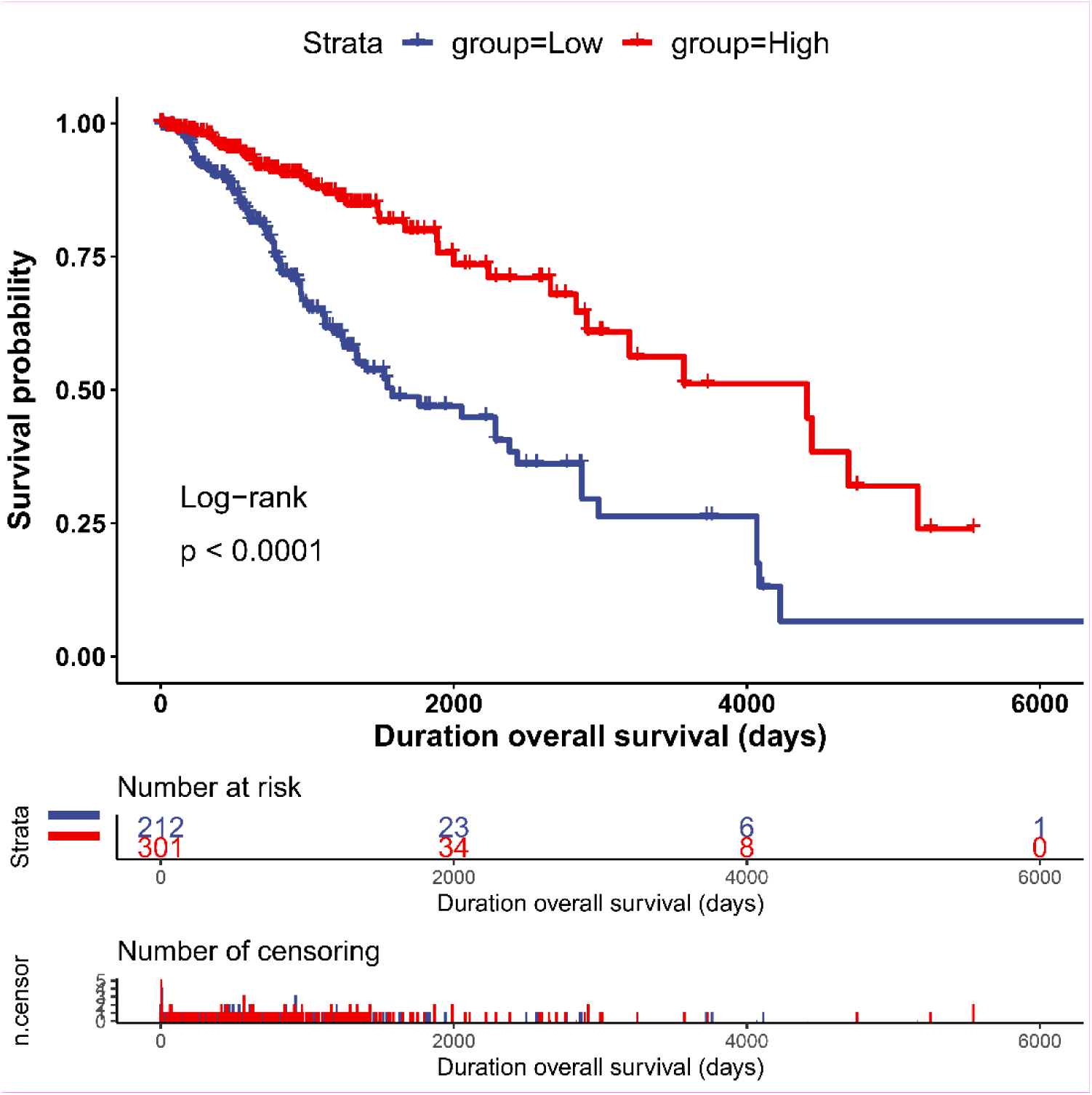
Kaplan-Meier plot of disease-specific outcome in low-grade glioma. The curves denote the contribution of the 4-component gene signature (*MT-CO1*, *MT-CO2*, *MT-ND5*, *MT-ND6*) to the disease-specific outcome in patients from the low (blue – downregulated) and high (red – upregulated) expression groups. The censoring number refers to patients who did not suffer the outcome of interest during the specified study period. The overexpression of the signature has a more significant prognostic factor value.

**FIGURE 12.**
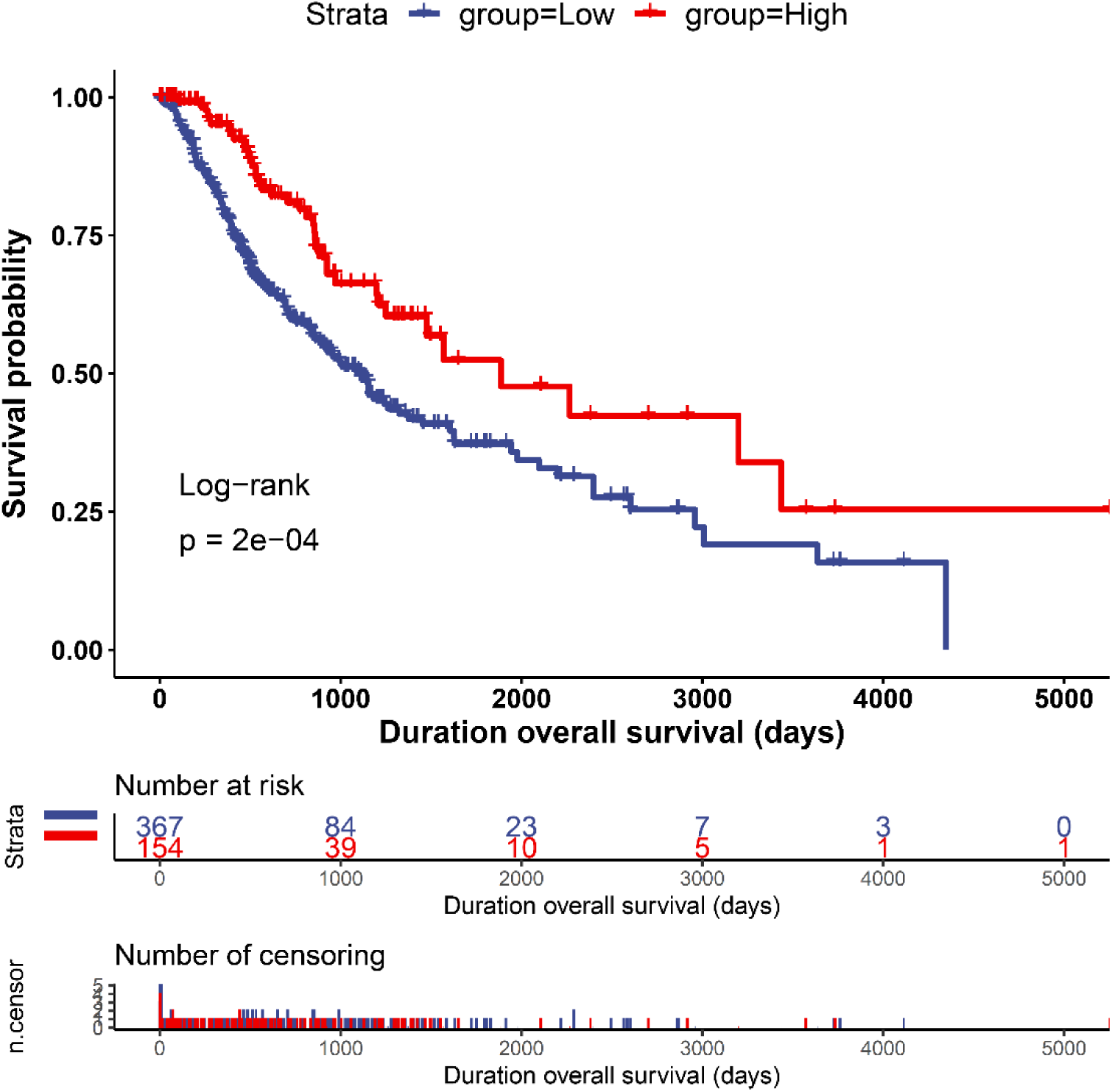
Kaplan-Meier plot of progression-free-interval outcome in low-grade glioma. The curves denote the contribution of the 4-component gene signature (*MT-CO1*, *MT-CO2*, *MT-ND5*, *MT-ND6*) to the progression-free-interval outcome in patients from the low (blue – downregulated) and high (red – upregulated) expression groups. The censoring number refers to patients who did not suffer the outcome of interest during the specified study period. The overexpression of the signature has a more significant prognostic factor value.

### Protective or Risk Effects of the 4-component Gene Signature in Different Cancers

We observed a protective effect of the 4-component gene signature in LGG (hazard ratio -0.17), kidney chromophobe (KICH – hazard ratio -0.32), kidney renal papillary cell carcinoma (KIRP - hazard ratio -0.10), adrenocortical carcinoma (ACC - hazard ratio -0.19), pancreatic adenocarcinoma (hazard ratio -0.08) and hepatocellular liver carcinoma (LIHC - hazard ratio -0.05) (Figure 12) (Table S12). In contracts, the signature has a risk effect on thymoma (THYM – hazard ratio 0.25) and uterine corpus endometrial carcinoma (UCEC – hazard ratio 0.06) (Figure 13; Table S22).

**FIGURE 13.**
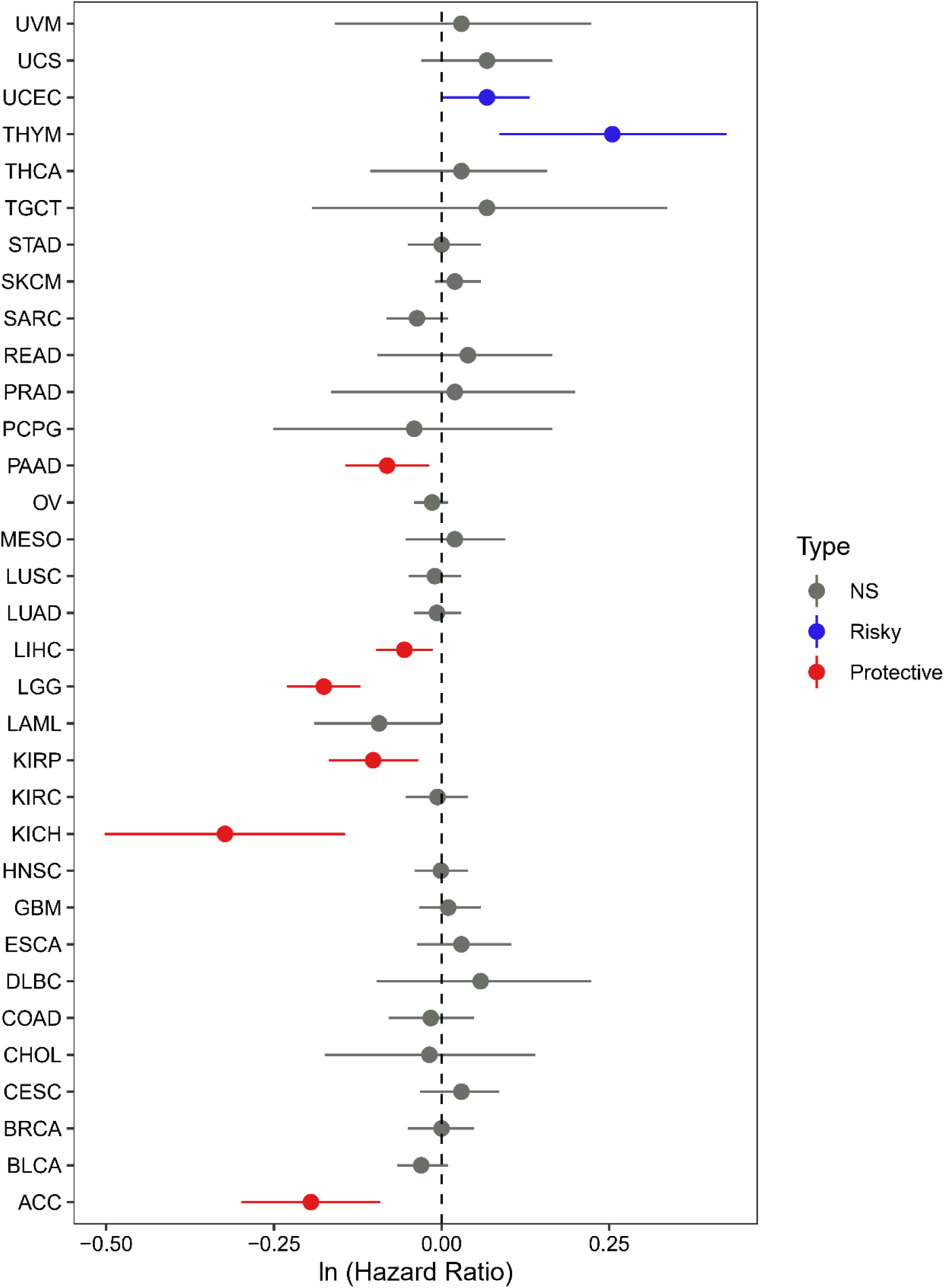
Boxplot of the molecular profile of mitochondrial protein-coding genes for protective or risk effects based on hazard ratio. Cox model analysis for the 4-component gene signature (*MT-CO1*, *MT-CO2*, *MT-ND5*, *MT-ND6*) *in 33* cancer types. The signature is protective (red) in *LGG*, *ACC*, *KICH*, *KIRP*, *LIHC*, *and PAAD* cancers (p-value < 0.0113) or risk (blue) in *THYM and UCEC* cancers (p-value < 0.0465).

### Correlation Analysis Between Molecular Profiles and Immune Infiltrates

We estimated the degrees of the correlation between the mRNA expression levels of the 13 mitochondrial protein-coding genes and the molecular expression profiles indicative of immune cell infiltrates in 33 cancer types (Table S23). For this analysis, we defined a gene signature as the molecular expression profile of a minimum of three genes exhibiting the same direction of correlation. The most extended, 13-component gene signature occurred in lymphoid neoplasm diffuse large B-cell Lymphoma (DBLC), being positively correlated (rho ≥ 0.50 p-value < 0.0003) with neutrophil, NK, T-cell CD4 memory, M1, /M2 macrophages, T-cell CD8+/CD4+, and B-cell memory cells (Figure 14).

**FIGURE 14.**
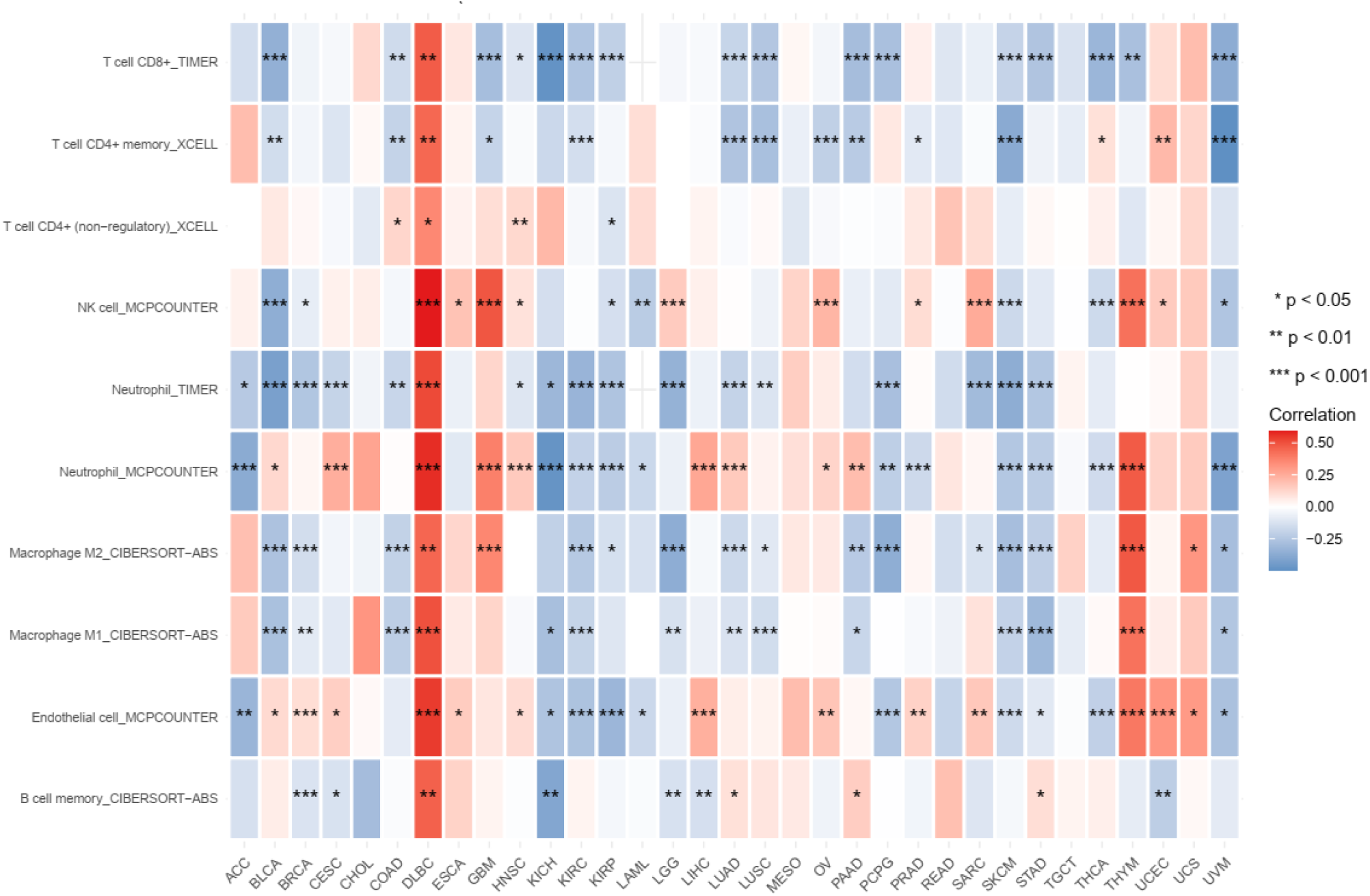
Heatmap of the correlations between the expression profiles of the 13-component gene signature and immune cell infiltrates. The red-blue heatmap scale shows the Spearman correlation coefficients’ significance levels (***, **, *). In DBLC, the 13-component gene signature is positively correlated with infiltrates of neutrophil, NK, T-cell CD4 memory, M1/M2 macrophages, T-cell CD8+/CD4+, and B-cell memory cells (rho ≥ 0.50 p-value < 0.0003).

The second-longest signature is a 7-component gene signature (*MT-ND4L*, *MT-CYB*, *MT-ND1*, *MT-ND4*, *MT-ND5*, *MT-ND6*, *MT-ATP8*), positively correlated (rho ≥0.50, p-value ≤ 0.001) with the following cell immune infiltrates in thymoma (THYM): endothelial cell, neutrophil, myeloid dendritic cell, T cell CD4+, and macrophage M2 (Figure S10) (Table S24).

In THYM, an 8-component gene signature (*MT-ATP8*, *MT-CO1*, *MT-CYB*, *MT-ND1*, *MT-ND2*, *MT-ND4*, *MT-ND4L*, *MT-ND5*) showed an inverse correlation with molecular profiles for the following cell infiltrates T cell CD4+ Th1, T cell, T cell regulatory (Tregs) (rho ≤ -0.5, p-value ≤ 3.05e-10), Figure S11 (Table S25).

We also estimated the degrees of the correlation between the mRNA expression levels for the 4-component gene signature (*MT-CO1*, *MT-CO2*, *MT-ND5*, *MT-ND6*) associated earlier with survival risk in LGG and the gene profiles indicative of immune cell infiltrates in 33 cancer types. The 4-component gene signature was positively correlated (rho ranging 0.06-0.6 p-value < 0.05) with 111 profiles of immune cell infiltrates and negatively correlated (rho ranging -0.56 to -0.059 p-value < 0.05) with 113 profiles of immune cell infiltrates in 33 cancer types. In LGG, the most significant positive correlation (rho > 0.3, p-value < 2.01e-12) was with monocyte, NK, and B cells (Figure 15), whereas the negative correlation (rho ≤ 0.3, p-value < 2.26E-12) were with T cell CD4+ Th2, macrophage M1 and M2, myeloid dendritic cell, neutrophil, and activated myeloid dendritic cell activated cell infiltrates (Figure 16) (Table S26).

**FIGURE 15.**
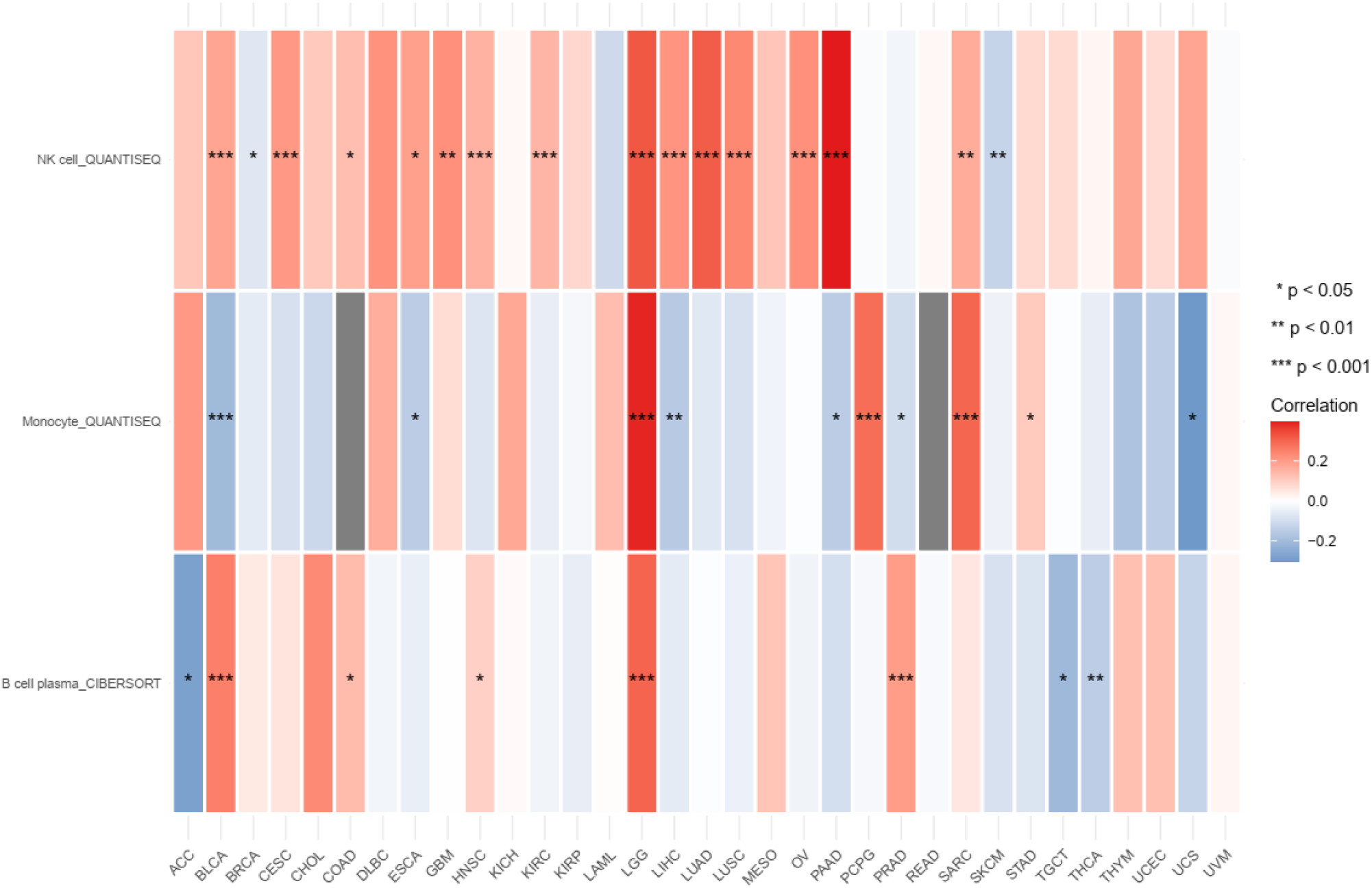
Heatmap of the correlations between the expression profiles of the 4-component gene signature and immune cell infiltrates. The red-blue heatmap scale shows the Spearman correlation coefficients’ significance levels (***, **, *). In LGG, the 4-component gene signature is positively correlated with monocyte, NK, and B cell infiltrates (rho > 0.3, p-value < 2.01e-12).

**FIGURE 16.**
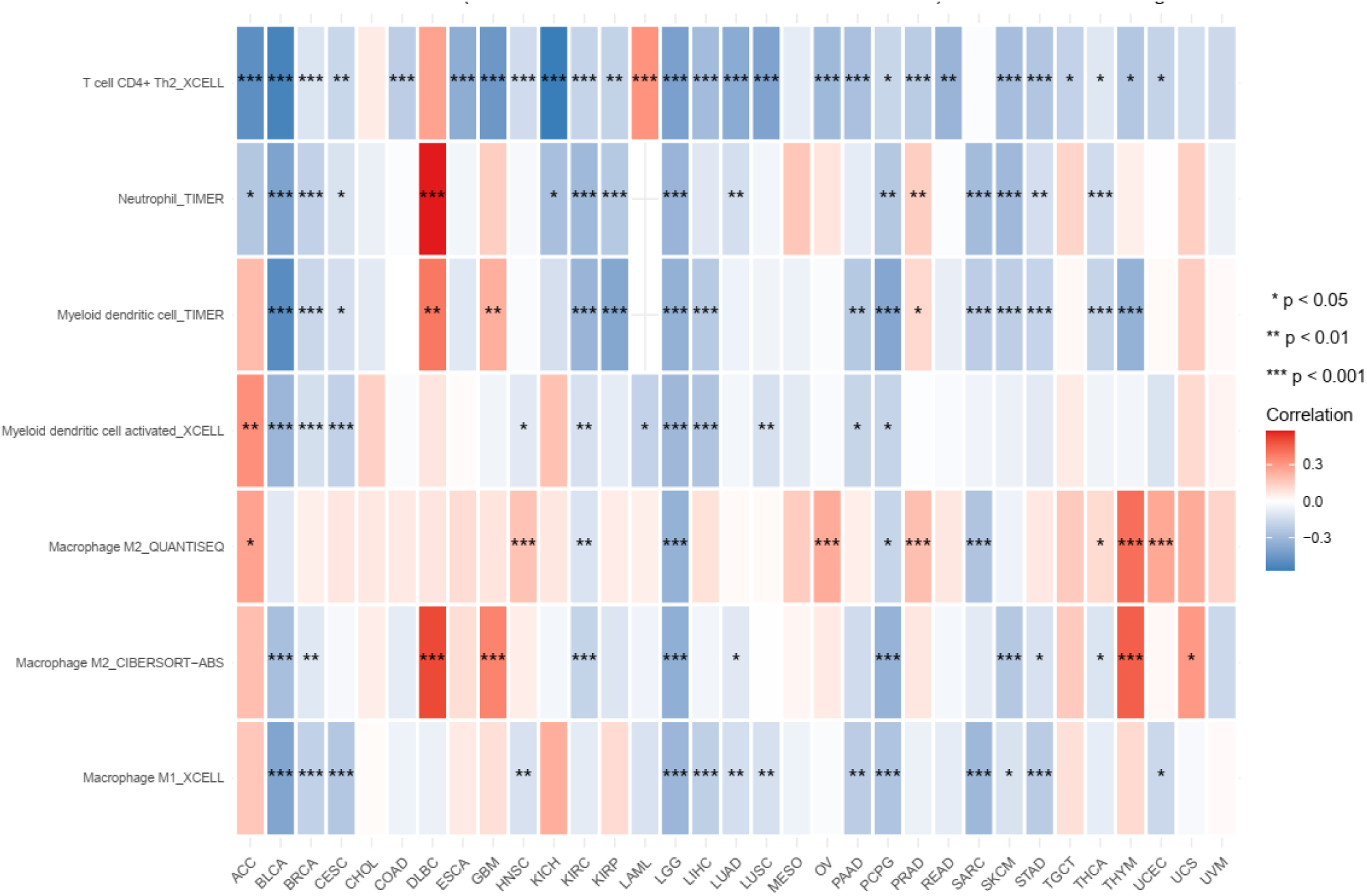
Heatmap of the correlations between the expression profiles of the 4-component gene signature and immune cell infiltrates. The red-blue heatmap scale shows the Spearman correlation coefficients’ significance levels (***, **, *). In LGG, the 4-component gene signature is negatively correlated with T cell CD4+ Th2, macrophage M1 and M2, myeloid dendritic cell, neutrophil, and activated myeloid dendritic cell activated cell infiltrates (rho ≤ 0.3, p-value < 2.26E-12).

### Correlation Analysis between Molecular Expression Profiles and Stemness

The molecular expression profiles of the 13 mitochondrial protein-coding genes exhibited positive (rho ranging 0.08-0.55, p < 0.045 in 27 cancer types) and negative correlations (rho ranging -0.51 to -0.06, p < 0.046 in 24 cancer types) with stemness (Table S27). The most extended signatures with 13 and 12 components positively correlated with stemness in KICH (rho = 0.50 p-value = 0) and LAML (rho =0.28 p-value = 0), respectively (Figure 17) (Table S18). The 13-component signature negatively correlated with stemness in 19 cancer types, with the most significant inverse coefficient in uterine corpus endometrial carcinoma, UCEC (rho = -0.41, p =0).

**FIGURE 17.**
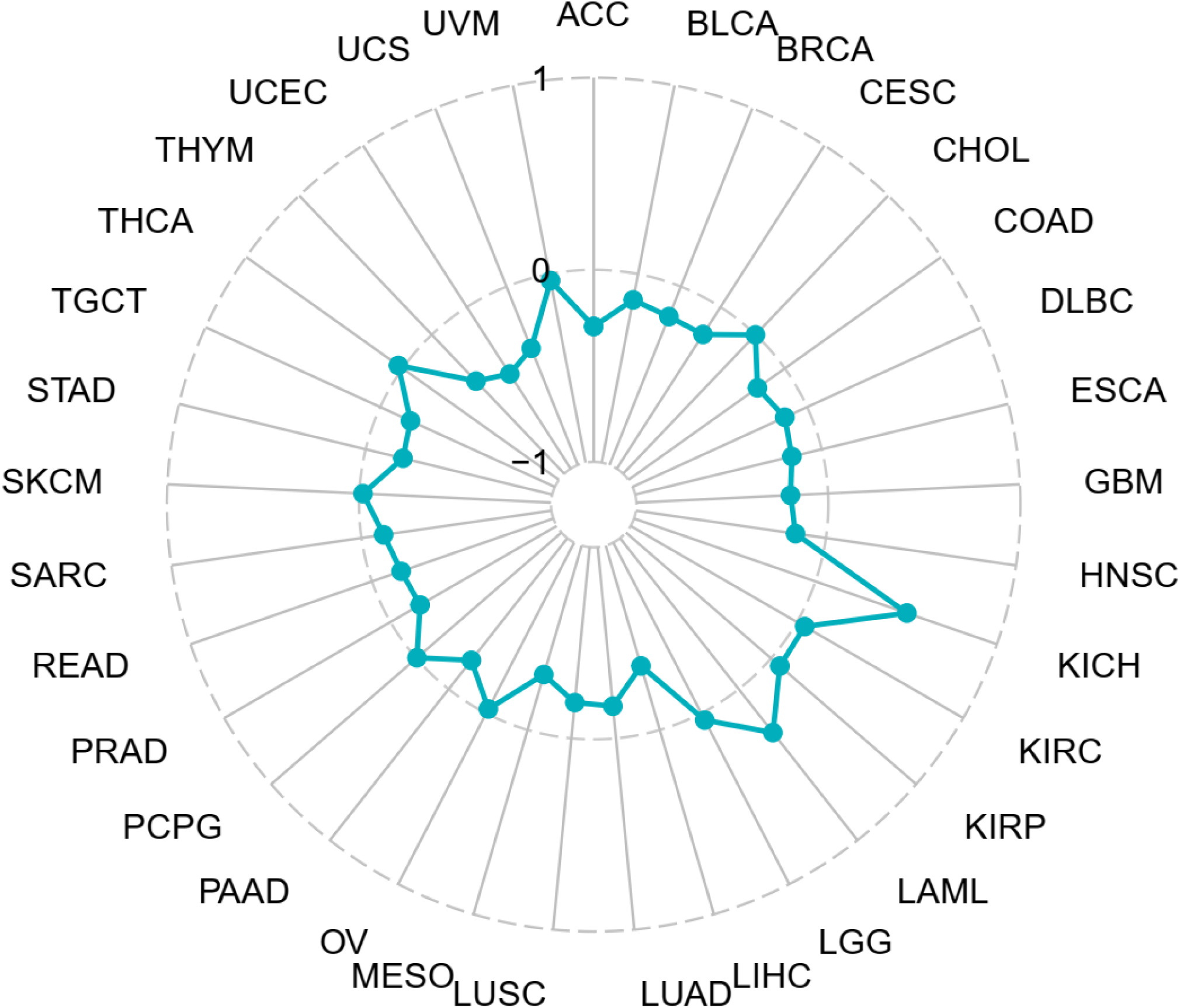
Radar plot for the correlation coefficients between the 13-component gene signatures and stemness in 33 cancer types. Shown is the distribution of the breadth of the Spearman correlation coefficients, where the association can be positive (+1), negative (−1), or neutral (0). The more significant positive correlation with stemness occurred in KICH and LAML.

## DISCUSSION

Here, we showed differential RNA expression of the 13 mitochondrial protein-coding genes in 33 cancer types and normal tissues, where eleven genes (*MT-ND2*, *MT-ND1*, *MT-ATP8*, *MT-ATP6*, *MT-CO2*, *MT-CYB*, *MT-CO3*, *MT-ND4L*, *MT-ND4*, *MT-ND3*, *MT-CO1*) exhibited an under-expression profile in brain cancer. Moreover, the 13 mitochondrial protein-coding genes also demonstrated a sex effect in expression profiles in different tissues. A male-biased expression across ages occurred in *MT-ND5* for Artery – Aorta, Brain – Substantia Nigra, and Esophagus – Mucosa (Wilcox test, p ≤0.001), and in the *MT-ND6* for Brain – Hypothalamus and Kidney – Cortex (Wilcox test, p ≤0.001). A female-biased across ages was also noticed in *MT-ATP6*, *MT-ATP8*, and *MT-CO1* for Artery – Coronary; *MT-CO1*, *MT-CO3*, and *MT-CYB* for Adipose – Visceral (Omentum); *MT-ND2* in Brain – Frontal Cortex (BA9) and Brain – Substantia Nigra; *MT-ND1* for Brain – Substantia Nigra. Interestingly, the *MT-ND5* also exhibits a biased expression across ages in Artery – Aorta, Brain – Frontal Cortex (BA9), and Colon – Transverse, but for females across different age groups. The same was noticed for *MT-ND6* in Brain – Frontal Cortex (BA9) and Brain – Substantia Nigra. The mitochondrial protein-coding genes exhibited a possible aging effect in expression profiles for the different age groups (20-49-year-old, 50-59-year-old, and 60-79-year-old) in different tissues, where the expression was higher in the 60-79-year-old group. In this group, we observed three genes (*MT-CO1*, *MT-ND5*, and *MT-ND6*, *MT-ND3*; Wilcox test, p ≤0.01) with the most significant expression profile in different tissues (Artery – Coronary, Brain – Amygdala, Colon – Sigmoid, Esophagus – Muscularis, Minor Salivary Gland, and Small Intestine – Terminal Ileum). Furthermore, a possible age-group-dependent sex-biased effect was observed across the 52 tissues analyzed for the 13 mitochondrial protein-coding genes and polymorphic site expression. A female age-group-dependent sex-biased effect was noticed for ten different tissues between the three age groups (20-49-year-old, 50-59-year-old, and 60-79-year-old; Tukey test, p ≤0.05), and a male age-group-dependent sex effects was observed for seven different tissues also between the three age groups (Tukey test, p ≤0.05). Thus, the polymorphic site expression of 13 mitochondrial protein-coding genes exhibited a sex-age-biased between the three age groups. The difference was higher in the last group (60-79-year-old).

The RNA profiles of the 13 mitochondrial protein-coding genes also exhibited a 4-component gene signature (*MT-CO1*, *MT-CO2*, *MT-ND5*, and *MT-ND6*) with under-expression in LGG (hazard ratio range 0.41-0.51, Logrank p ≤0.00026, p(HR) ≤0.00033). This signature was associated with survival outcomes in LGG for different categories, where the overexpression was associated with OS, DSS, and PFI (Log-rank p ≤ 0.0076). The 4-component gene signature was not affected by sex, OS, DSS, or PFI. Only for DFI outcome, we observed a sex-specific association in female patients, with a greater prognostic factor value (Log-rank p= 0.019) compared to males (Log-rank p= 0,14). The 13 mitochondrial protein-coding genes demonstrated a protective effect for the 4-component gene signature in LGG (hazard ratio -0.17) and a risk effect on THYM – hazard ratio 0.25 and UCEC – hazard ratio 0.06. We noticed different component gene signatures for the 13 mitochondrial protein-coding genes in the immune cell infiltrates. The most extended, 13-component gene signature occurred in DBLC, positively correlated with neutrophil, NK, T-cell CD4 memory, M1 and M2 macrophages, T-cell CD8+/CD4+, and B-cell memory cells. Moreover, the 4-component gene signature’s most significant positive correlation in LGG (rho > 0.3, p-value < 2.01e-12) was with monocyte, NK, and B cells, whereas the negative correlation (rho ≤ 0.3, p-value < 2.26E-12) were with T cell CD4+ Th2, macrophage M1 and M2, myeloid dendritic cell, neutrophil, and activated myeloid dendritic cell activated cell infiltrates. The molecular expression profiles of the 13 mitochondrial protein-coding genes exhibited positive and negative correlations in 24 cancer types with stemness. The most extended signatures with 13 and 12 components were positively correlated with stemness in KICH (rho = 0.50 p-value = 0) and LAML (rho =0.28 p-value = 0), respectively. The 13-component signature negatively correlated with stemness in 19 cancer types, with the most significant inverse coefficient in UCEC (rho = -0.41, p =0). The differentially expressed profiles of the 13 mitochondrial protein-coding genes in cancer have prognostic factor values in clinical studies. The correlation between the differentially expressed gene signatures and tumor stemness in LGG and immune cell infiltrates suggests that the reported signatures are proxies of oncogenic dedifferentiation.

Genome-wide screening of normal versus cancer transcription provides knowledge about the molecular information of cancer progression. It enables identifying transcriptional changes in differentially expressed genes (DEGs) to determine their prognostic role in clinical outcomes. The gene-by-gene analysis studies in most of the 13 mitochondrial protein-coding genes demonstrated that their RNA expression is significantly suppressed in several cancer types, such as breast, colon, kidney, and liver, the exception being KICH, with all genes overexpressed (Reznik et al., 2017;Kim et al., 2020). However, these studies considered the full range of logFC values and significant FDR values. In our analysis, we used a more restrictive method accepting only magnitudes of differential expression logFC values (≥ 1.5 or ≤ - 1.5) with genome-wide FDR values ≤ 5e-5. Thus, our restrictive methods allowed the identification of eleven genes downregulated in brain cancer (*MT-ATP6*, *MT-ATP8*, *MT-CO1*, *MT-CO2*, *MT-CO3*, *MT-CYB*, *MT-ND1*, *MT-ND2*, *MT-ND3*, *MT-ND4*, *MT-ND4L*), six in liver cancer (*MT-ATP6*, *MT-ATP8*, *MT-CYB*, *MT-ND2*, *MT-ND4*, *MT-ND4L)* and four in breast cancer (*MT-ATP8*, *MT-CYB*, *MT-ND1*, *MT-ND2*). In contrast, overexpression happens in AML (*MT-ND4*, *MT-ND5*, *MT-CO1*, *MT-ND4L*) and ovary cancer (*MT-ND6*, *MT-ND1*, *MT-CO1*).

The sex differences in gene expression have already been observed in different tissues, but nowadays, the mechanism behind the tissue-specific sex differences in gene expression is not understood (Rinn and Snyder, 2005;Ellegren and Parsch, 2007;Parsch and Ellegren, 2013). Thus, the sexual dimorphism in gene expression is still hard to describe, partly because the sex-related differences in the autosomal gene expression occur in various tissues (Mele et al., 2015). Additionally, gene expression during aging has already been studied between the sexes for several tissues (Tower, 2017). However, there are few references in the literature regarding the effects of aging on the expression of different tissues. A few reports demonstrated a low level of gene expression in older tissues, such as muscle, epicardial adipose tissue, and brain (Welle et al., 2003;Berchtold et al., 2008;Iacobellis, 2021). Thus, we wished to investigate whether there are sex-specific differences in the expression of the 13 mitochondrial protein-coding genes and if there is a difference in the expression of these genes between the tissues at different ages. Here, in males, two genes exhibited a male-biased expression in different tissues (*MT-ND5* for Artery – Aorta, Brain – Substantia Nigra, and Esophagus – Mucosa, and in the *MT-ND6* for Brain – Hypothalamus and Kidney – Cortex). Other nine genes showed a female-biased expression in different tissues *(MT-ATP6*, *MT-ATP8*, and *MT-CO1* for Artery – Coronary, *MT-CO1*, *MT-CO3* and *MT-CYB* for Adipose – Visceral (Omentum); *MT-ND2* for Brain – Frontal Cortex (BA9) and Brain – Substantia Nigra; *MT-ND1* for Brain – Substantia Nigra). *MT-ND5* also exhibits a biased expression in Artery – Aorta for females and in other tissues (Brain – Frontal Cortex (BA9) and Colon – Transverse). In agreement, Tower has reported that gene expression during aging has already been studied between the sexes for several tissues, such liver, which exhibits high sex-specific tissue expression with changes during aging (Tower, 2017).

Considering the evidence of a differential expression with age in coding genes (Frenk and Houseley, 2018), we decided to investigate possible differences in the expression of our mitochondrial protein-coding genes between 52 different tissues in the three age groups. Thus, we observe thirty-four tissues (n=34) have significantly different expression profiles (Wilcox test, p ≤0.05) in 20-49-year-old and 60-79-year-old groups for the 13 mitochondrial protein-coding genes. However, when we look more specifically at the tissue-specific expression for the different ages, we observe significant differences in different tissues’ expression. The artery – Coronary tissue exhibited more significant expression in the 20-49-year-old group (Wilcox test, p ≤0.001) than other tissues in this group. For the 50-59-year-old group (Wilcox test, p ≤0.01), the most significant differences in tissue expression occurred in the Brain – Cerebellar Hemisphere, Adipose – Visceral Omentum, and Bladder. This difference in expression of tissues also occurred in the group 60-79-year-old for Colon -Sigmoid, Artery – Coronary, Adipose – Visceral Omentum, and Minor Salivary Gland (Wilcox test, p ≤0.05). In accordance, Frenk and colleagues discuss that the aging effect in gene expression may occur for reasons not necessarily equivalent to programmed aging. Thus, gene expression changes may be reactive to physiological or cellular changes (Frenk and Houseley, 2018). Other studies also demonstrated that the most association of aging transcriptome is a consistent change of mitochondrial protein mRNAs, which was also observed in humans (de Magalhaes et al., 2009;van den Akker et al., 2014;Peters et al., 2015). No other evidence was found about the age-biased or sex-biased effect in humans’ 13 mitochondrial protein-coding genes.

Although most aging-related changes are correlated with declining mitochondrial function and ROS overproduction, oxidative damage, and accumulation of mtDNA mutations in somatic tissues, it’s necessary to emphasize the relationship between aging and defects of a wide range of proteins encoded by mtDNA (Poyton and McEwen, 1996;Scarpulla, 2008). In this context, we investigated the sex-dependent polymorphic site differences for mitochondrial protein-coding genes in different tissues. In this way, we observe a possible age-group-dependent sex effects expression in ten different tissues for females and seven for males between the three tested age groups. In agreement, another study reports evidence that aging occurs differently among males and females as expected from the phenotypic differences associated with aging (Mele et al., 2015;Kim et al., 2016;Gershoni and Pietrokovski, 2017;Naqvi et al., 2019). Other studies still report that breast tissue has the most sexual differentiation (Mele et al., 2015;Gershoni and Pietrokovski, 2017;Lopes-Ramos et al., 2020), but interestingly, in our study, we notice a sex-biased gene expression for breast tissue related to age groups. Thus, human aging phenotypes may be not only sex-specific but, in a more complex way, age-group-dependent.

In the last ten years, the sequencing and bioinformatic approaches expanded the possibility of accessing and handling gene signature data. This improves the analysis of survival outcomes for patients with several cancers (Rao et al., 2011;Gao et al., 2019;Li et al., 2019;Zhang et al., 2020;Pawar et al., 2022). Thus, investigating gene signature expression from mitochondrial protein-coding genes related to cancer survival outcomes was possible, which led us to perform additional analysis of these approaches in this study. In this context, Schopf and colleagues performed an analysis of gene-expression signatures of tumor sample transcriptomes that allowed the identification of several tumors associated with a shorter survival time (Schopf et al., 2020). Here we investigate whether 13 mitochondrial protein-coding genes exhibit a gene signature for survival outcomes in 33 different cancer types across donors of the TCGA cohort and whether the sex-specific differences are risk factors in cancer. We identified a novel 4-component mitochondrial protein-coding gene signature (*MT-CO1*, *MT-CO2*, *MT-ND5*, *MT-ND6*) whose molecular expression profile has a prognostic factor value for survival outcomes in LGG. Xiao and colleagues demonstrate that other gene signatures, such as CD44-related genes, with high-risk expression in LGG, are associated with greater survival rates in OS and disease-free survival (Log-rank p˂ 0.0001) (Xiao et al., 2020b). Here, we observe the 4-component gene signature exhibiting an over-expression profile in OS and DSS for LGG with elevated survival rates in the high-risk group (Log-rank p < 0.0001). In contrast, Xiao and colleagues showed that patients in the high-risk group had significantly poorer survival results (Log-rank p= 0.000012) in not mitochondrial gene signatures for LGG (Xiao et al., 2020a). Other studies have already demonstrated a better prognosis in survival results for the low-risk group using gene signatures (Log-rank p ˂0.0001) (Chen et al., 2021;Guo et al., 2021). Additionally, the low-risk group related to the 4-component gene signature exhibits a poor prognosis compared to the high-risk group in OS, DSS, and PFI survival outcomes.

In a study restricted to lung squamous cell carcinoma (LUSC) and lung adenocarcinoma (LUAD) (Li et al., 2018), the two major subtypes of lung cancer, the 13 mitochondrial respiratory genes were reported to be downregulated in tumor tissues compared with matched control tissues. The under-expression of *MT-ND5* or *MT-ND6* genes was associated with overall survival outcomes and tumor progression in LUAD and LUSC. Our gene-by-gene analysis confirmed the above associations with overall survival outcomes in LUAD (*MT-ND6* Log-rank test statistics = 10.04, p-value = 0.001529; *MT-ND5* Log-rank = 4.186, p-value = 0.04076), when the samples were clustered into two expression groups (Figures S10-S11), as well in LUSC, both *MT-ND6* and *MT-ND5* (Figure S18 and S25) was not significant for the overall survival patients (*MT-ND6* Log-rank p-value = 0.4; *MT-ND5* Log-rank p-value = 0.19). However, we extended the association with three other survival outcomes: DSS, DFI, and PFI (Figure S10-S25).

Identifying gene signatures as risk effect factors may be relevant for proposing personalized treatment of several cancers, such as hepatocellular carcinoma, colon cancer, and gliomas (Li et al., 2020;Liu and Li, 2021;Zhao et al., 2021). We aimed to identify a correlation between protective or risk effects associated with survival outcome for the 13 mitochondrial protein-coding genes in 33 different cancer types by using the TCGA cohort. Concerning the protective effect associated with survival outcome in this study, we found and 4-component gene signature (*MT-CO1*, *MT-CO2*, *MT-ND5*, *MT-ND6*) with the association in six different cancer types (ACC, KICH, KIRP, LIHC, LGG, PAAD; hazard ratio ranging -0.08 to -032). Regarding the risk effects, two tissues exhibited associations (THYM and UCEC; hazard ratio ranging from 0.06 to 0.25). To LGG, Zhang and colleagues showed a protective/risk effect using gene signatures (non-mitochondrial) correlation (Zhang et al., 2019). Accordingly, autophagy gene signature also exhibited a risk effect factor for LGG patients (Lin and Lin, 2021). Equally, Wang and colleagues demonstrate evidence of risk signatures genes (Wang et al., 2022). In the same way, other gene signatures, extracellular matrix related, exhibited a risk effect for gliomas in the TCGA cohort but also demonstrated an association with immune infiltrates (Liu and Li, 2021). In contrast, we observed a protective effect factor in LGG patients to four mitochondrial components gene signature, which is not shown in the literature now.

Studying gene signatures associated with immune infiltrates may provide a potential clinical implication for cancer treatment and prognosis (Iglesia et al., 2016;Yan et al., 2020). Thus, we use the 13 mitochondrial protein-coding genes in the context of gene signature to correlate with immune cell infiltrates in 33 cancer types from the TCGA cohort. Studies already show evidence about gene signatures, such as autophagy genes related, strongly associated with six immune cell infiltrates (macrophages M0 and M1, Neutrophils, T cells CD8, T cells follicular helper, T cells reg) in LGG (Quan et al., 2021). Additionally, other gene signatures, such as immune-related genes, demonstrate positive and negative correlations with several immune cell infiltrates in LGG (macrophages, dendritic cells, B cells, and CD4 T cells) (Pan et al., 2021). In the same way, our study showed positive and negative correlations with immune infiltrates in LGG, with monocyte, NK, and B cells, T cell CD4+, macrophage (M1 and M2), and myeloid dendritic cells.

In this way, our mitochondrial protein-coding gene signature positively correlates with M1 and M2 macrophages from immune cells infiltrated in LGG. Similarly, in DLBC the 13-component gene signature show a positive correlation for six cells including neutrophils. In agreement, M2 macrophage has been confirmed to promote immunosuppression and proliferation of LGG; on the other hand, T cells and natural killer (NK) cells can be associated with poorer OS and DSS in LGG, both in non-mitochondrial gene signature (Ye et al., 2021;Zhu et al., 2022). Concerning DLBC prognosis, the tumor-associated neutrophils show a poor prognostic, in the same way, Manfroi and colleagues demonstrated that in DLBC patients with a higher ratio of neutrophils had a worse prognosis, both in non-mitochondrial gene signature (Manfroi et al., 2018;Mu et al., 2018). In other studies, it is also possible to observe T-lymphocyte signatures impacting the prognosis of patients with DBLC, demonstrating the importance of survival gene signatures in aggressive B-cell lymphomas (Ansell et al., 2001;Keane et al., 2013;Keane et al., 2015).

Using gene signatures to study tumor stem cells has provided new insights about stemness for measuring the degree of oncogenic dedifferentiation. A correlation between stemness signatures and unfavorable outcome for some cancers, including gliomas, was observed in the literature for the TCGA cohort (Malta et al., 2018). In agreement, there is evidence of gene signatures, such as the 5-mRNA signature, that correlate with stemness in melanoma and hepatocellular carcinoma, which exhibit a negative correlation for these cancers (Cai et al., 2021;Zhang et al., 2021). Furthermore, cancer stemness has been demonstrated to be associated with kidney cancers, such as KIRC, but not for KICH, and negatively correlates with immune infiltrates (Xiao et al., 2021). Additionally, other studies demonstrated a stemness-related gene signature expression in cancers such as hepatocellular carcinoma, pancreatic ductal adenocarcinoma, and endometrial cancer as a novel prognostic marker for survival (Hong et al., 2021;Huang et al., 2021;Xu et al., 2021). In this study, we report a significant positive correlation between the expression profiles of the 13 mitochondrial gene signatures and stemness in KICH and LAML, as well as a negative correlation in THYM and UCEC. Thus, identifying gene signatures with prognostic models based on stemness provides a powerful tool for cancer treatment.

## CONCLUDING REMARKS

LGG downregulates mitochondrial protein-coding genes. The downregulated 4-component gene signature (*MT-CO1*, *MT-CO2*, *MT-ND5*, *MT-ND6*) exhibits good OS, DSS, and PFI predictive value, and DFI and PFI survival were female-biased. The expression of *MT-ND5* and *MT-ND6* is female-biased in brain tissues. Age-dependent sex bias occurs in brain tissues, with higher gene expression in the 50-59 and 60-79-year-old populations. Distinct immune cell infiltrates were associated with mitochondrial protein-coding gene signature in LGG. We found a stemness-related gene signature for all 13 mtDNA genes in KICH. This study identifies mitochondrial prognostic genes.

## WEB RESOURCES

The URLs for data presented herein are as follows:

UCSC Genome Browser, https://genome.ucsc.edu/

UCSCXenaShiny, https://cran.r-project.org/web/packages/UCSCXenaShiny/index.html

GTEx Portal, https://www.gtexportal.org/

GEPIA2, http://gepia2.cancer-pku.cn/#index

MITOMAP, https://mitomap.org/foswiki/bin/view/MITOMAP/WebHome

R software package, http://www.R-project.org

## SUPPLEMENTARY TABLES

Table S1. Mitochondrial Protein-Coding Genes Location and eSNVs

Table S2. Primary Tissues Available by GTEx

Table S3. Types of Cancers in the TCGA

Table S4. Differentially Expressed Genes for mtDNA protein-coding genes in all tissues from GTEx and TCGA

Table S5. Female Expression of mtDNA Protein-coding Genes Across Different Ages for each Tissue.

Table S6. Male Expression of mtDNA Protein-coding Genes Across Different Ages for each Tissue.

Table S7. Comparison of Mitochondrial Polymorphic Sites Expression by Gender in 20-49 years

Table S8. Comparison of Mitochondrial Polymorphic Sites Expression by Gender in 50-59 years

Table S9. Comparison of Mitochondrial Polymorphic Sites Expression by Gender in 60-79 years

Table S10. Survival Map of 13 Mitochondrial Protein-Coding Genes in Different Cancers

Table S11. Overall Survival Expression Profile of 4-component Gene Downregulated Signatures in LGG

Table S12. Disease-specific-survival Expression Profile of 4-component Gene Downregulated Signatures in LGG

Table S13. Progression-free-interval Expression Profile of 4-component Gene Downregulated Signatures in LGG

Table S14. Overall Survival Expression Profile of 4-component Gene Downregulated Signatures in LGG for Males

Table S15. Overall Survival Expression Profile of 4-component Gene Downregulated Signatures in LGG for Females

Table S16. Progression-free-interval Expression Profile of 4-component Gene Downregulated Signatures in LGG for Females

Table S17. Progression-free-interval Expression Profile of 4-component Gene Downregulated Signatures in LGG for Males

Table S18. Disease-specific-survival Expression Profile of 4-component Gene Downregulated Signatures in LGG for Females

Table S19. Disease-specific-survival Expression Profile of 4-component Gene Downregulated Signatures in LGG for Males

Table S20. Disease-free-interval Expression Profile of 4-component Gene Downregulated Signatures in LGG for Males

Table S21. Disease-free-interval Expression Profile of 4-component Gene Downregulated Signatures in LGG for Females

Table S22. Mitochondrial Protein-Coding Genes Profile for Protective or Risk Effect on Survival in Different Cancers

Table S23. Association Between Molecular Profile of 13 Mitochondrial Protein-Coding Genes and Immune Cells Infiltrate Signatures

Table S24. Correlations Between the Expression Profiles of the 7-component Gene Signature and Immune Cell infiltrates

Table S25. Correlations Between the Expression Profiles of the 8-component Gene Signature and Immune Cell infiltrates

Table S26. Correlations Between the Expression Profiles of the 4-component Gene Signature and Immune Cell infiltrates

Table S26. Correlations Between the Molecular Profile of 13 Mitochondrial Protein-Coding Genes and Stemness

## SUPPLEMENTARY FIGURES

**Supplementary Figure 1.** The landscape of Mitochondrial Protein-Coding Genes and Polymorphic Sites distribution Across all Samples and Tissues.

**Supplementary Figure 2.**
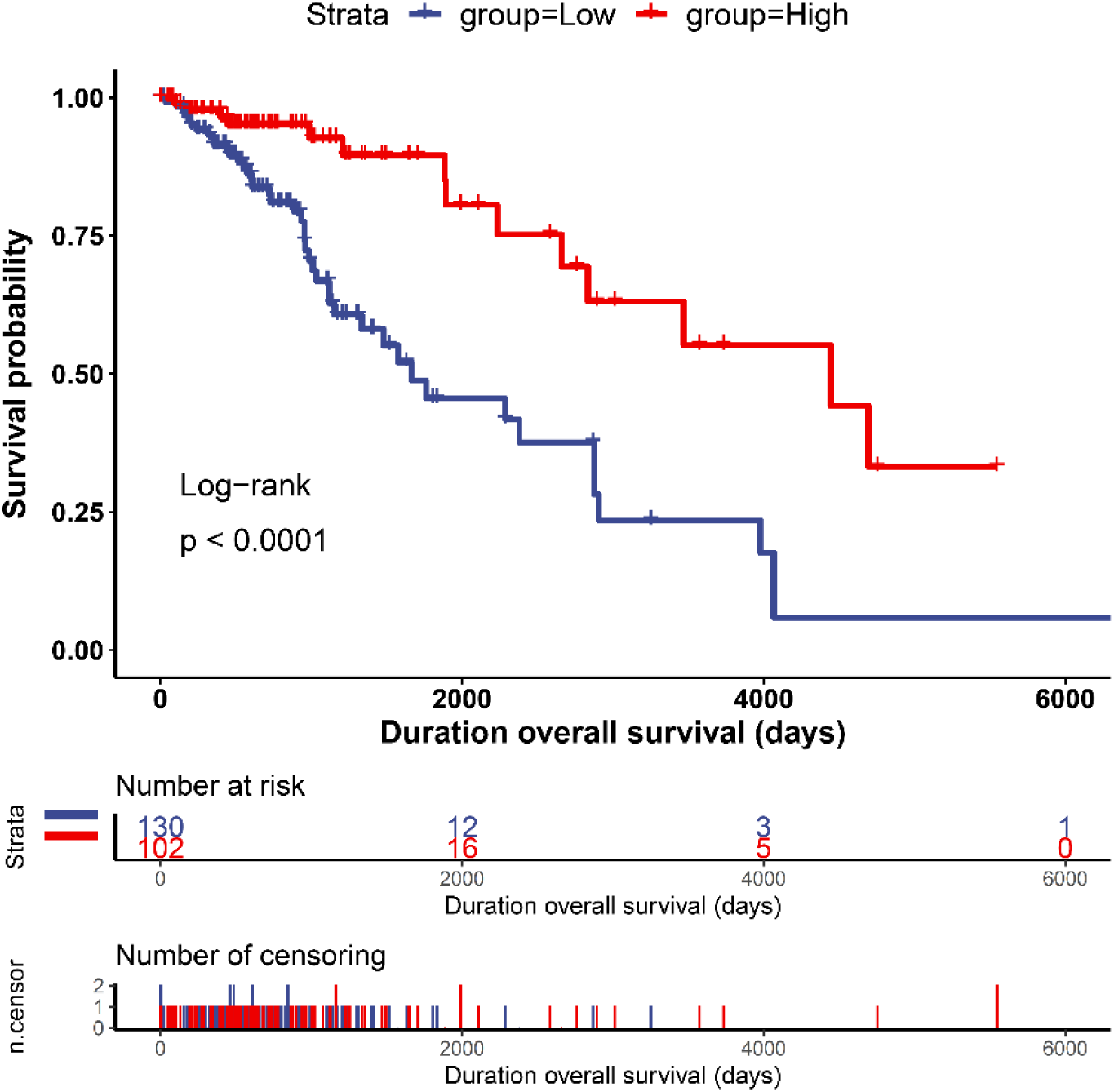
Kaplan-Meier plot of overall survival outcome in low-grade glioma. The curves denote the contribution of the 4-component gene signature (*MT-CO1*, *MT-CO2*, *MT-ND5*, *MT-ND6*) to the overall survival outcome in patients from the low (blue – downregulated) and high (red – upregulated) expression groups. The censoring number refers to patients who did not suffer the outcome of interest during the specified study period. The overexpression of the signature has a more significant prognostic factor value.

**Supplementary Figure 3.**
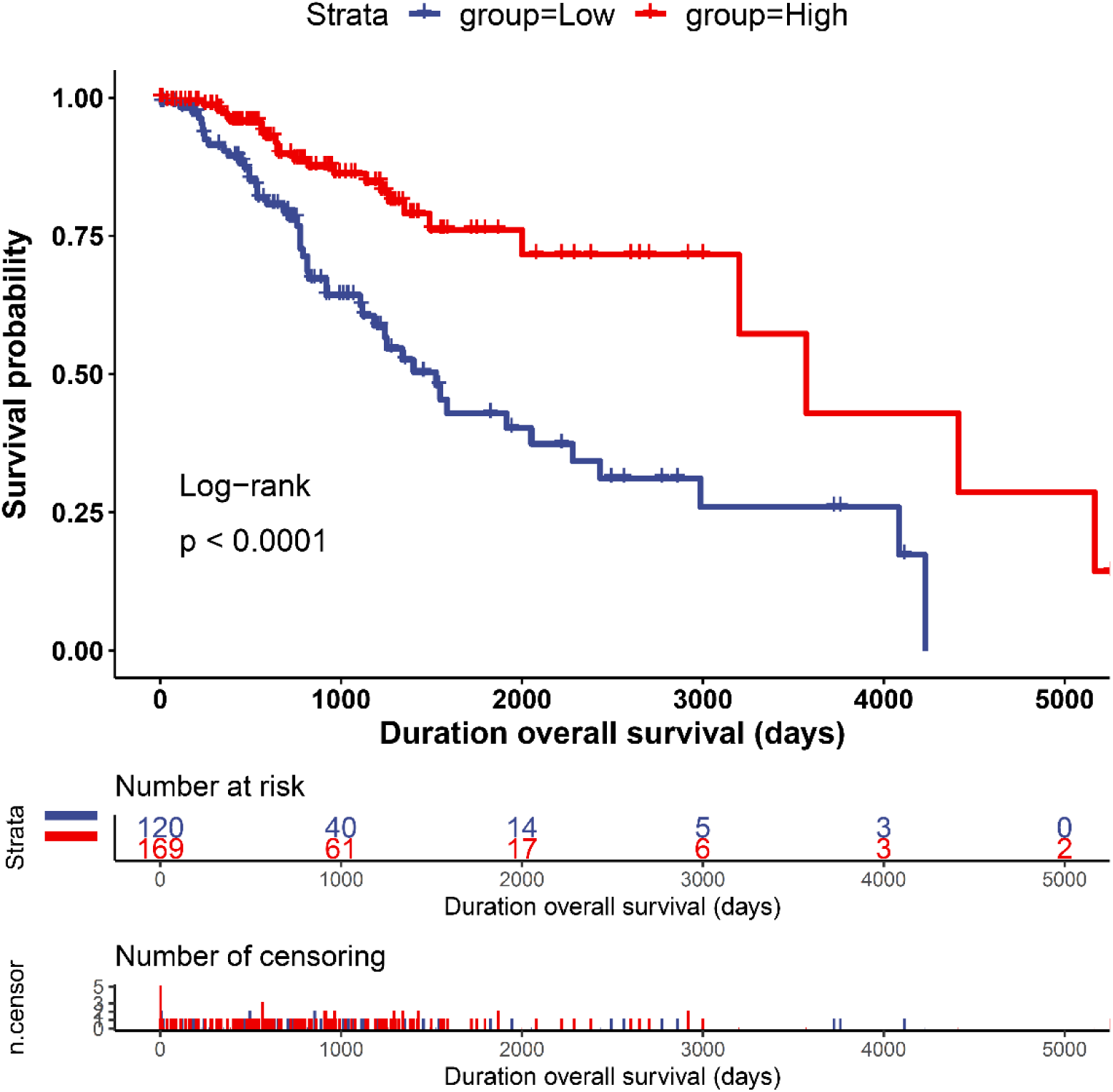
Kaplan-Meier plot of overall survival outcome in low-grade glioma. The curves denote the contribution of the 4-component gene signature (*MT-CO1*, *MT-CO2*, *MT-ND5*, *MT-ND6*), in males, to the overall survival outcome in patients from the low (blue – downregulated) and high (red – upregulated) expression groups. The censoring number refers to patients who did not suffer the outcome of interest during the specified study period. The overexpression of the signature has a more significant prognostic factor value.

**Supplementary Figure 4.**
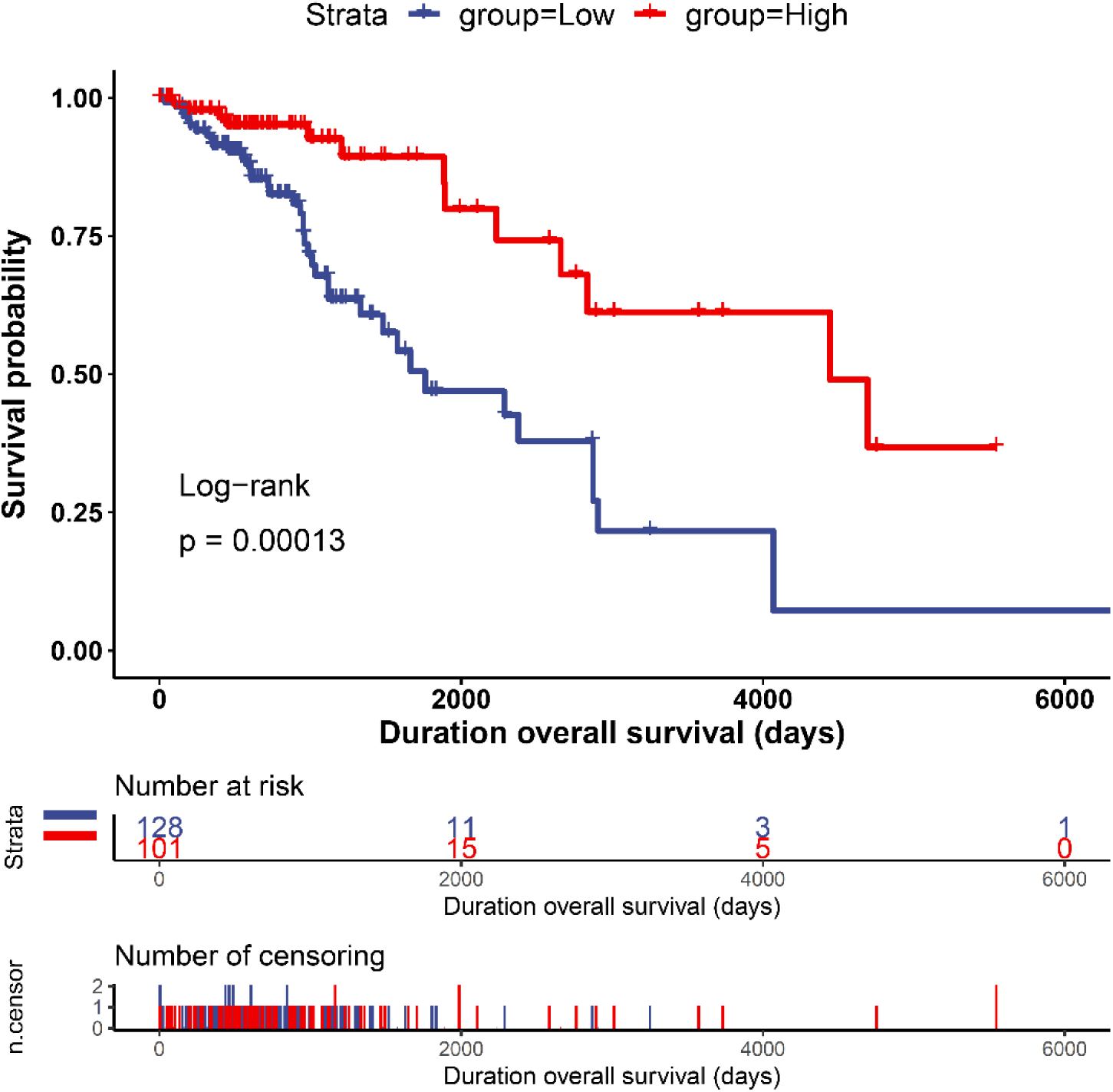
Kaplan-Meier plot of disease-specific survival outcome in low-grade glioma. The curves denote the contribution of the 4-component gene signature (*MT-CO1*, *MT-CO2*, *MT-ND5*, *MT-ND6*), in females, to the disease-specific survival outcome in patients from the low (blue – downregulated) and high (red – upregulated) expression groups. The censoring number refers to patients who did not suffer the outcome of interest during the specified study period. The overexpression of the signature has a more significant prognostic factor value.

**Supplementary Figure 5.**
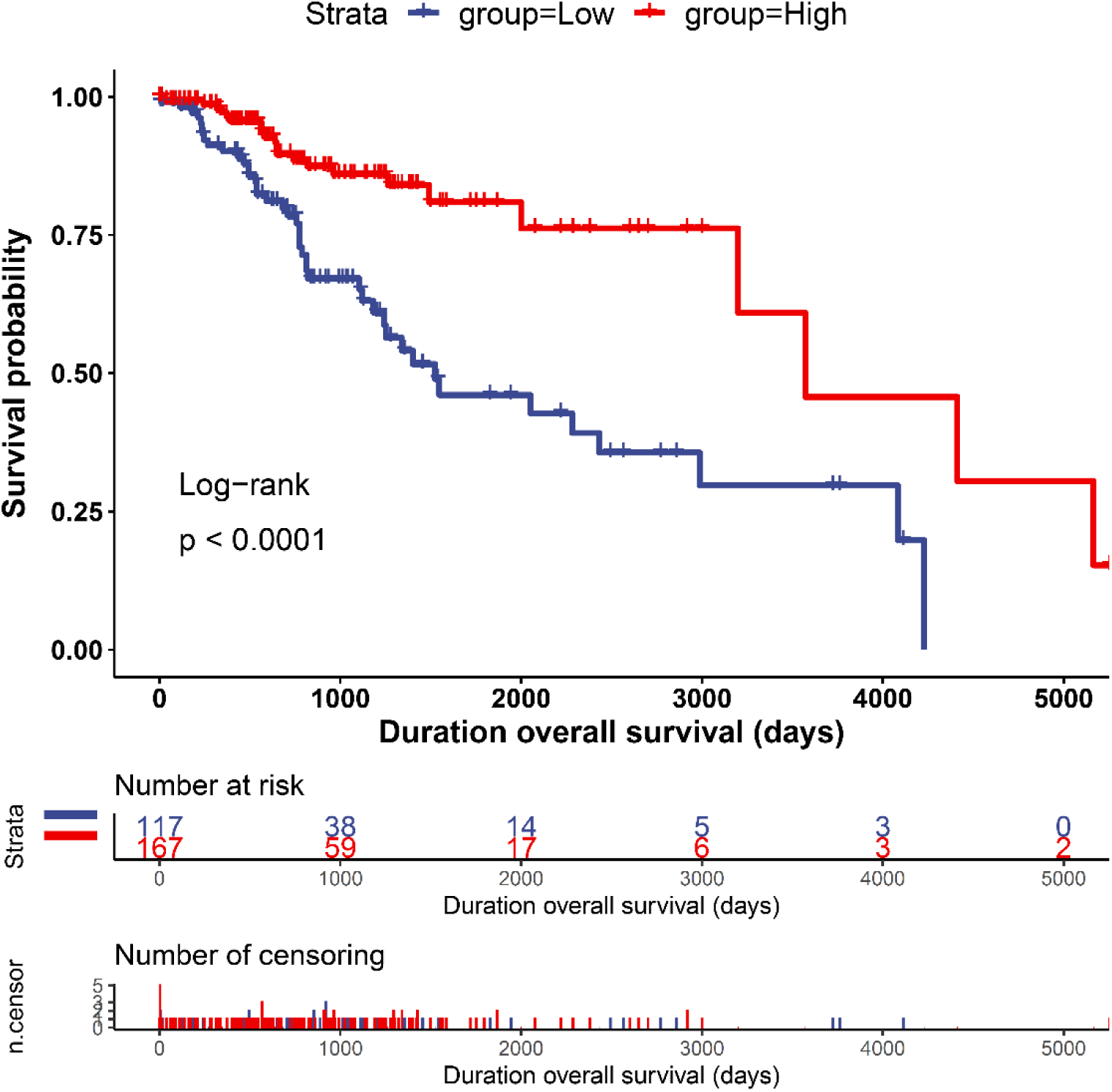
Kaplan-Meier plot of disease-specific survival outcome in low-grade glioma. The curves denote the contribution of the 4-component gene signature (*MT-CO1*, *MT-CO2*, *MT-ND5*, *MT-ND6*), in males, to the disease-specific survival outcome in patients from the low (blue – downregulated) and high (red – upregulated) expression groups. The censoring number refers to patients who did not suffer the outcome of interest during the specified study period. The overexpression of the signature has a more significant prognostic factor value.

**Supplementary Figure 6.**
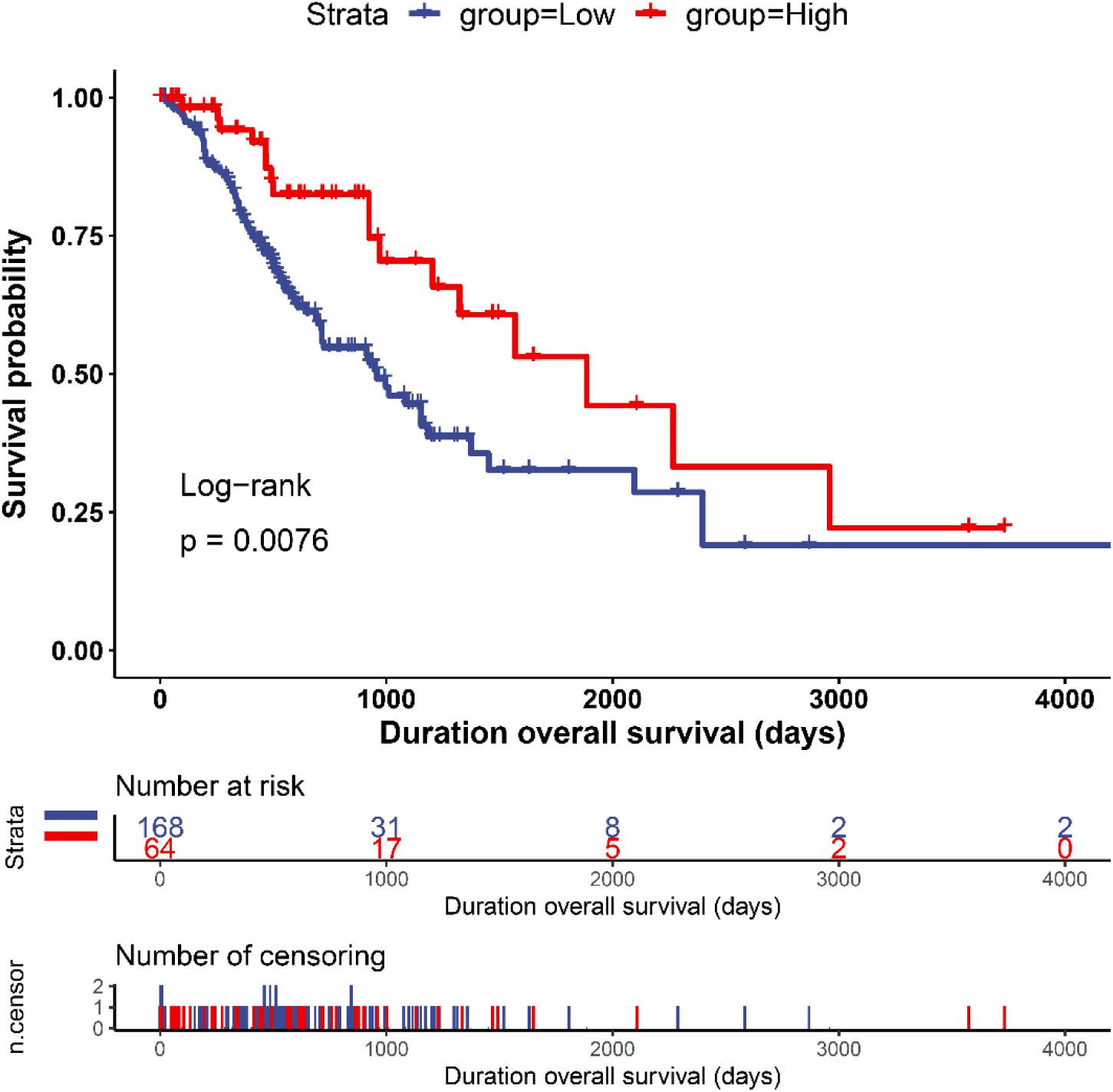
Kaplan-Meier plot of progression-free-interval outcome in low-grade glioma. The curves denote the contribution of the 4-component gene signature (*MT-CO1*, *MT-CO2*, *MT-ND5*, *MT-ND6*), in females, to the progression-free-interval outcome in patients from the low (blue – downregulated) and high (red – upregulated) expression groups. The censoring number refers to patients who did not suffer the outcome of interest during the specified study period. The overexpression of the signature has a more significant prognostic factor value.

**Supplementary Figure 7.**
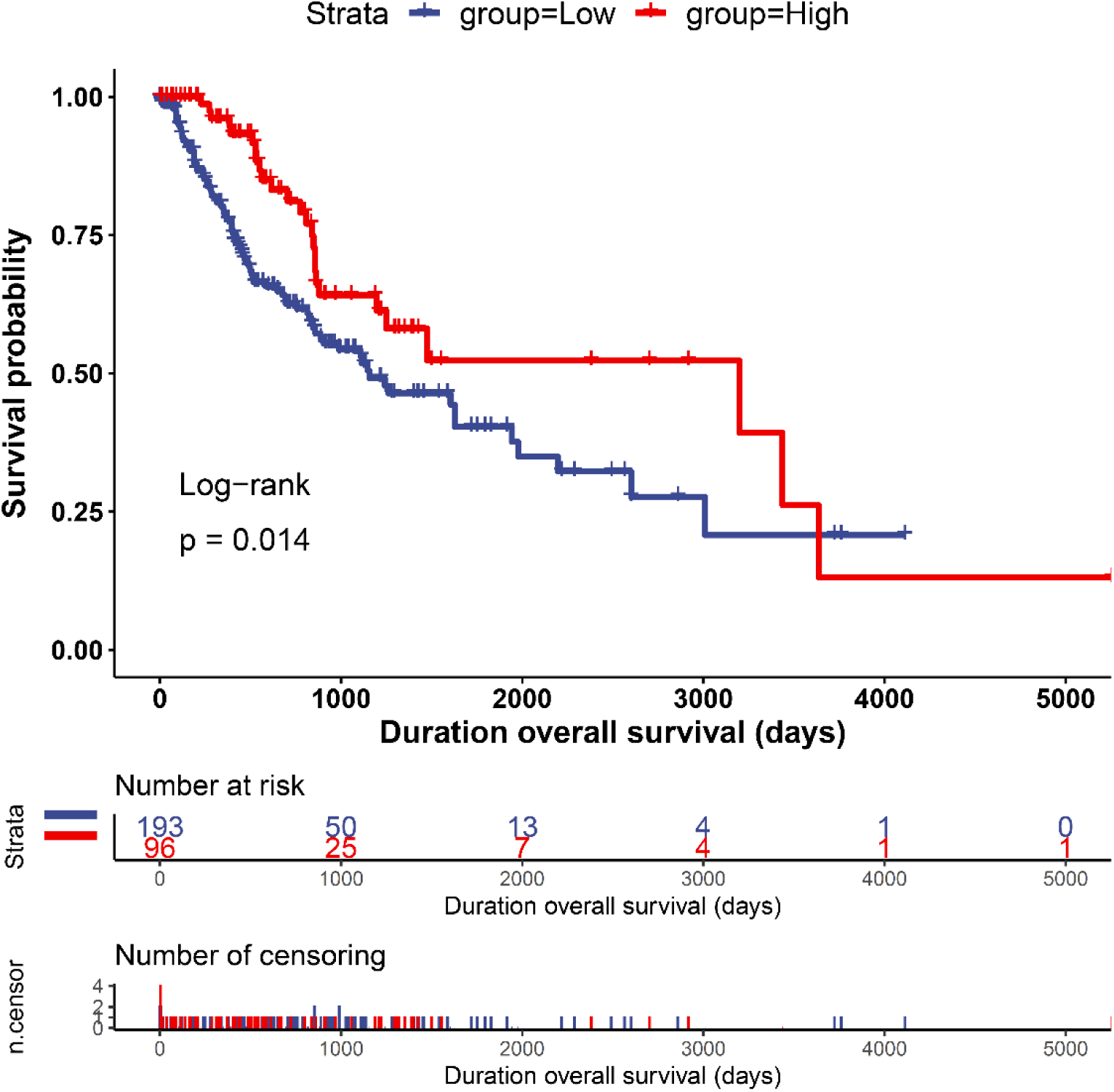
Kaplan-Meier plot of progression-free-interval outcome in low-grade glioma. The curves denote the contribution of the 4-component gene signature (*MT-CO1*, *MT-CO2*, *MT-ND5*, *MT-ND6*), in males, to the progression-free-interval outcome in patients from the low (blue – downregulated) and high (red – upregulated) expression groups. The censoring number refers to patients who did not suffer the outcome of interest during the specified study period. The overexpression of the signature has a more significant prognostic factor value.

**Supplementary Figure 8.**
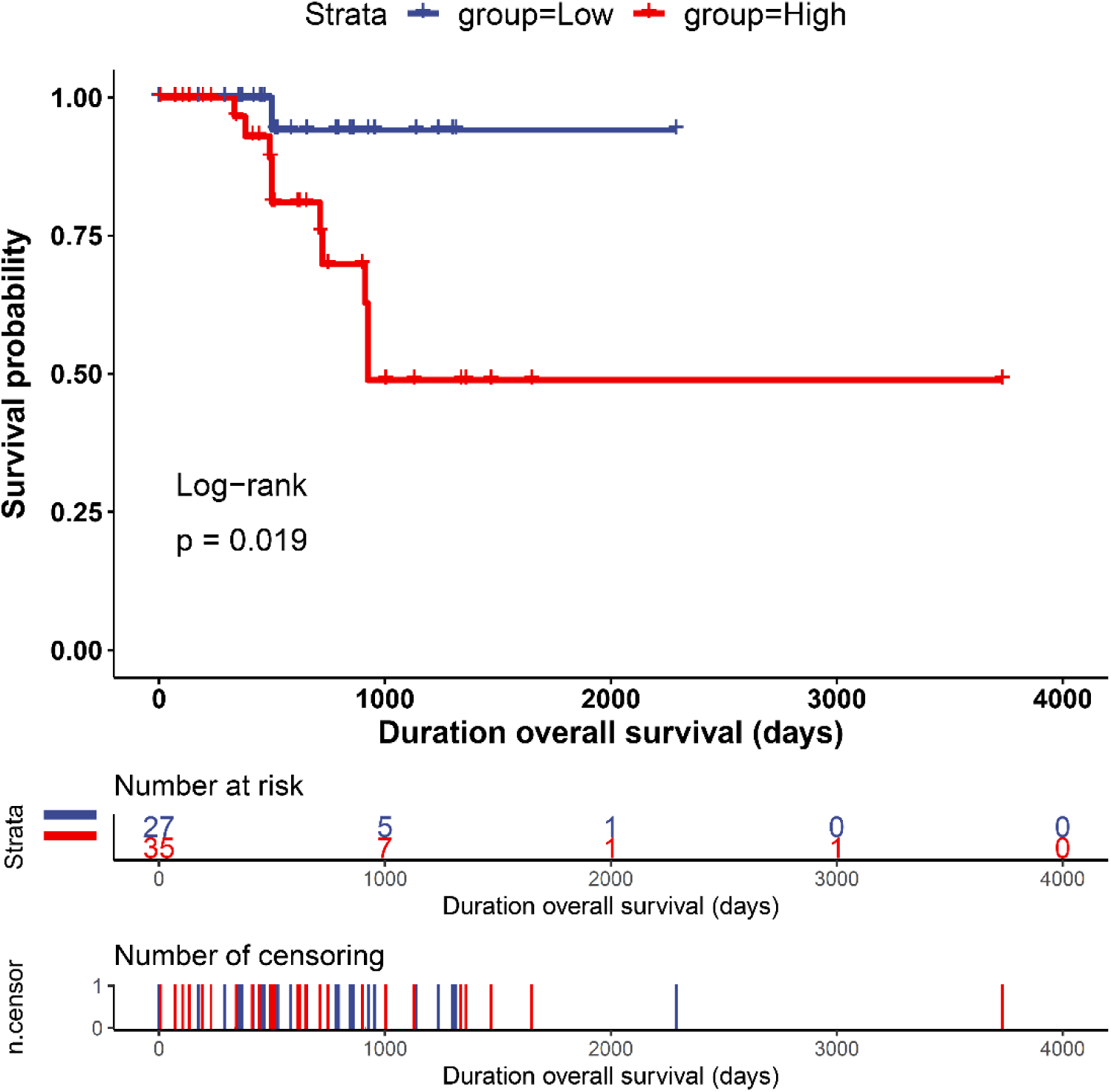
Kaplan-Meier plot of disease-free interval outcome in low-grade glioma in females. The curves denote the contribution of the 4-component gene signature (*MT-CO1*, *MT-CO2*, *MT-ND5*, *MT-ND6*) to the disease-free interval outcome in female patients from the low (blue – downregulated) and high (red – upregulated) expression groups. The censoring number refers to patients who did not suffer the outcome of interest during the specified study period, and the downregulation of the signature has a more significant prognostic factor value.

**Supplementary Figure 9.**
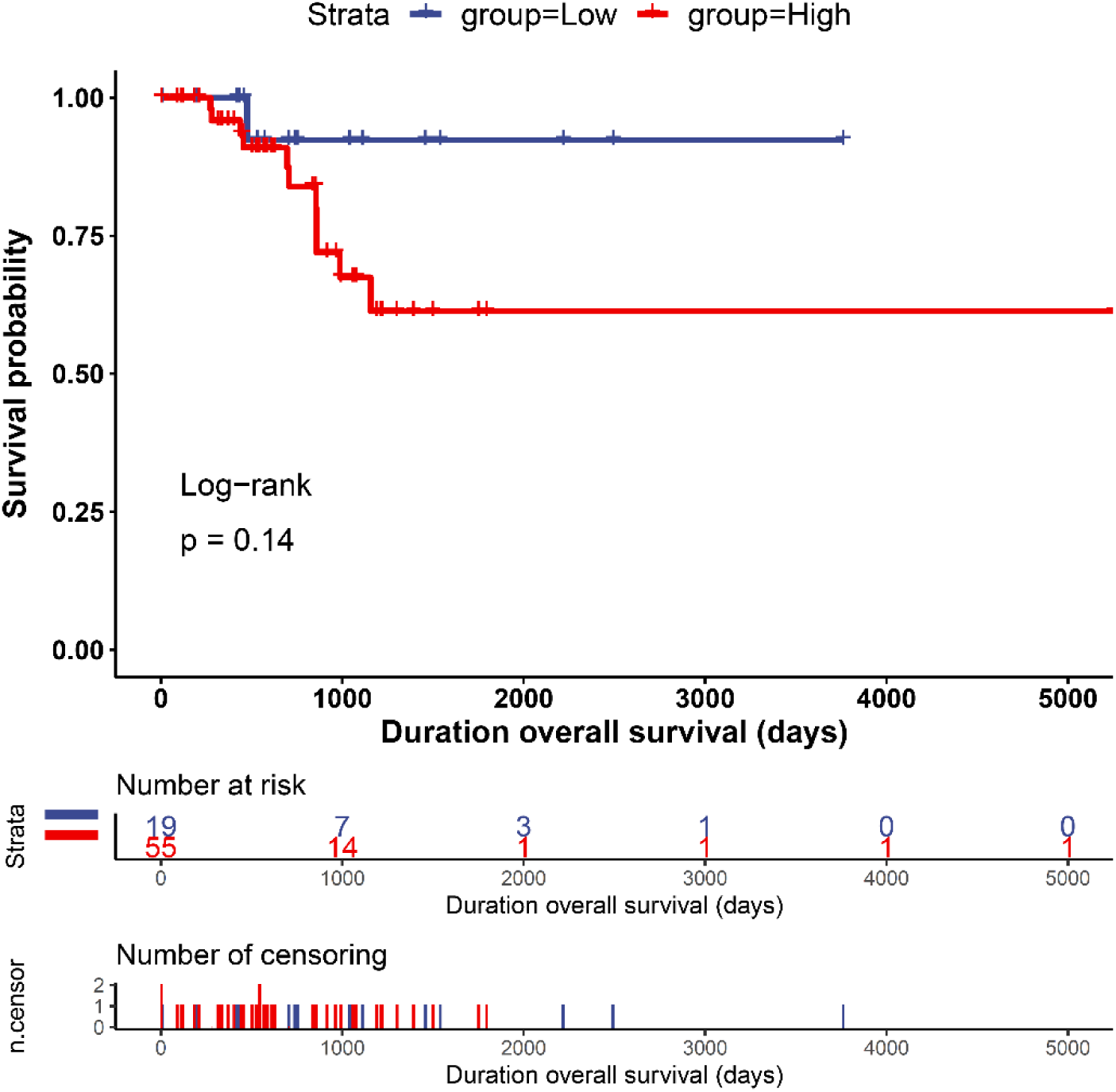
Kaplan-Meier plot of disease-free interval outcome in low-grade glioma. The curves denote the contribution of the 4-component gene signature (*MT-CO1*, *MT-CO2*, *MT-ND5*, *MT-ND6*), in males, to the disease-free interval outcome in patients from the low (blue – downregulated) and high (red – upregulated) expression groups. The censoring number refers to patients who did not suffer the outcome of interest during the specified study period. The overexpression of the signature has a more significant prognostic factor value.

**Supplementary Figure 10.**
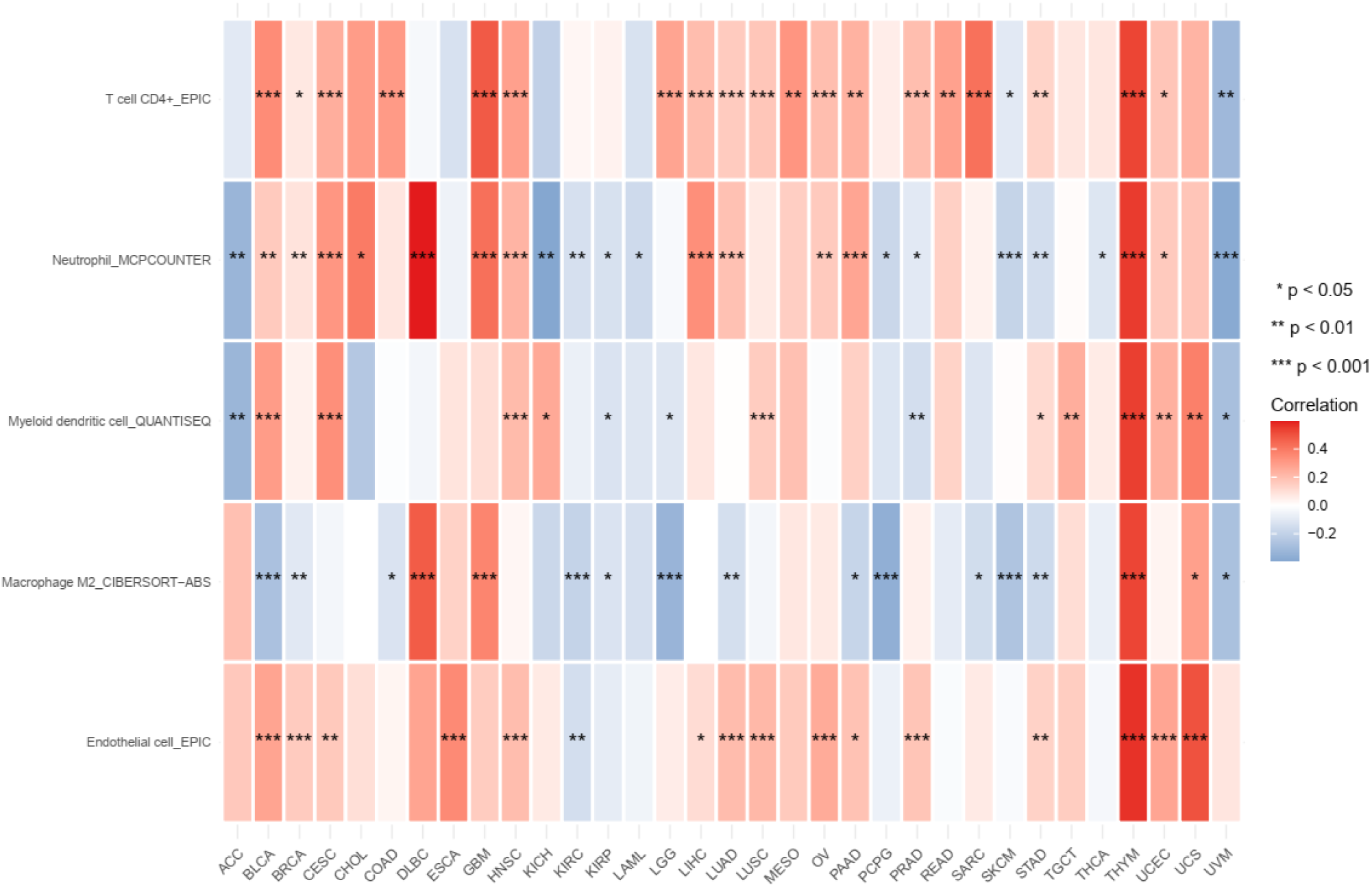
Heatmap of the correlations between the expression profiles of the 7-component gene signature and immune cell infiltrates. The red-blue heatmap scale shows the significance levels of Spearman correlation coefficients (***, **, *). In THYM, the 7-component gene signature is positively correlated with infiltrates of endothelial cells, neutrophils, myeloid dendritic cells, T cell CD4+, and macrophage M2 (rho ≥0.50, p-value ≤ 0.001).

**Supplementary Figure 11.**
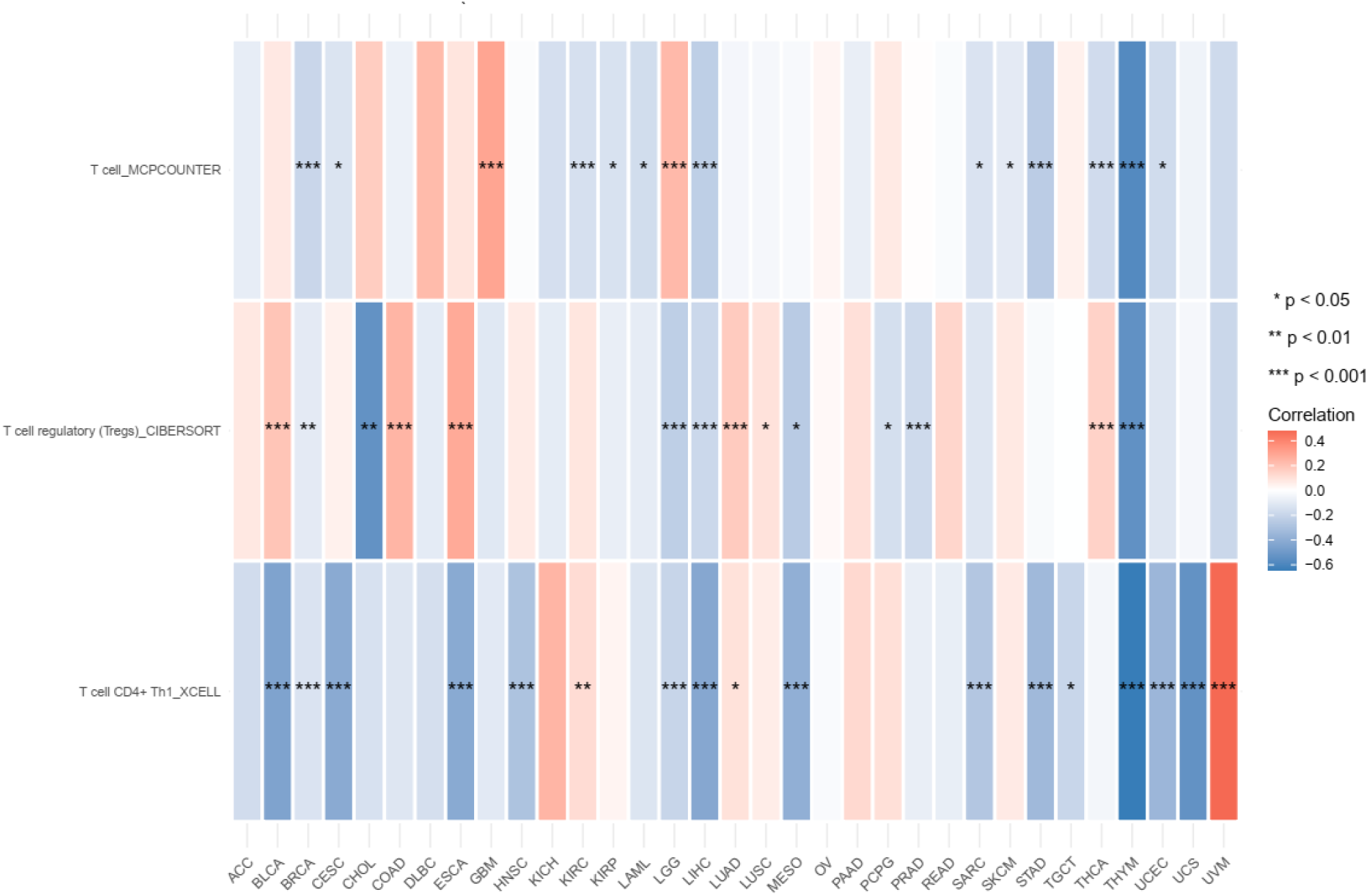
Heatmap of the correlations between the expression profiles of the 8-component gene signature and immune cell infiltrates. The red-blue heatmap scale shows the significance levels of Spearman correlation coefficients (***, **, *). In THYM, the 7-component gene signature is negatively correlated with infiltrates of T cell CD4+ Th1, T cell, and T cell regulatory (Tregs) (rho ≤ -0.5, p-value ≤ 3.05e-10).

## DRIVE ACCESS FOR SUPPLEMENTARY DATA

https://drive.google.com/drive/folders/1JggbN51c0xpeUNrDJugP4yCqersz_F0_?usp=sharing

## Acknowledgments

The authors wish to thank the members of the Medina-Acosta laboratory for their insights and productive lab brainstorming meetings.

---

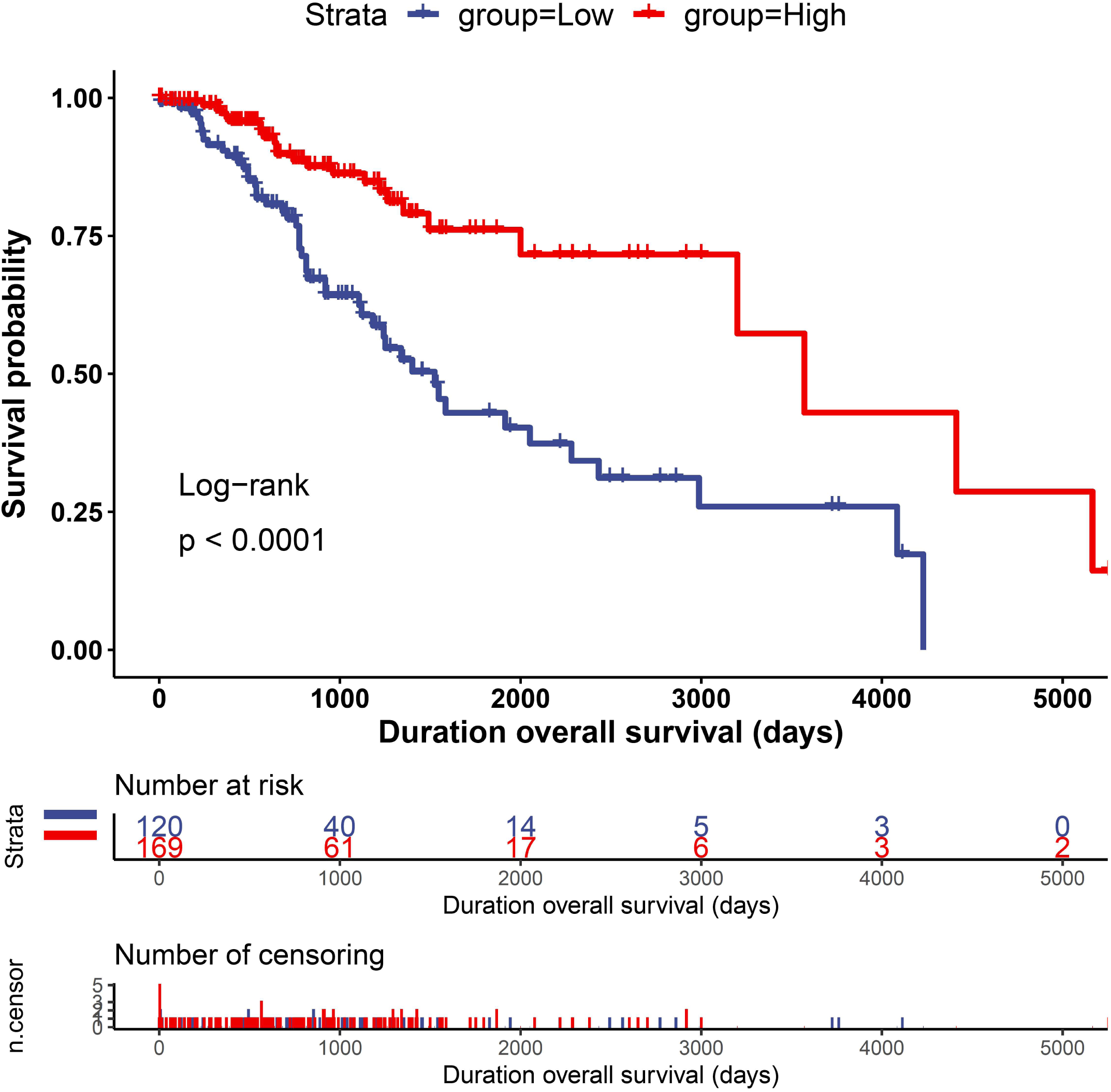

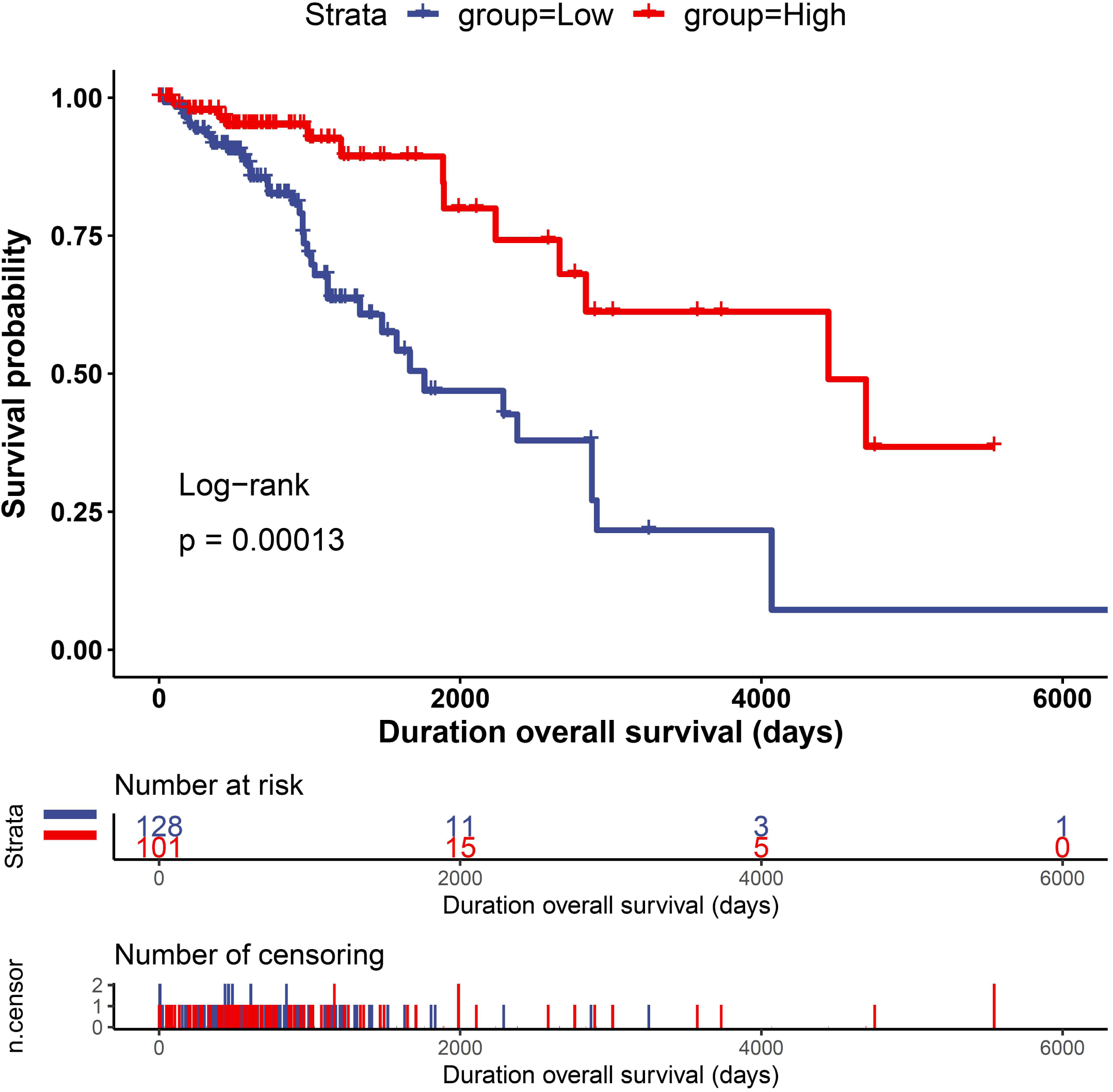

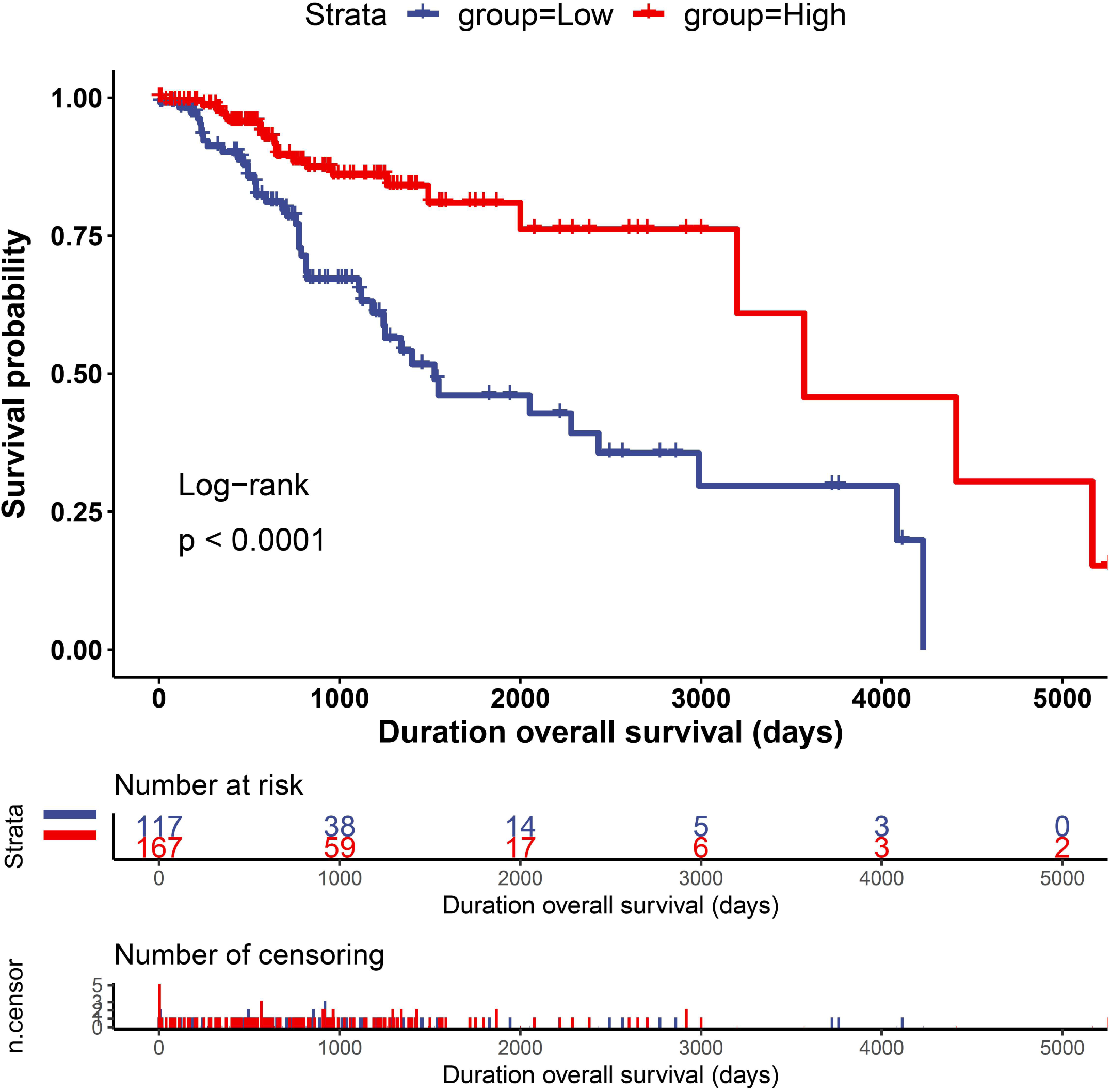

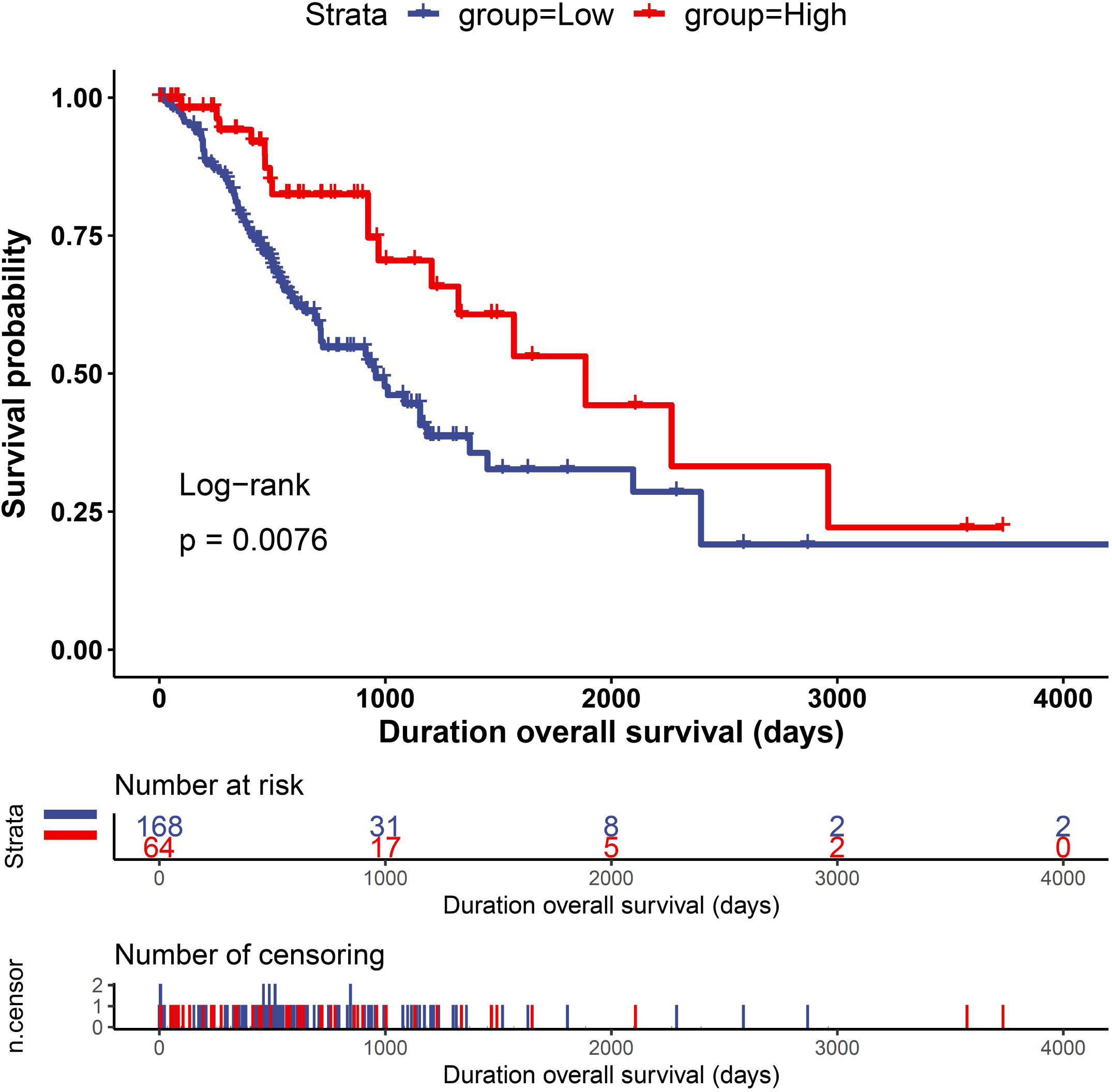

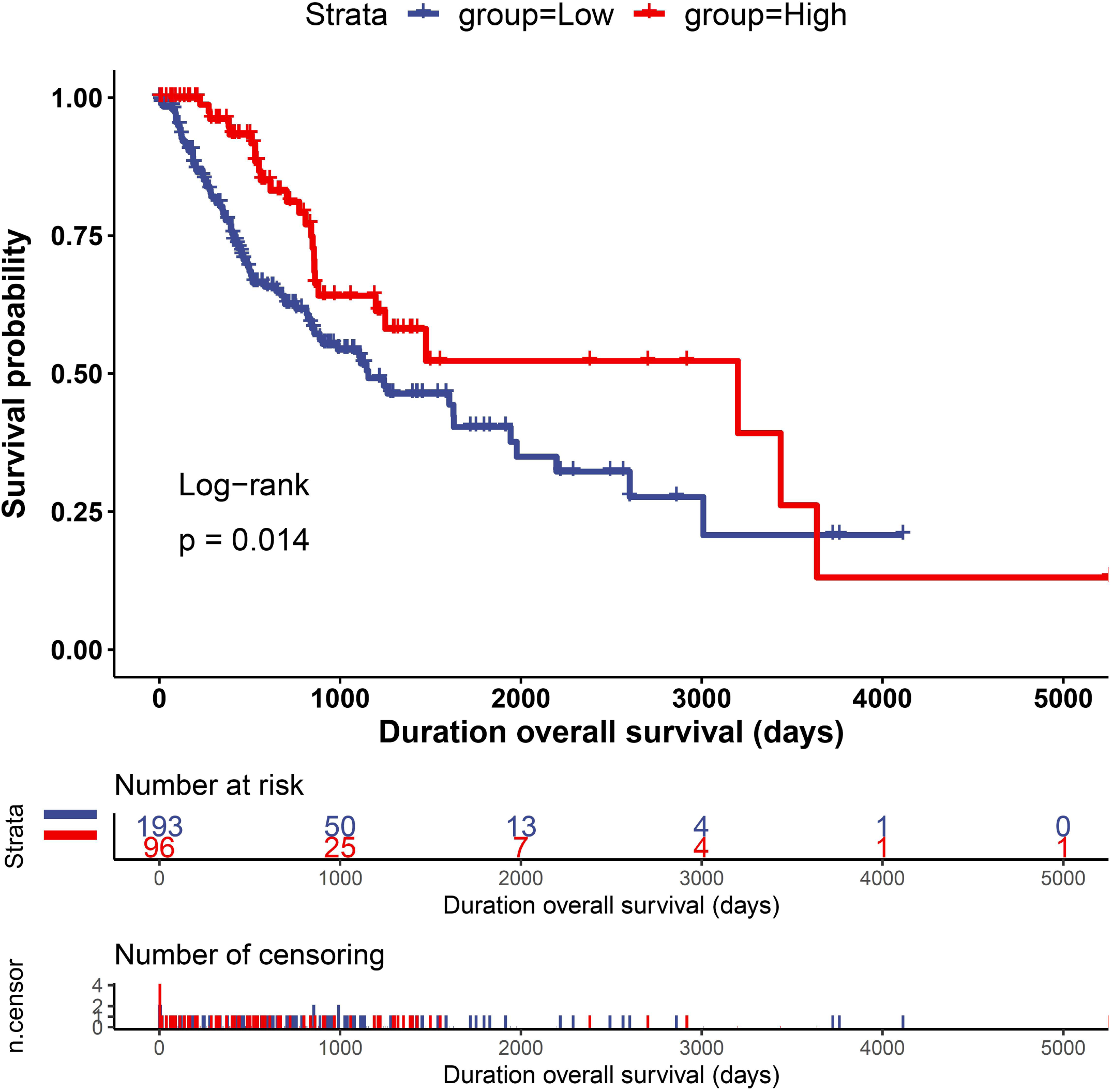

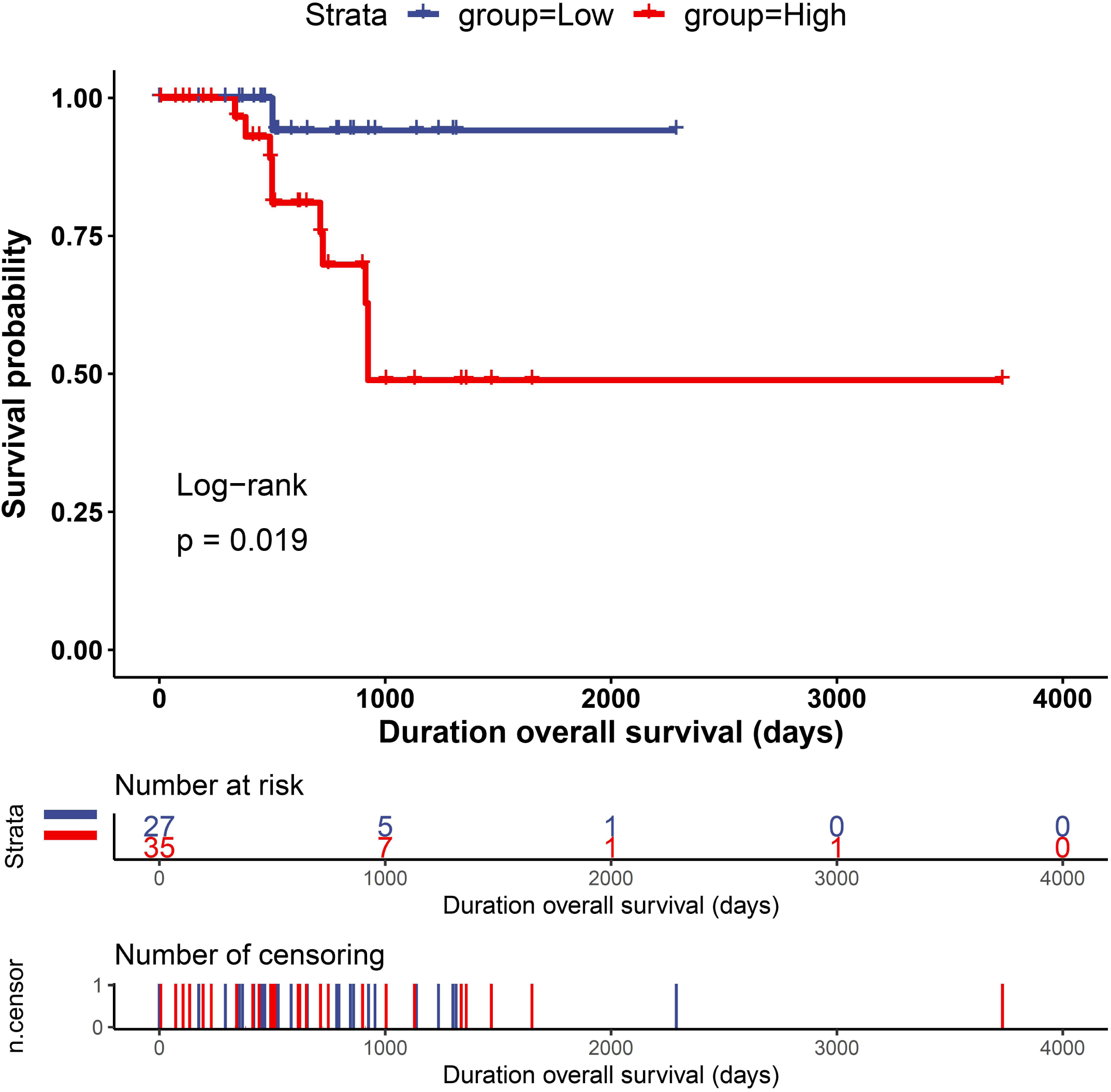

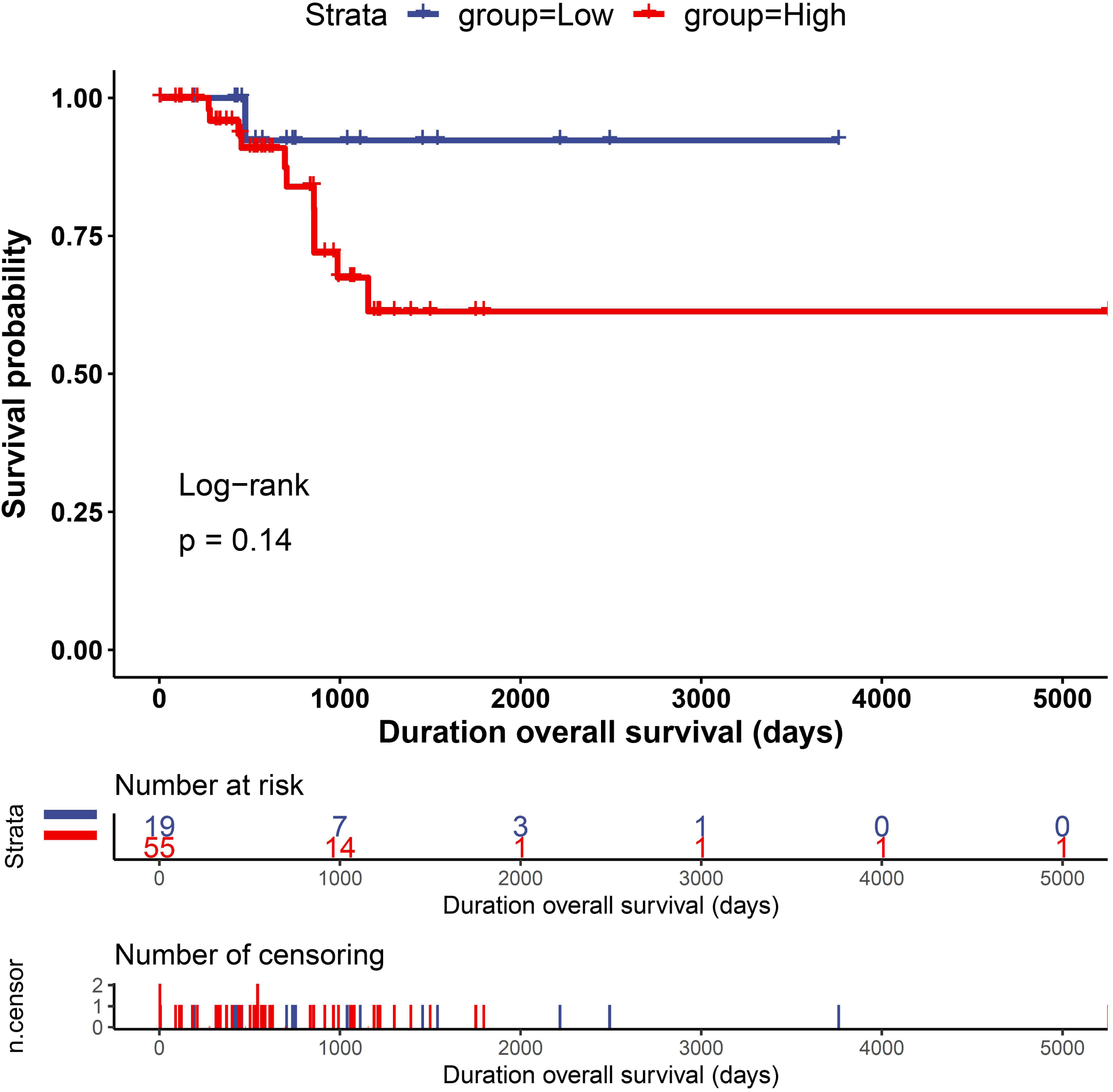

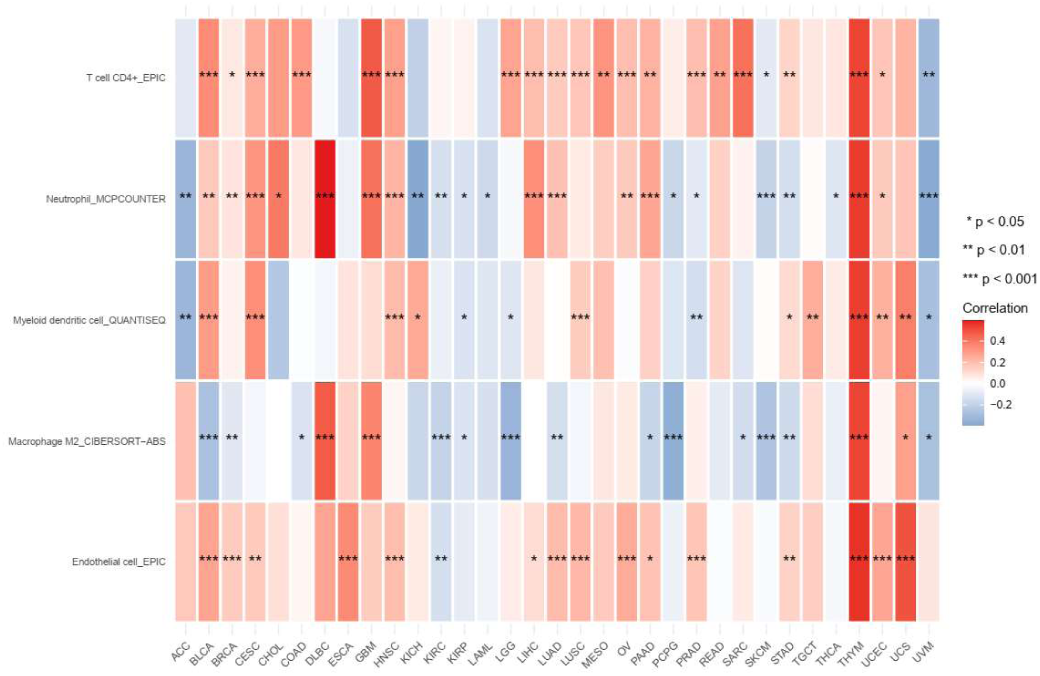

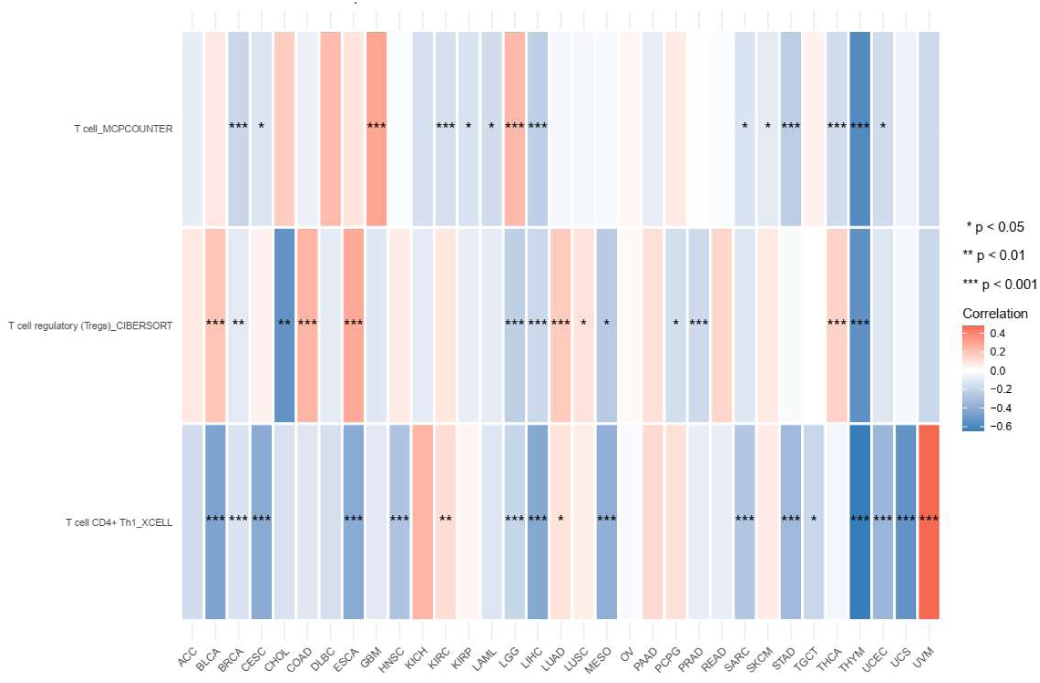

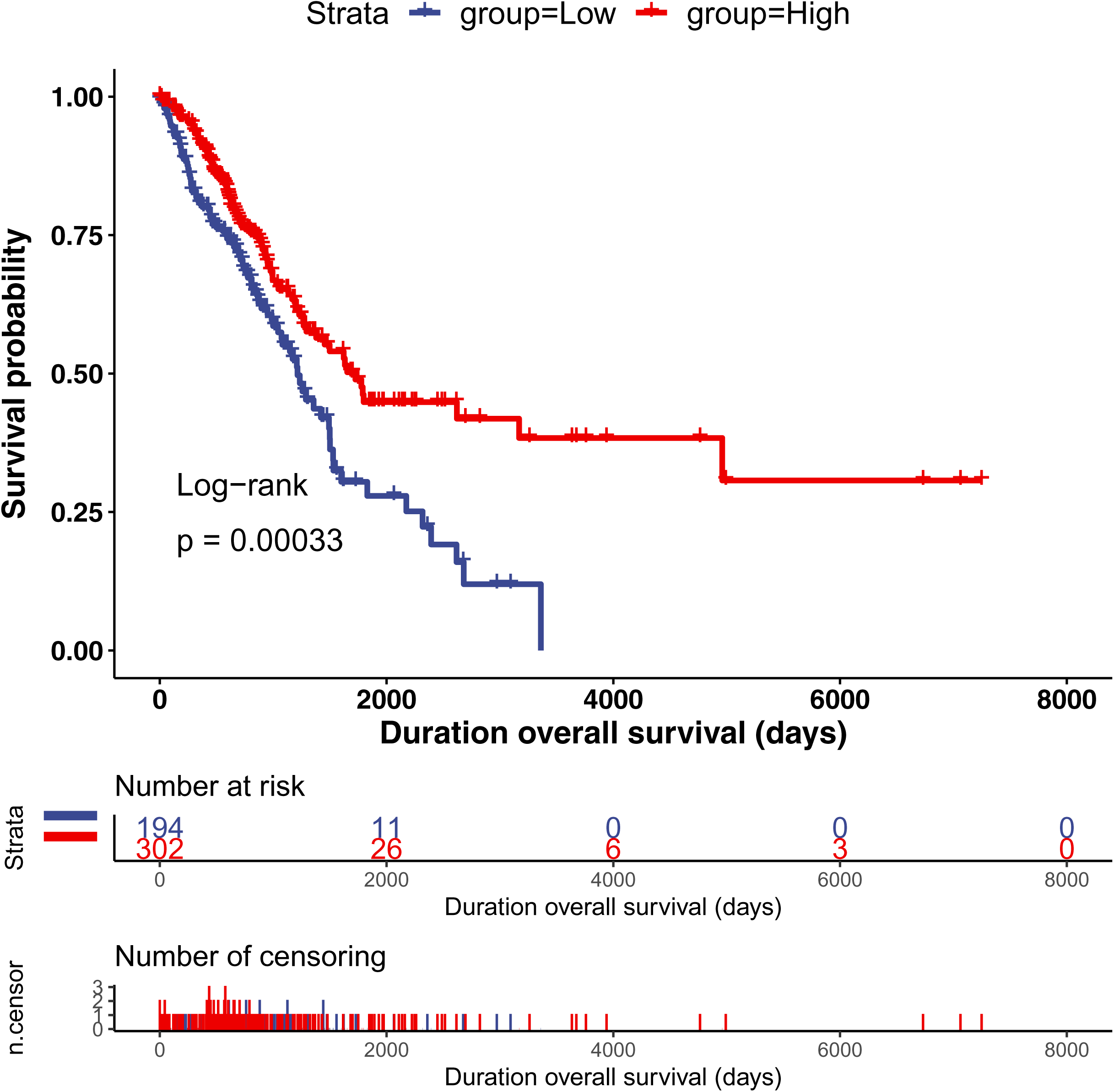

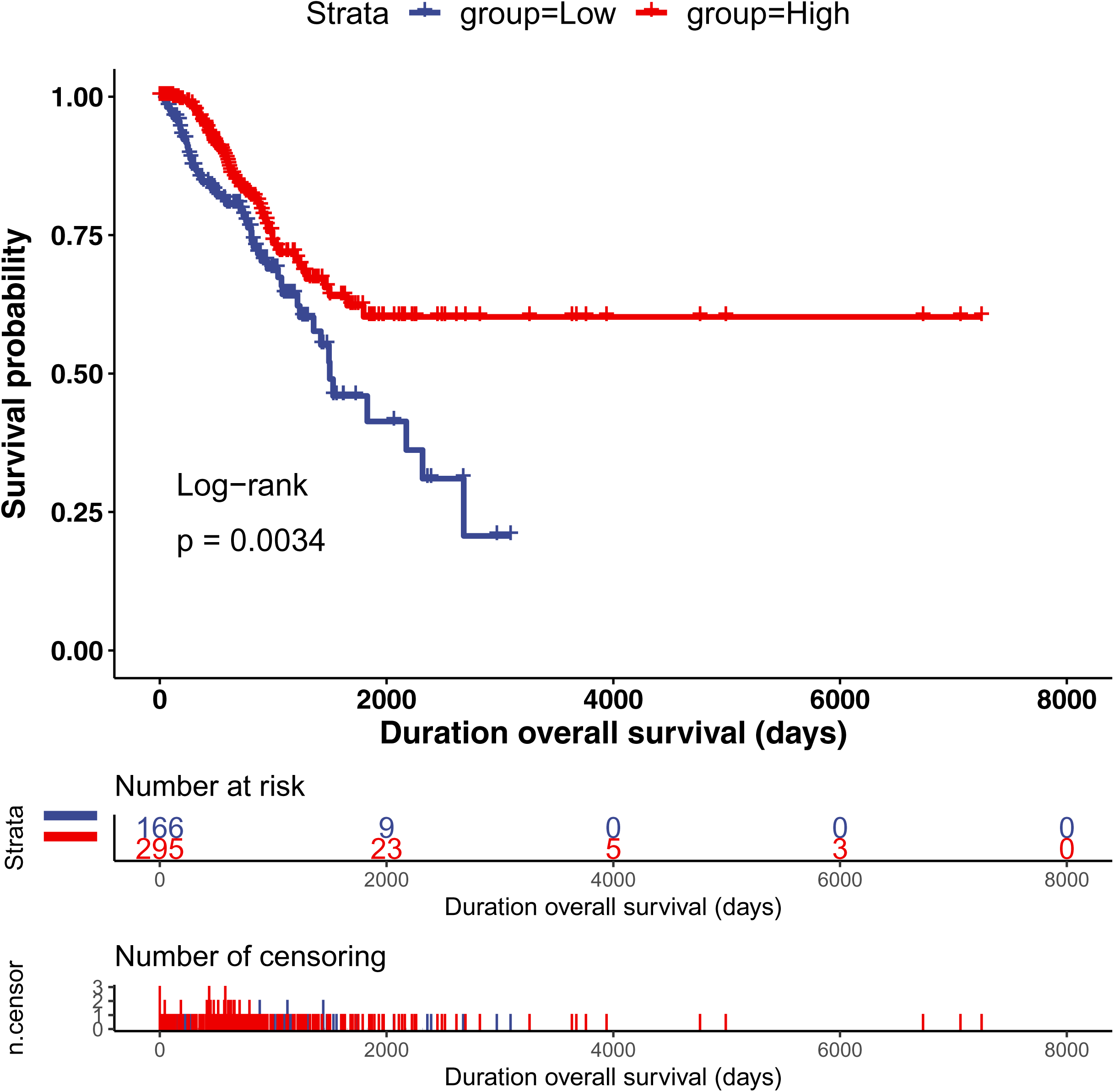

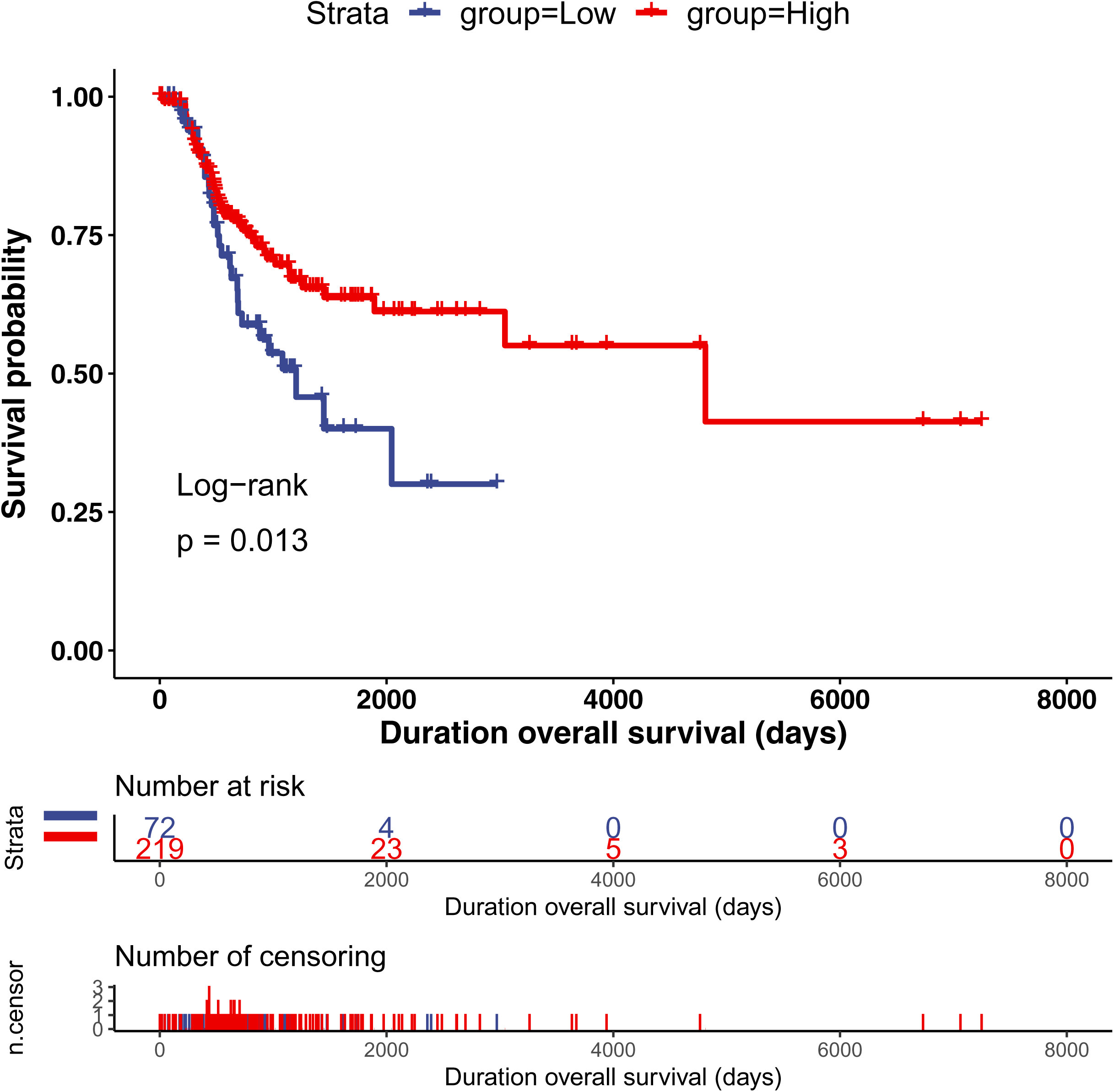

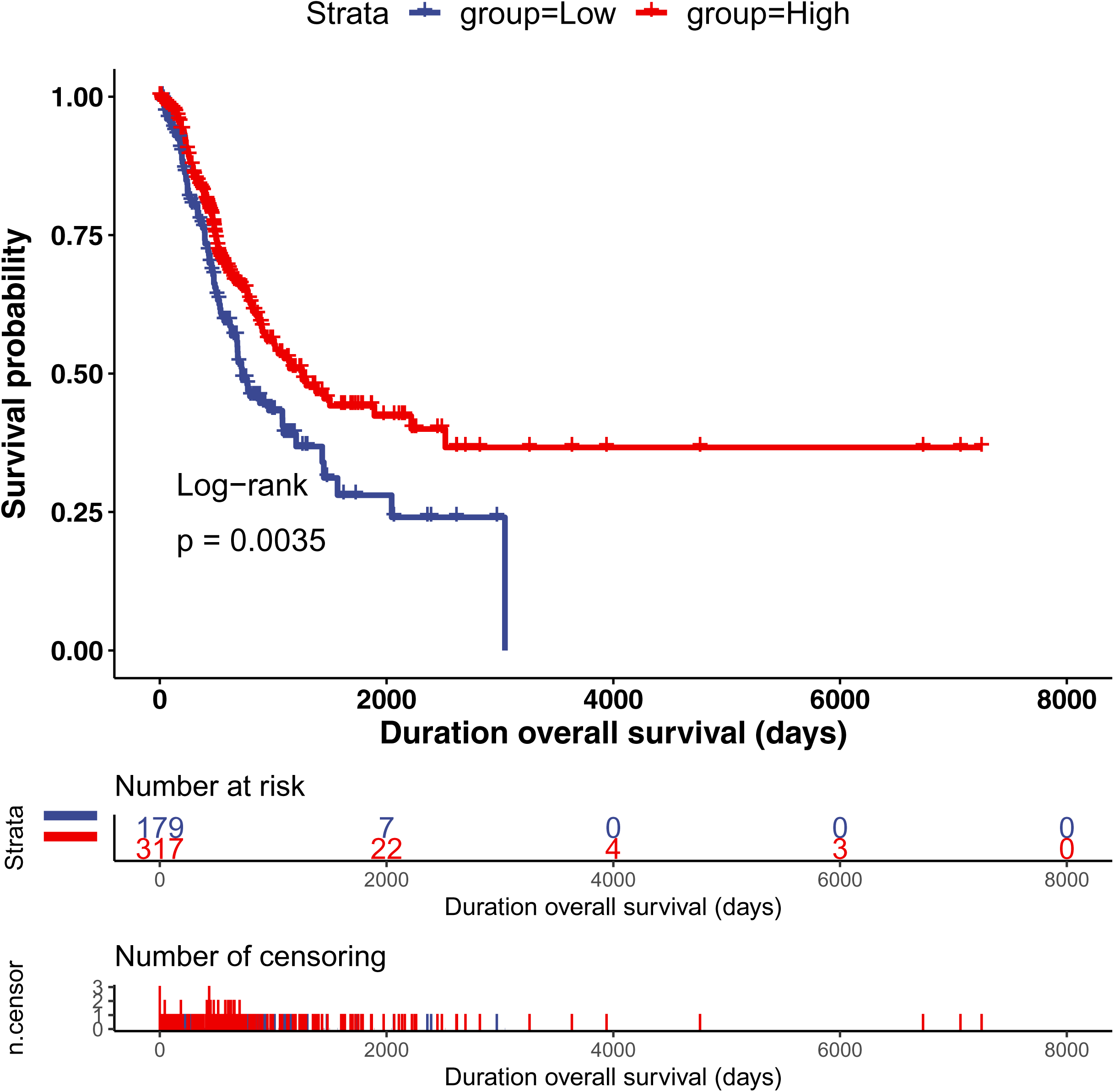

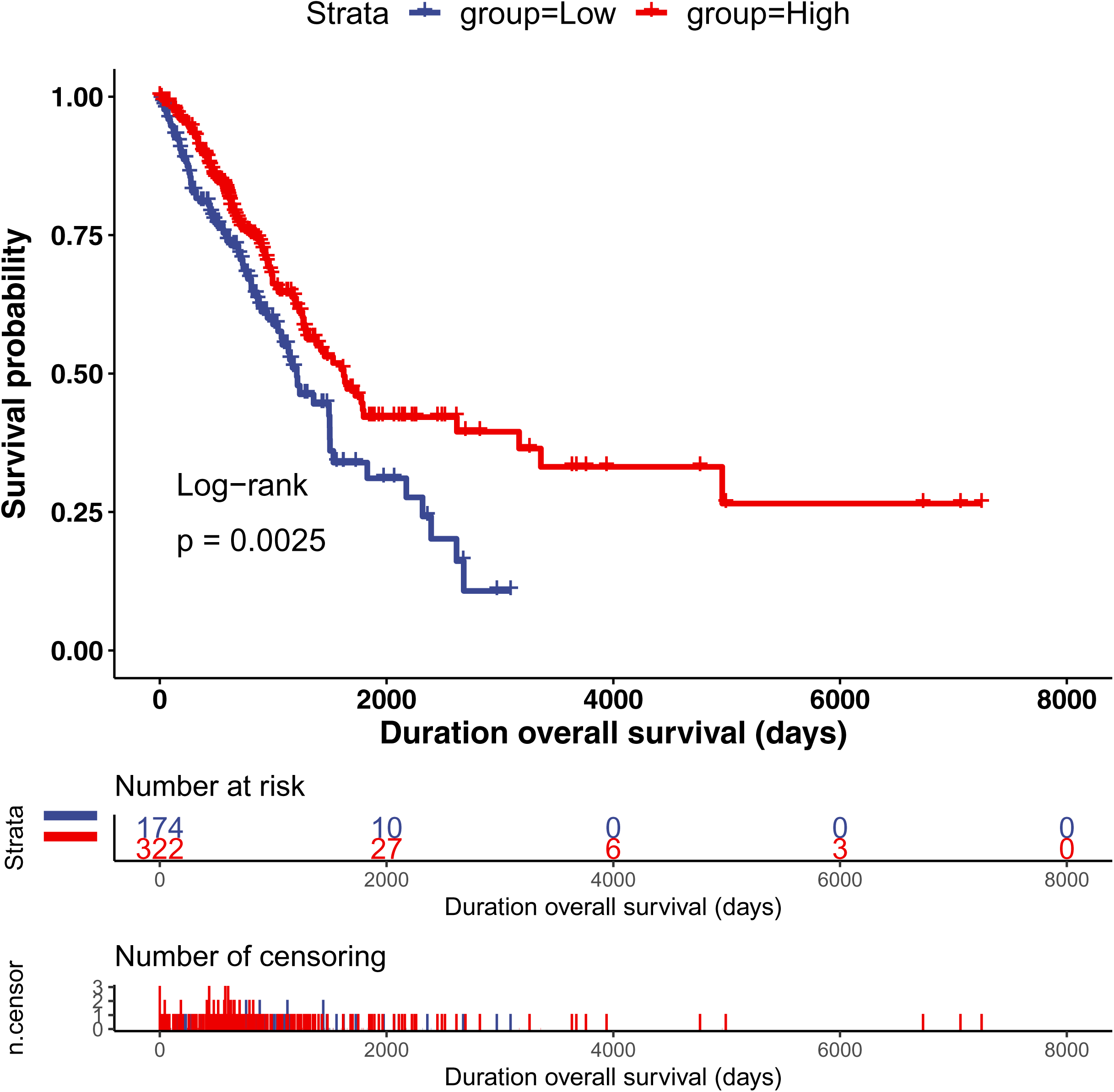

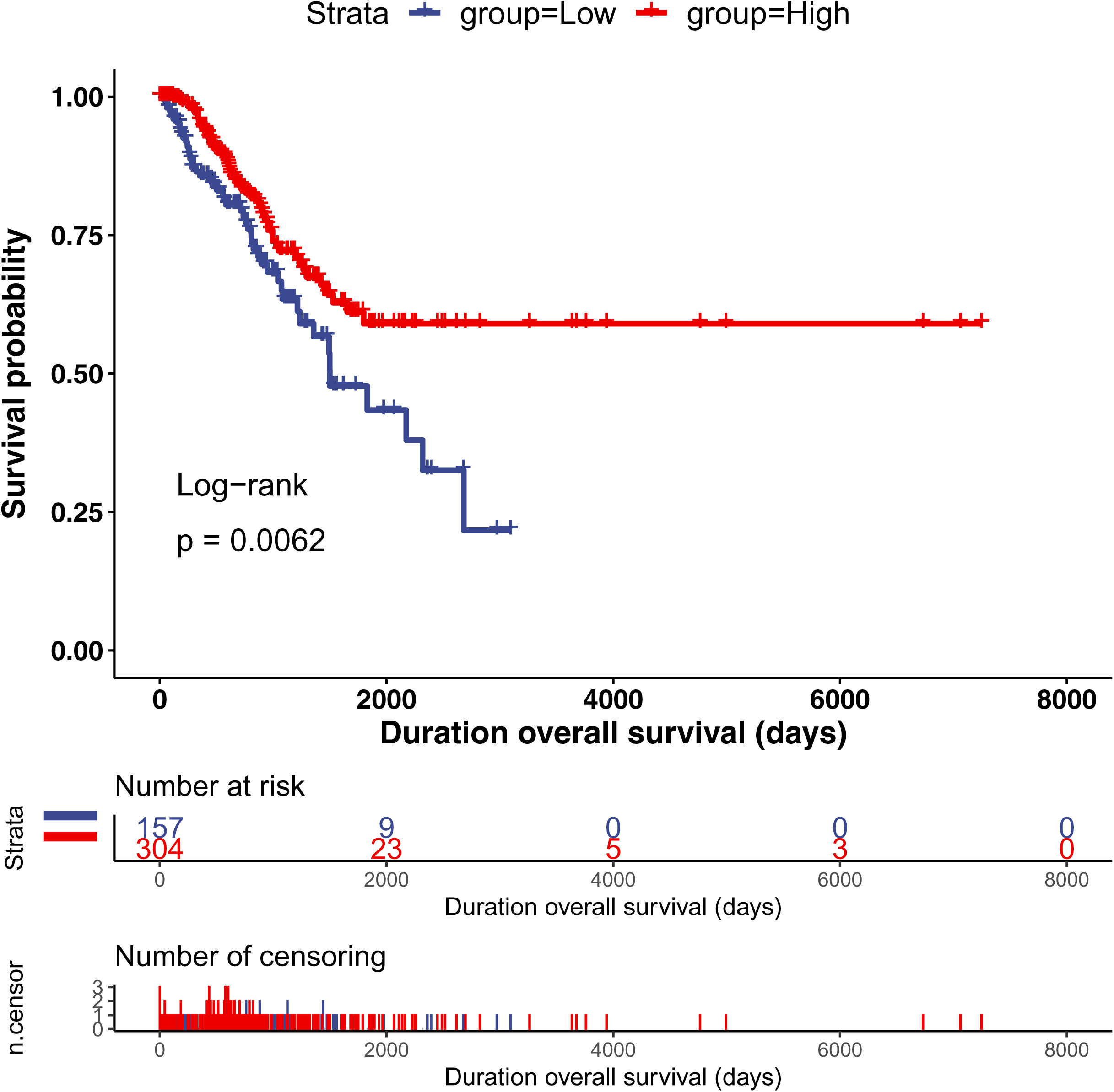

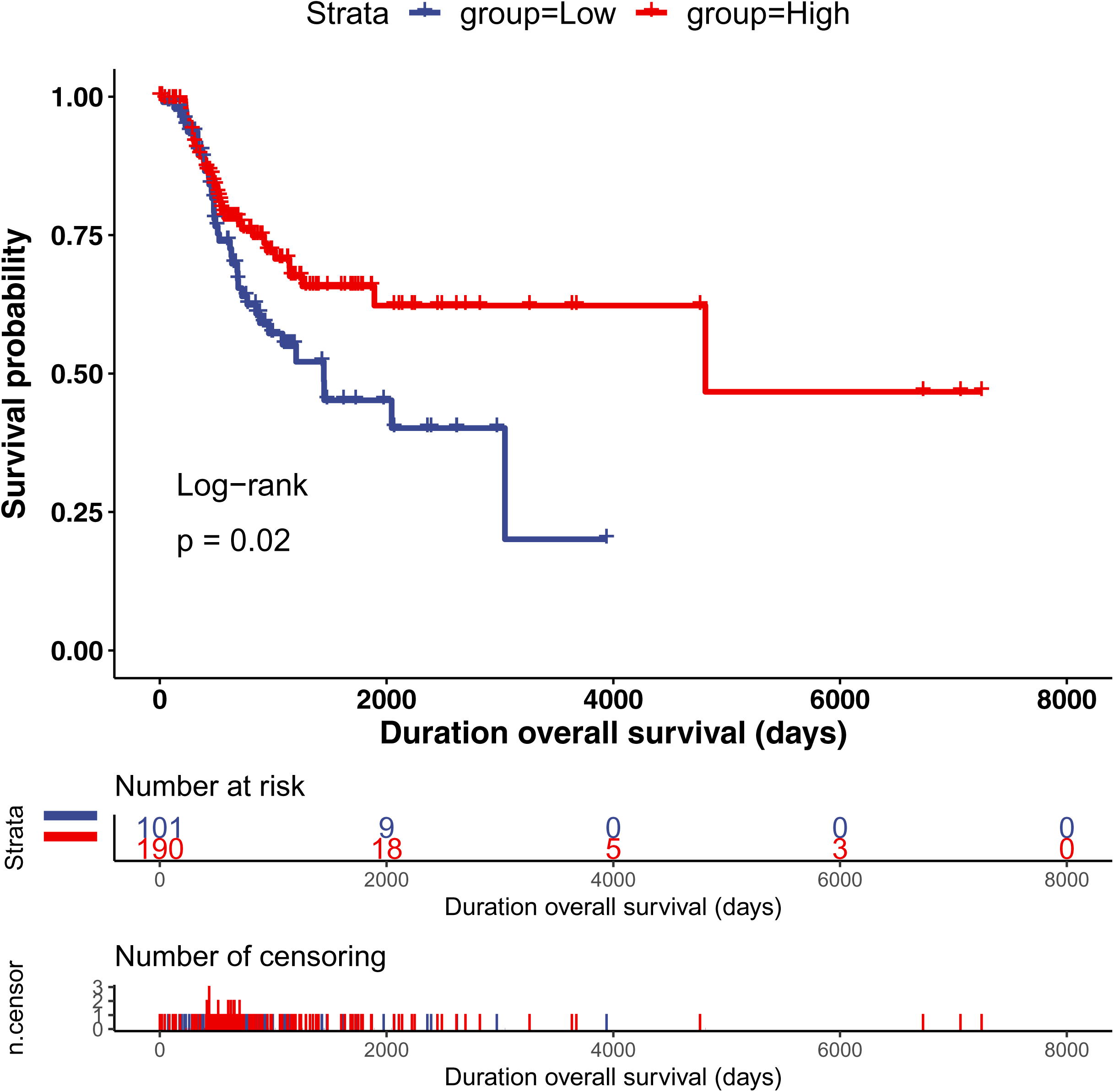

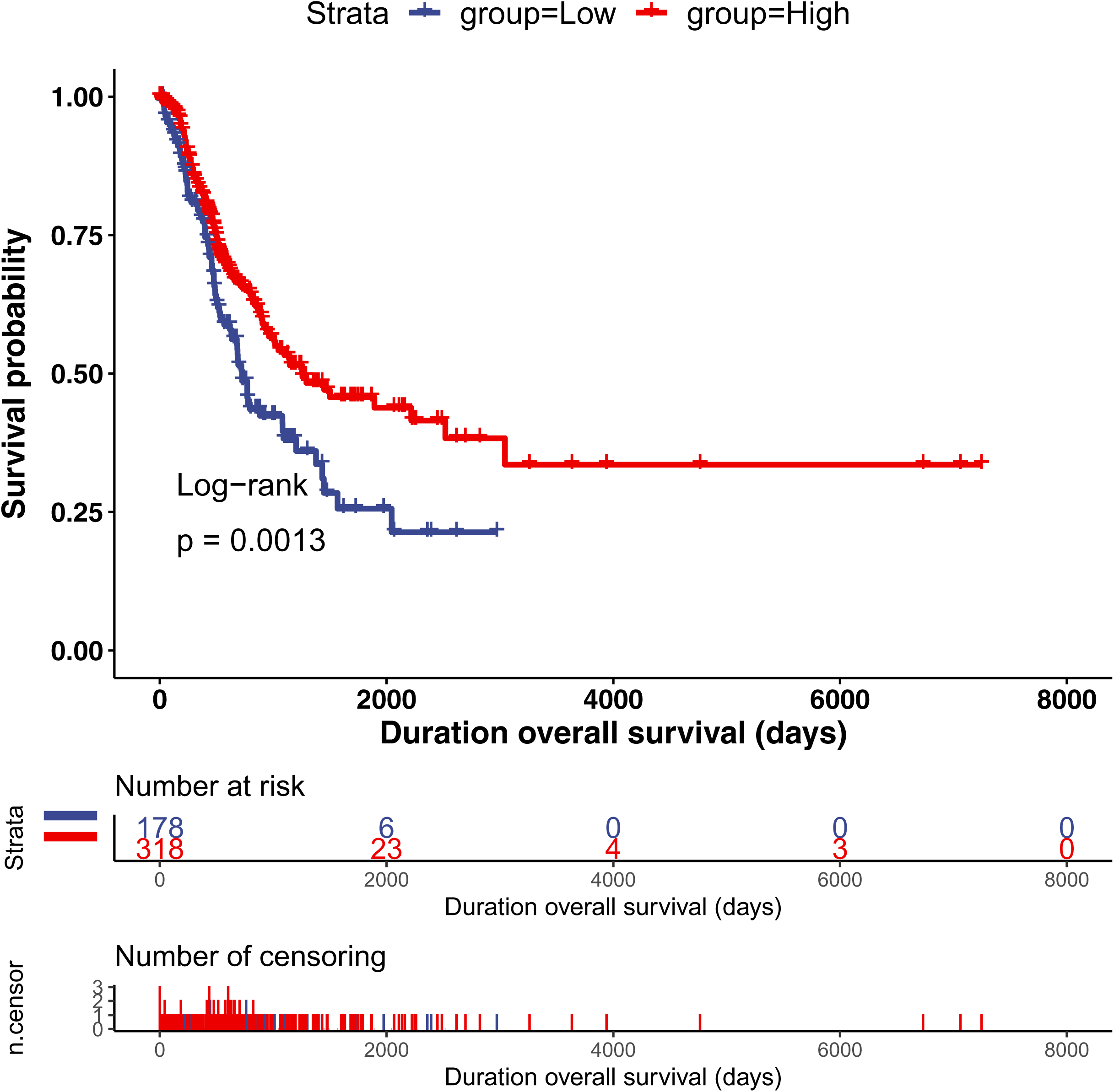

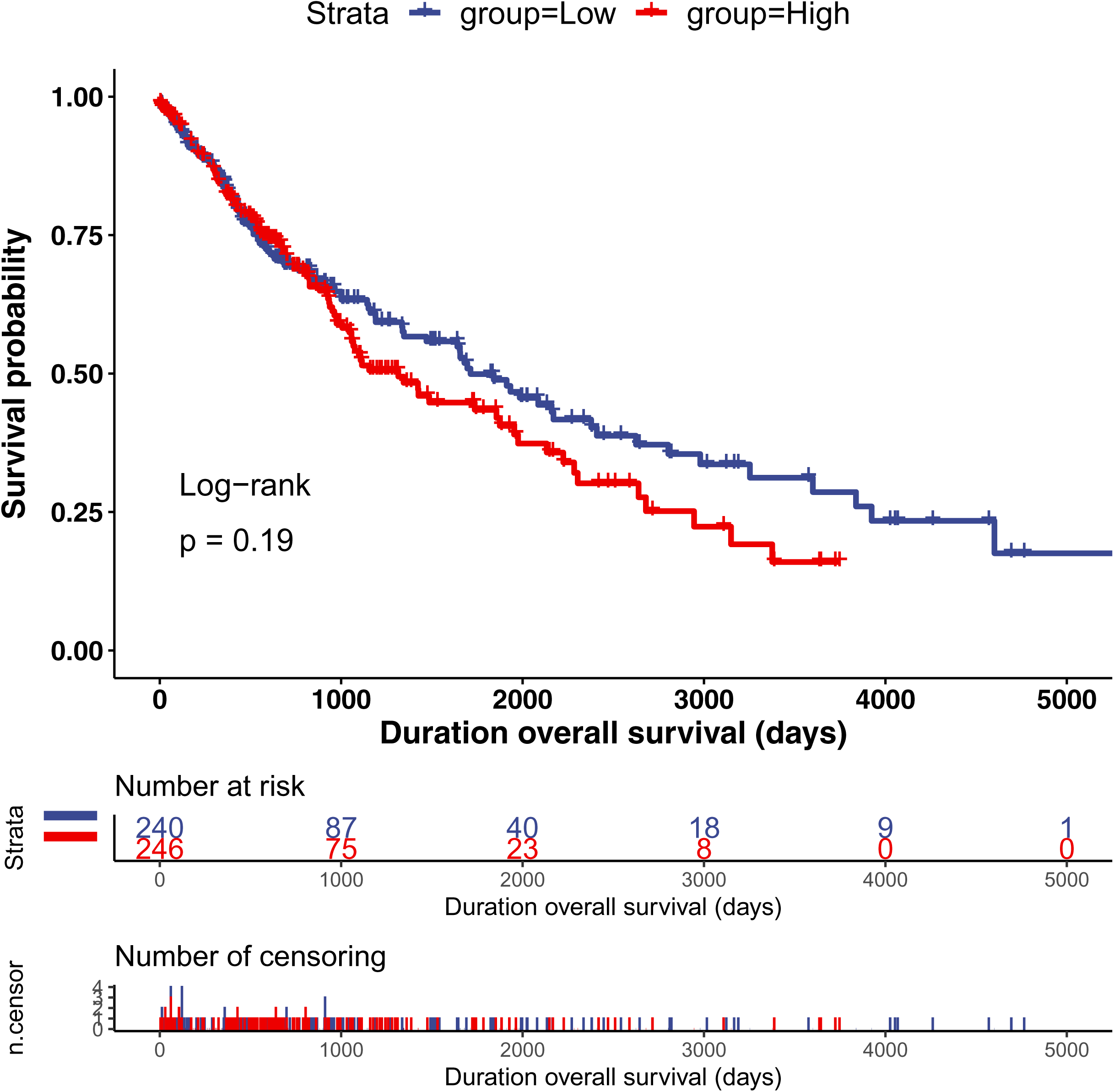

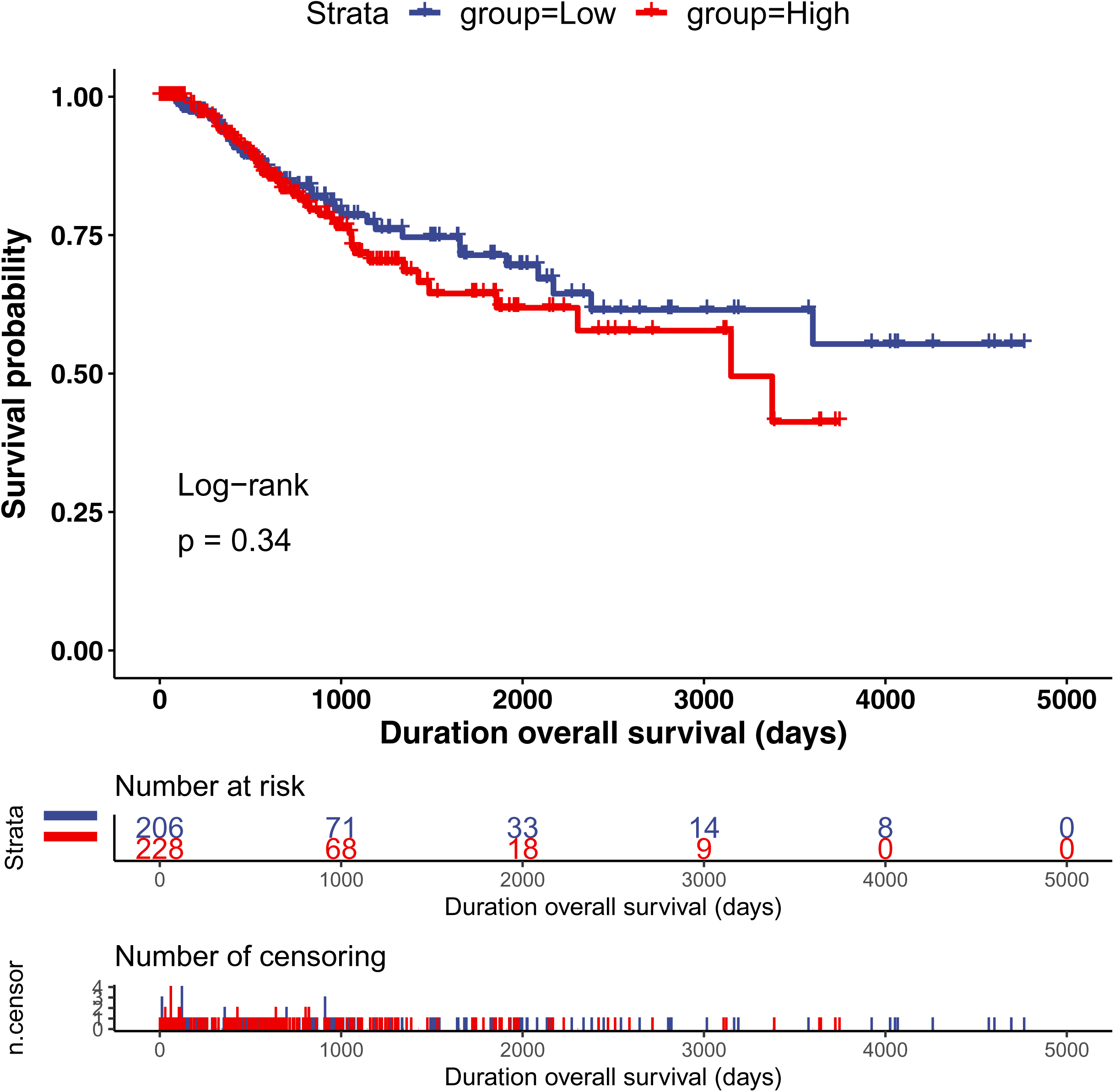

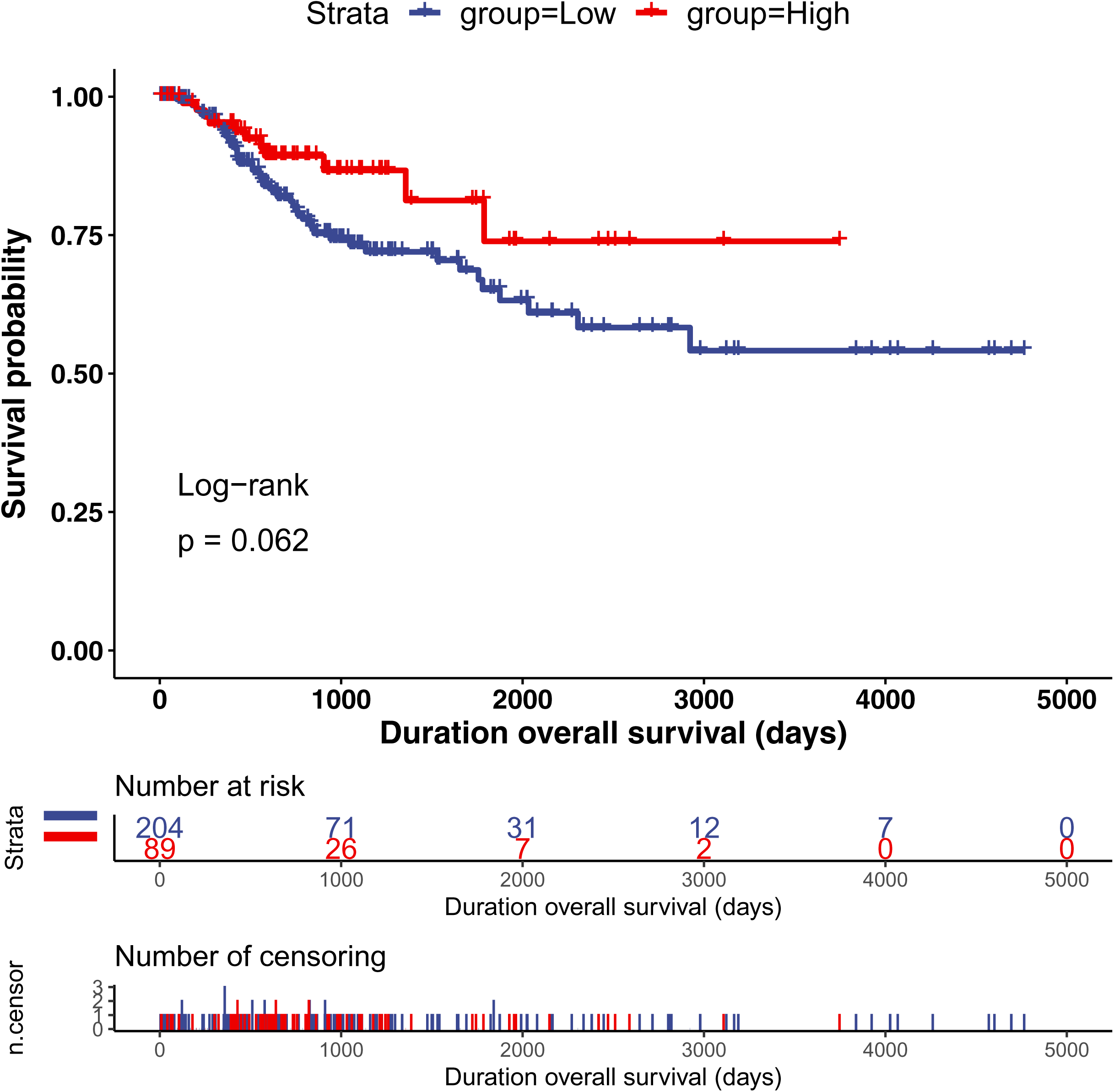

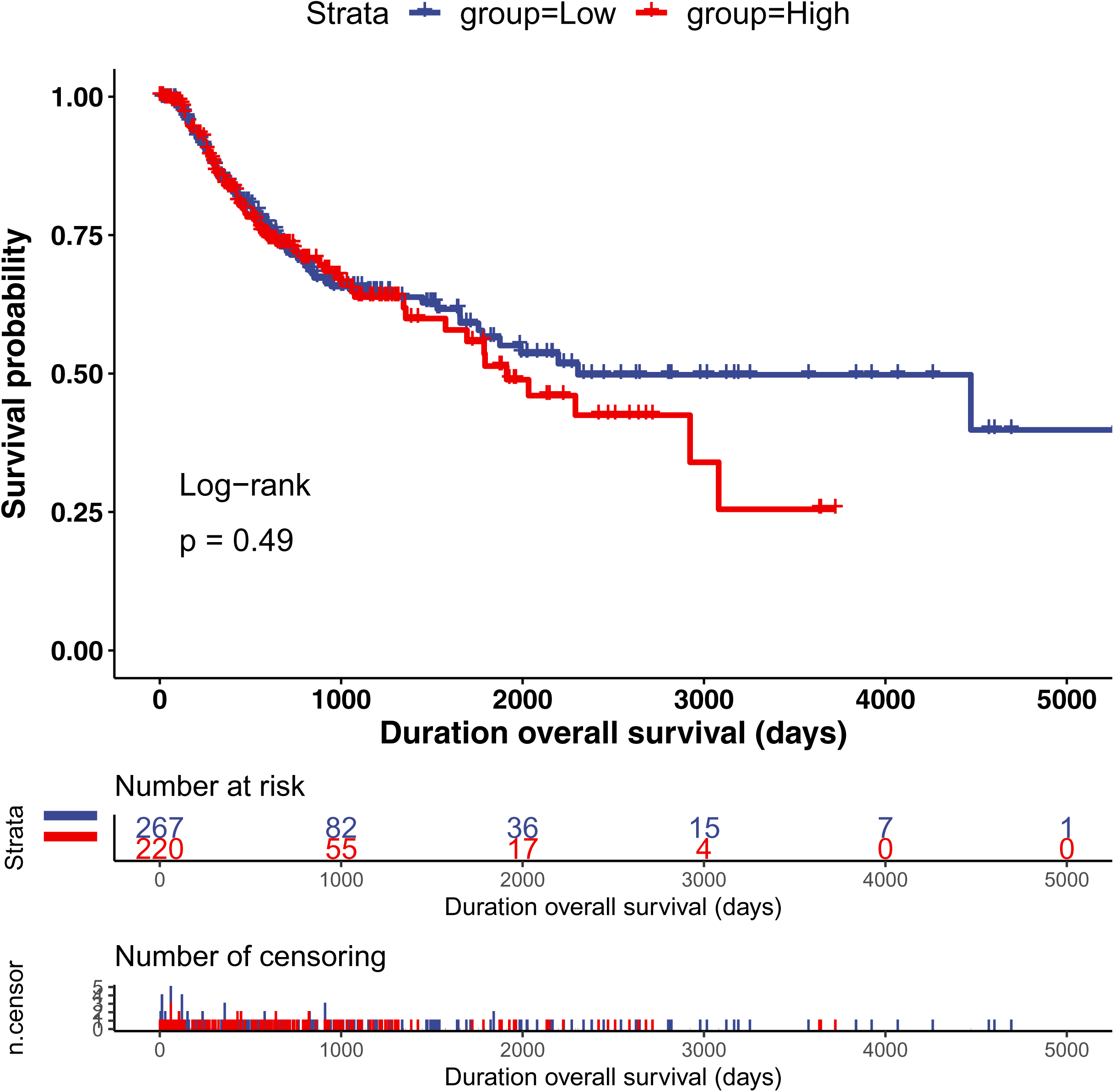

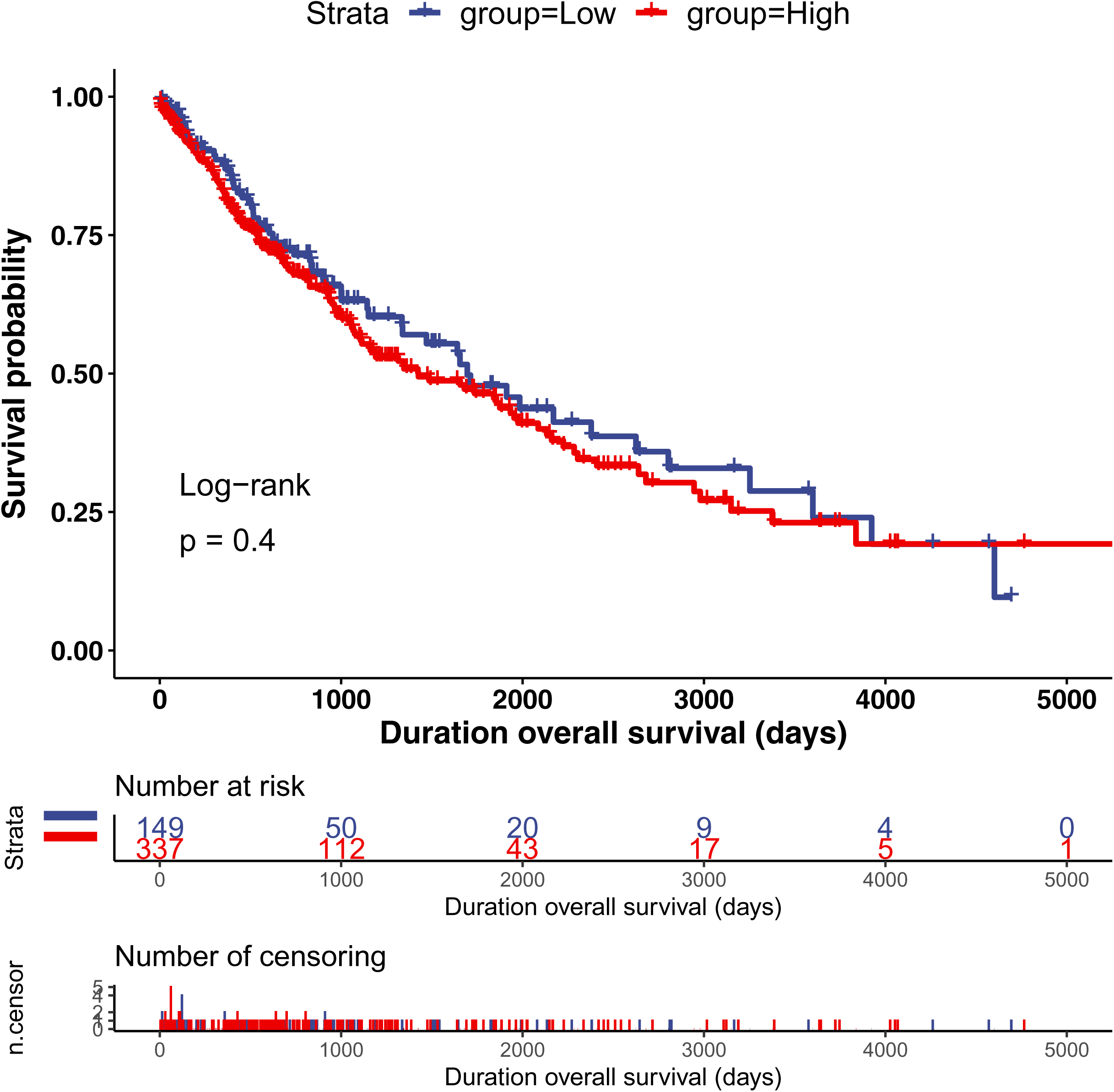

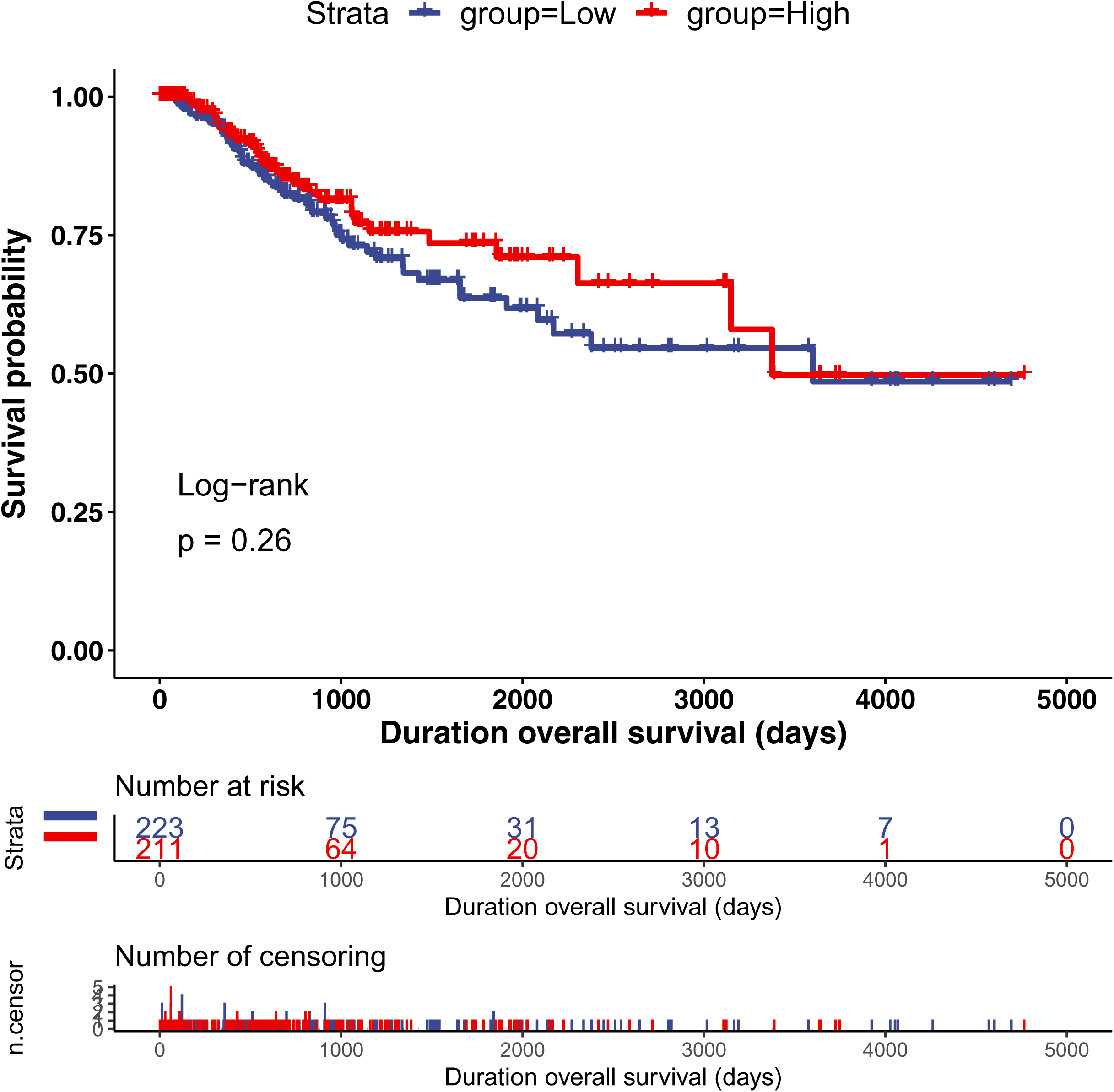

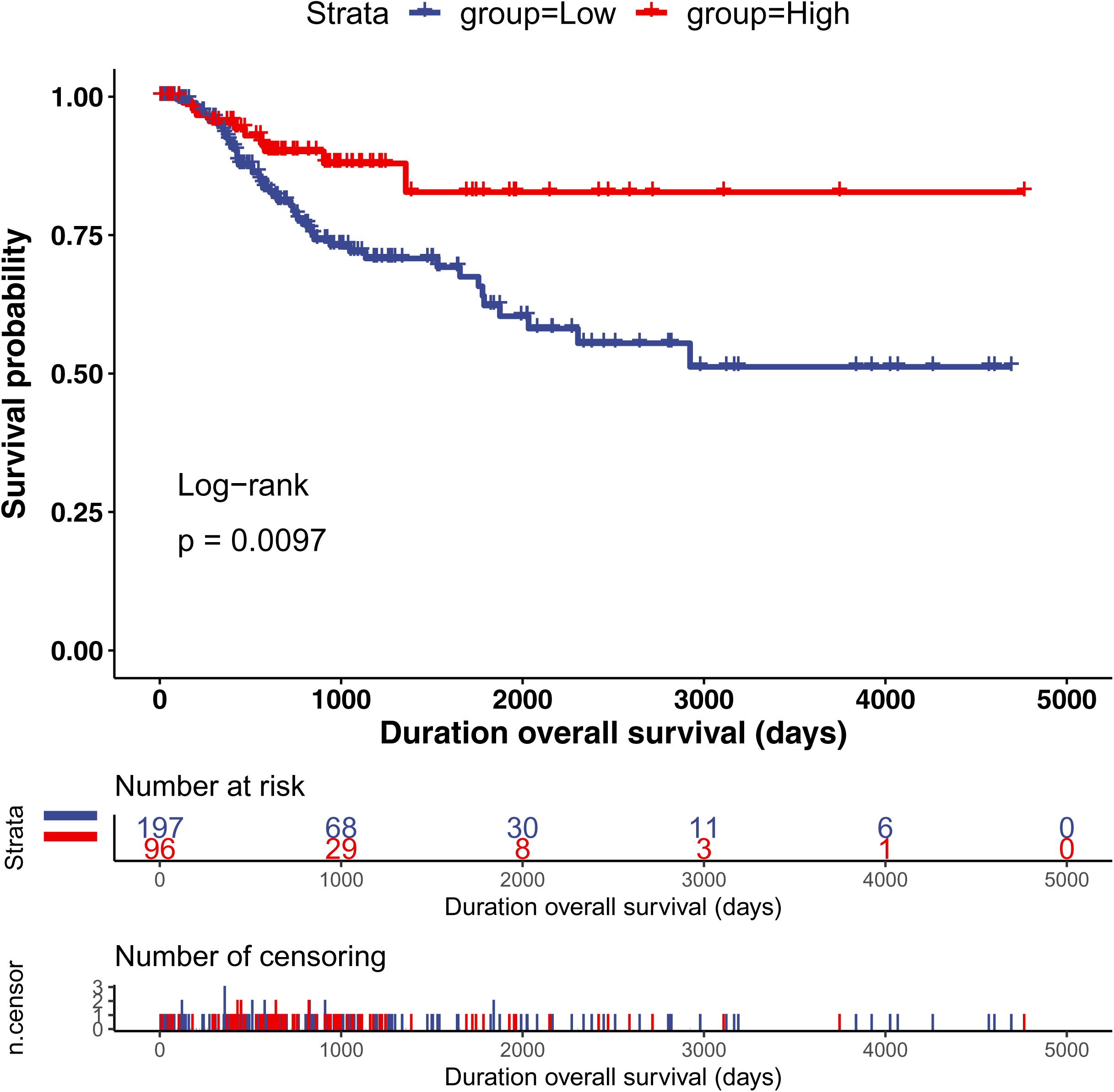

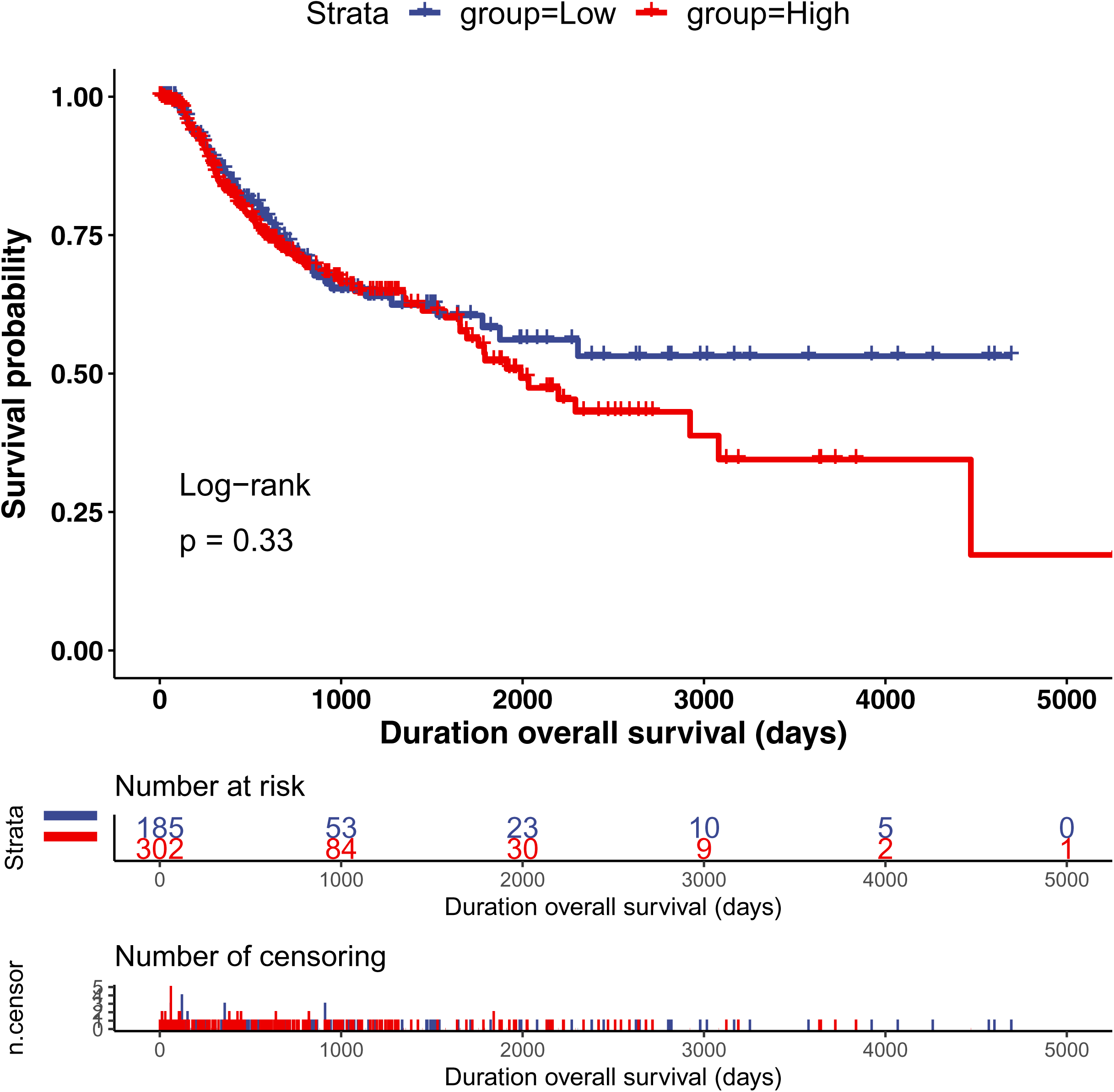

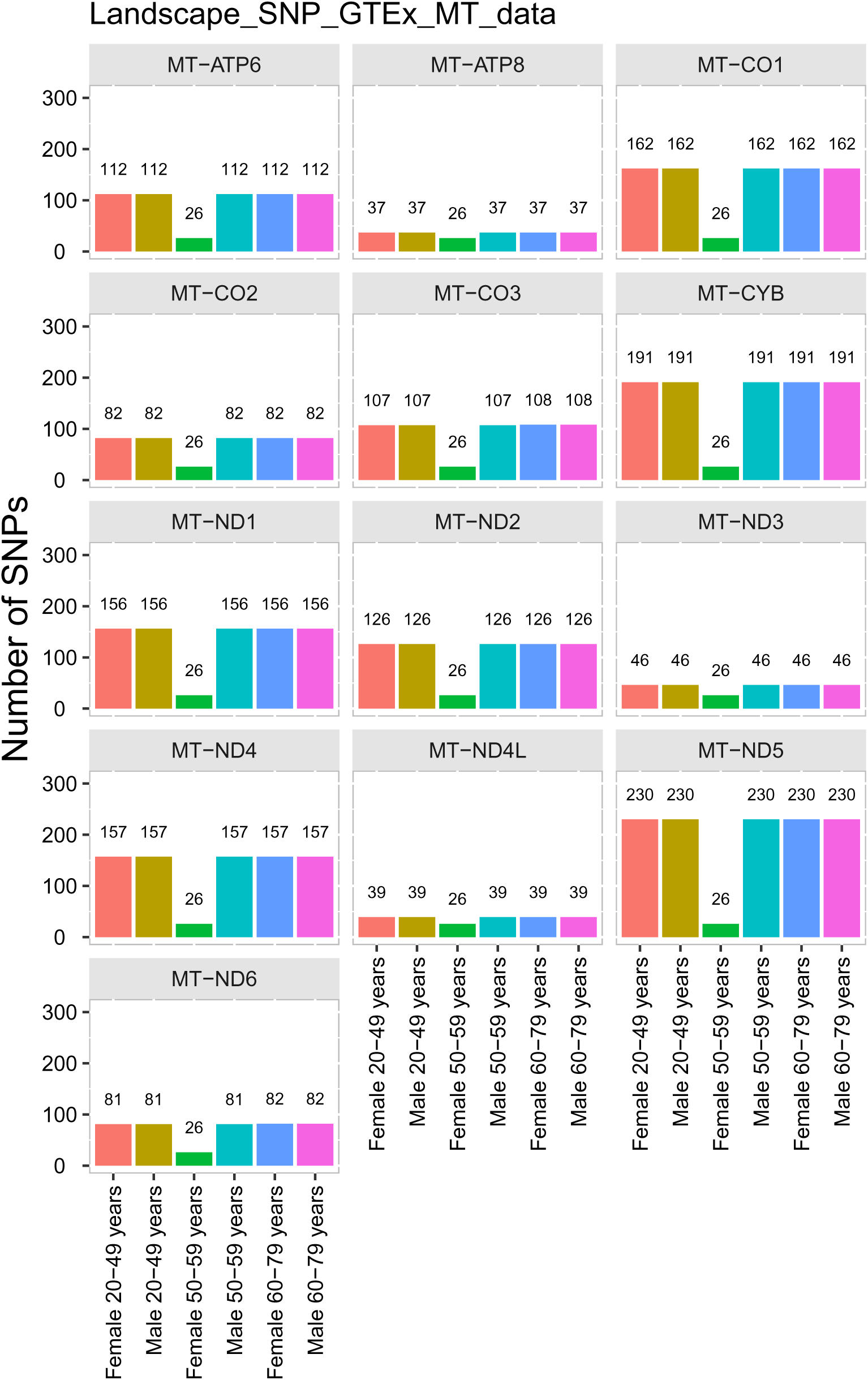

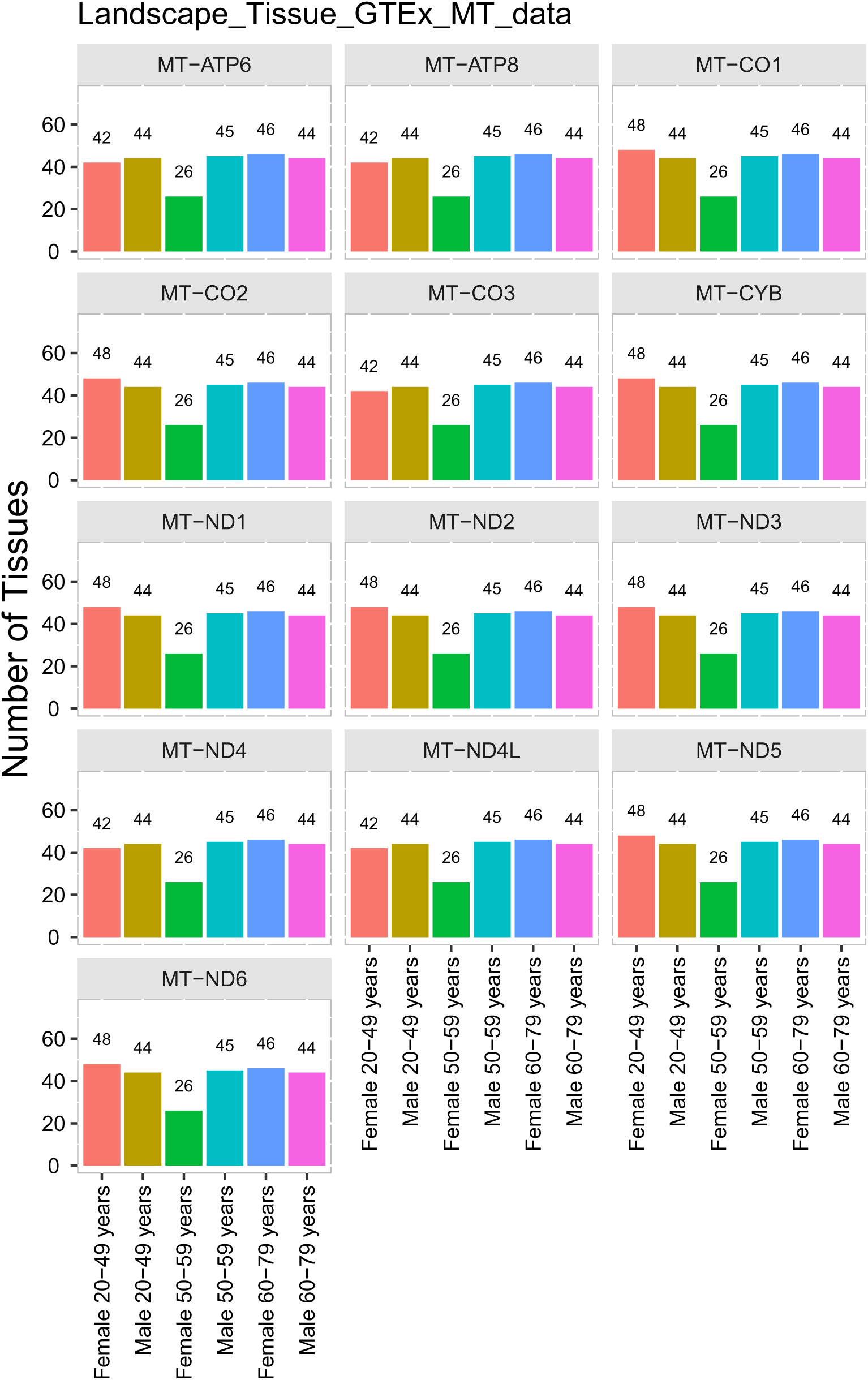

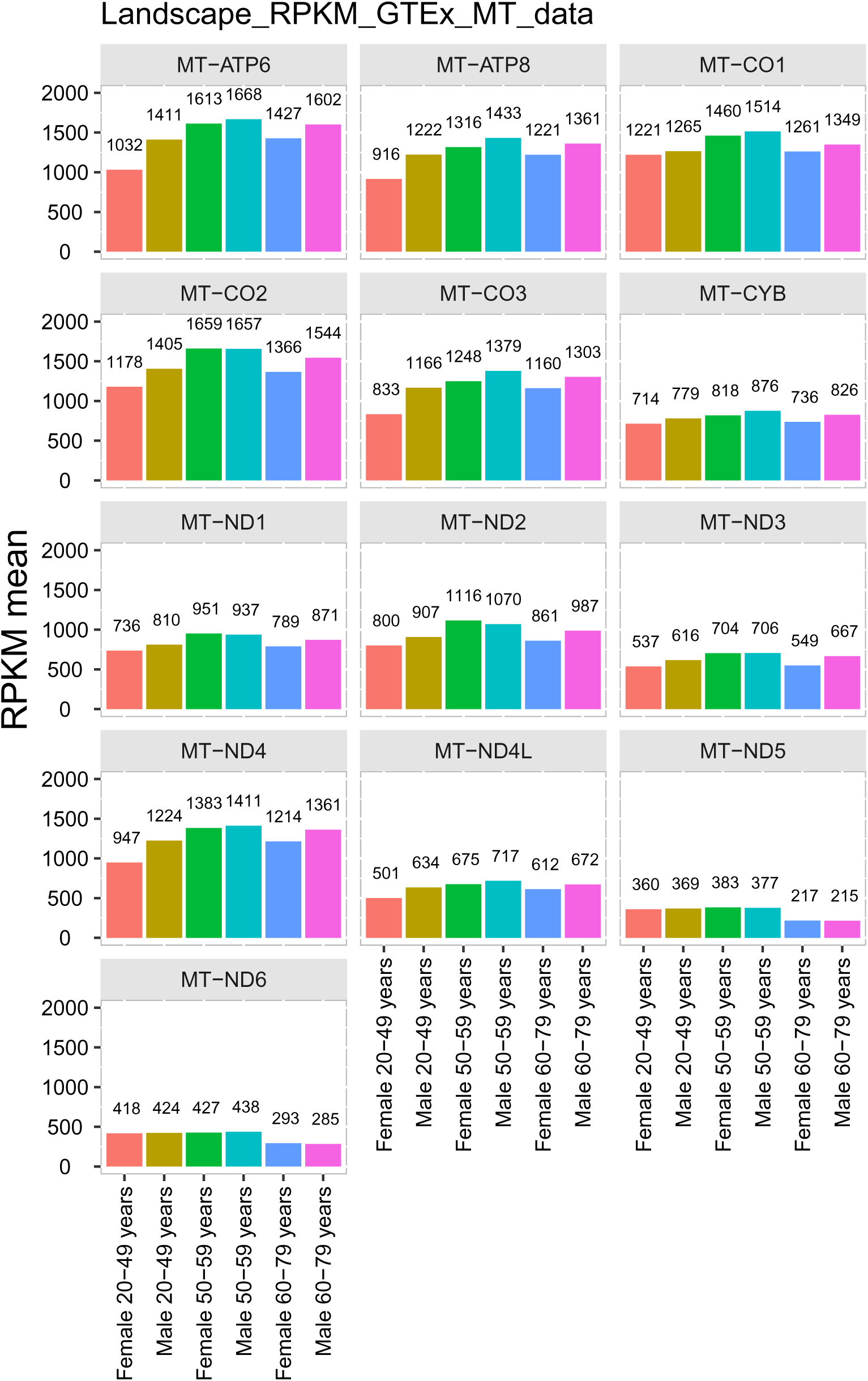

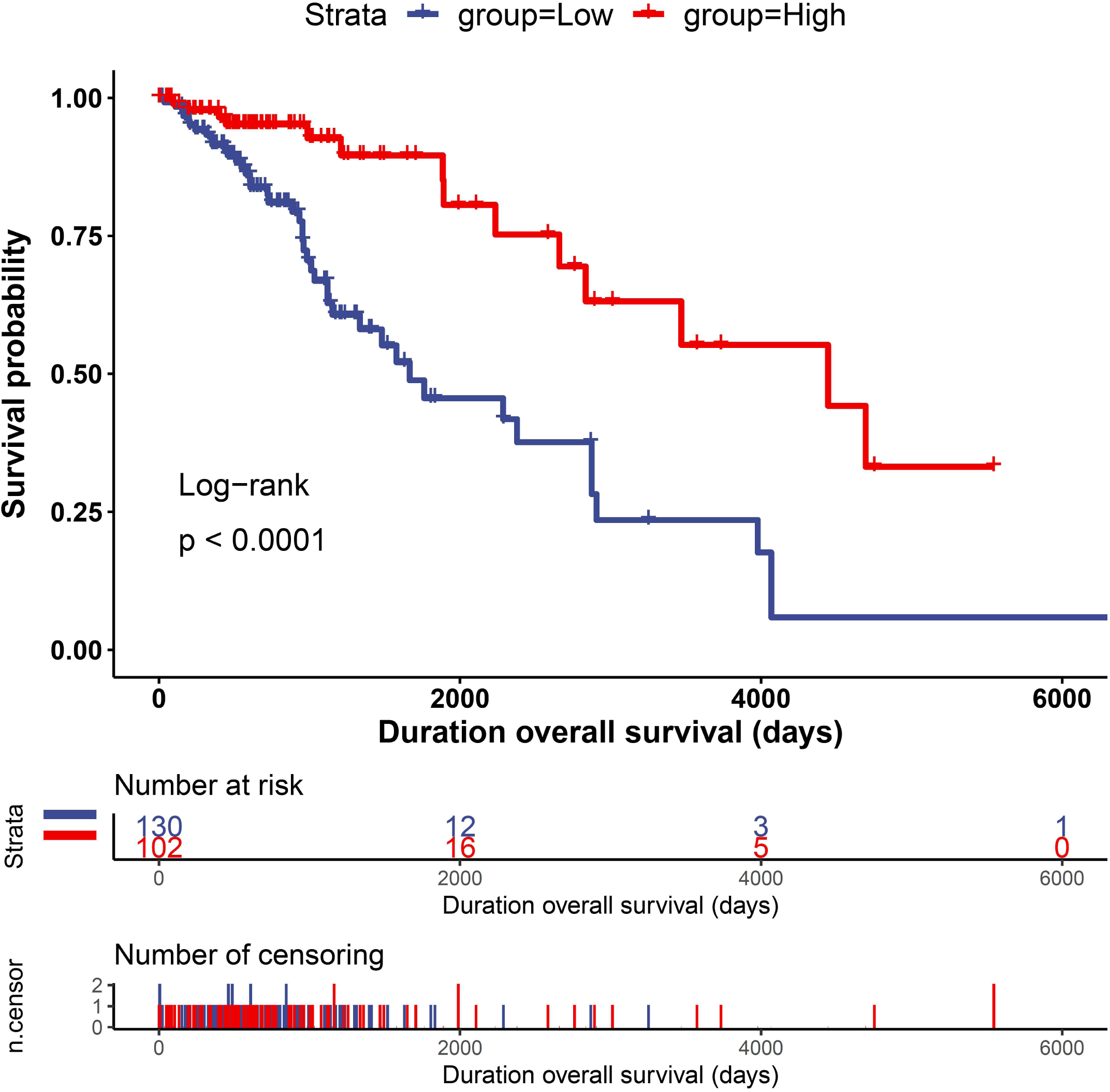

